# Spatial analysis reveals the cellular microenvironments and mechanisms of inflammation and kidney injury in acute interstitial nephritis

**DOI:** 10.1101/2025.10.29.684811

**Authors:** Megan L. Baker, Vijayakumar R. Kakade, Tifanny Budiman, Marlene Weiß, Joseph M. Cunningham, Sagar Sadarangani, Gabriel Lerner, Gilbert Moeckel, Avi Z. Rosenberg, Chirag R. Parikh, Yuval Kluger, Dennis G. Moledina, Lloyd G. Cantley

## Abstract

Acute interstitial nephritis (AIN) causes 15-20% of all acute kidney injury cases but lacks effective therapies beyond corticosteroids. Using high-resolution imaging mass cytometry and single-cell spatial transcriptomics to analyze human kidney biopsies with AIN, non-immunologic acute tubular injury (ATI), and reference tissue, the CXCL9-CXCR3 axis was identified as the defining immunologic signature of AIN, with 44-fold higher predicted CXCL9-CXCR3 interactions than ATI, creating homotypic inflammatory T cell amplification networks concentrated in lymphoid aggregates. *C3AR1*^+^ immune cells were enriched in peritubular neighborhoods of complement 3-expressing injured tubule cells, predominantly *VCAM1*^+^ injured proximal tubules, linking tubular injury to immune activation in AIN. Nicotinamide phosphoribosyltransferase (*NAMPT*) was the strongest predictor of *VCAM1*^+^ tubular microenvironments, with expression by both injured tubules and surrounding immune cells coordinating metabolic-inflammatory niches. These findings reveal distinct molecular circuits underlying AIN pathogenesis and identify potential therapeutic targets for improving clinical management and preventing progression to chronic kidney disease.

---

Acute kidney injury (AKI) affects 10-15% of hospitalized patients and up to 50% of critically ill patients, with mortality reaching 50% in severe cases requiring dialysis [1–4]. Among the various causes of AKI, acute interstitial nephritis (AIN) is present in 15-20% of biopsy-proven cases and carries significant morbidity, with high rates of progression to chronic kidney disease (CKD) despite treatment [5–10]. The clinical burden of AIN is increasing, particularly due to the broadening range of medications identified as triggers, most notably immune checkpoint inhibitors [11–13]. Treatment remains limited to discontinuing the offending agent and empirical corticosteroid therapy, which fails in 20-40% of patients [14, 15]. Despite well-characterized histopathology [12, 13], the molecular circuits underlying AIN inflammation remain undefined, limiting targeted therapy development. The therapeutic challenge is compounded by the narrow 7–10-day window for steroid efficacy [10] and significant steroid-associated complications, particularly in cancer patients receiving immune checkpoint inhibitors.

While the immune cell infiltrate in AIN has been carefully described [7, 12, 16–18], understanding AIN pathogenesis and outcomes requires detailed characterization of the bidirectional interactions between immune cells and tubular epithelium. Upon injury, proximal tubule (PT) cells express KIM-1 (kidney injury molecule-1, encoded by *HAVCR1*) and can upregulate VCAM1 (vascular cell adhesion molecule-1). VCAM1 mediates cellular interactions with VLA-4^+^ leukocytes and marks a failed-repair tubular state that predominates in CKD [19–22]. Elucidating how distinct tubular injury states participate in the orchestration of immune responses—and how these immune responses perpetuate tubular injury— could reveal targetable pathways for interrupting this pathogenic cycle.

Spatial proteomics and transcriptomics enable comprehensive tissue mapping at single-cell resolution [22–24], offering opportunities to identify targetable pathways in AIN. We applied imaging mass cytometry (IMC, a spatial proteomics technique) and single cell spatial transcriptomics (ST) to kidney tissues from patients with pathologist-adjudicated AIN, acute tubular injury (ATI, a non-immune-mediated AKI comparator), and reference tissue to: (1) identify disease-specific tubular injury phenotypes and their immune interactions, (2) map inflammatory signaling networks unique to AIN, (3) define the cells and signals that underlie the spatial organization of immune infiltrates, and (4) correlate molecular patterns with clinical outcomes to identify therapeutic targets.

## Results

### Spatial proteomics and transcriptomics reveal distinct cellular architectures in AIN and ATI

We analyzed 106 kidney tissues from patients with pathologist-adjudicated AIN (n=37), ATI (n=37), and reference tissue (n=32) across discovery (n=65) and validation (n=41) cohorts (**Extended Data Fig. 1a,b**). Most patients with AIN received corticosteroid treatment after biopsy (77% in discovery cohort). Clinical parameters including baseline eGFR, nadir eGFR, and 6-month eGFR change were otherwise comparable between AIN and ATI in both cohorts (**Extended Data Fig. 1c**), although validation reference tissues had lower baseline eGFR and higher injury scores compared to discovery reference tissues (**Supplementary Tables 1,2**), likely reflecting underlying kidney disease.

IMC using 31 antibodies (**Supplementary Table 3**) generated single-cell spatial proteomic profiles from 2,565,136 cells across all 106 kidney tissues (**Fig. 1a, Extended Data Fig. 2a**). Unsupervised clustering defined 49 clusters simplified to 18 core cell populations and two mixed cell populations (**Fig. 1c; Extended Data Fig. 2b; Supplementary Table 4**), that were reproduced in the validation cohort (**Extended Data Fig. 1d**). Single-cell ST was performed on 20 tissues from the IMC discovery cohort (8 AIN, 7 ATI, 5 reference), identifying 321,318 cells with transcriptional designation of 26 core cell types and one mixed cell type (**Fig. 1b,d; Extended Data Fig. 3a,b; Supplementary Tables 5,6**).

**Figure 1.**
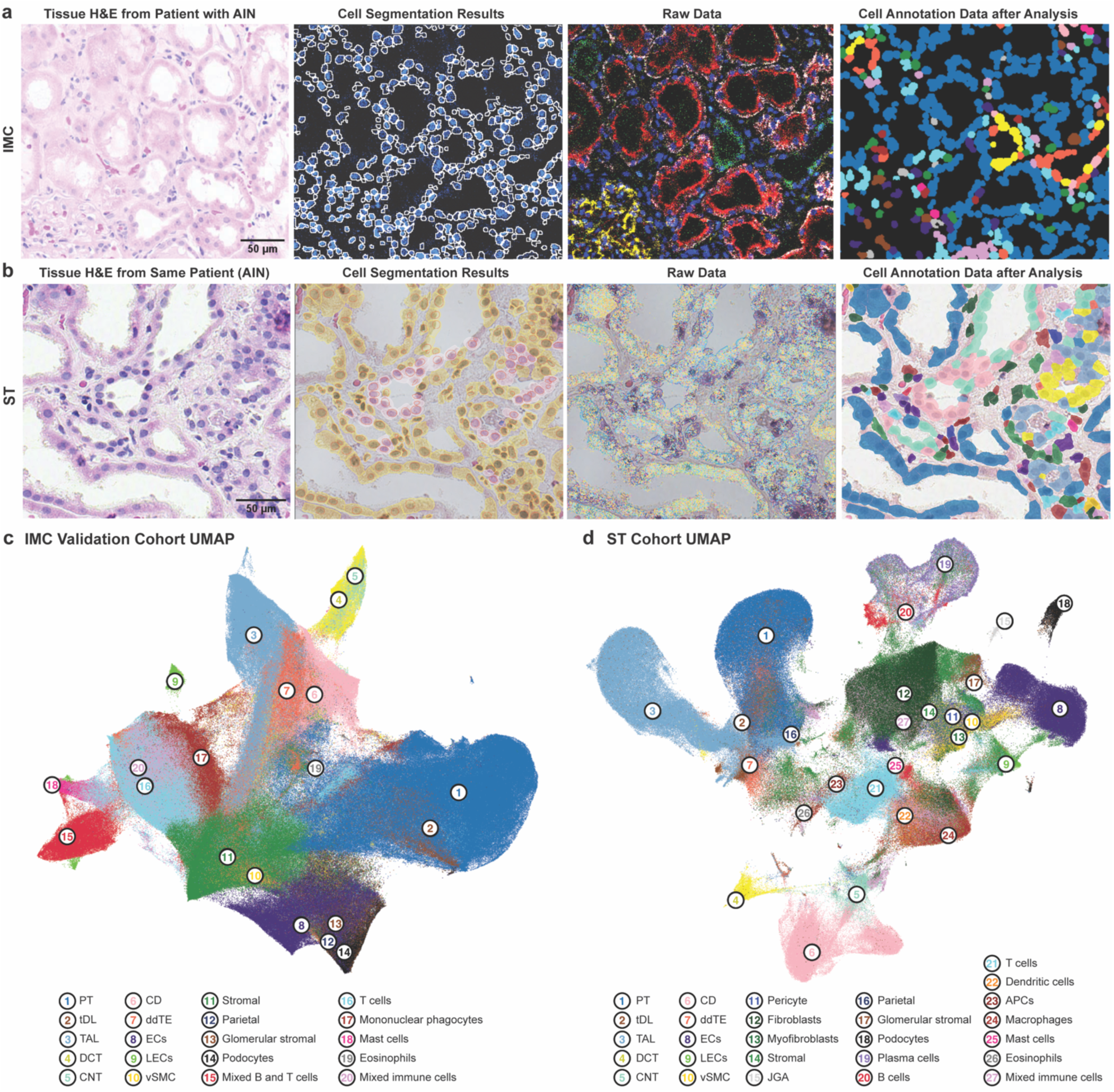
Multi-modal spatial profiling reveals conserved cellular architecture across imaging and transcriptomic platforms in acute kidney injury. **a,** Imaging mass cytometry (IMC) workflow for kidney tissue. Left to right: H&E morphology of AIN tissue biopsy; IMC cell segmentation mask (white outline) from adjacent tissue section overlaid on digitally pseudo-colored DNA intercalator (blue); digitally pseudo-colored protein expression for tubular markers (DNA intercalator for nuclei, blue; megalin and aquaporin 1 for proximal tubule, red; beta-catenin for pan-tubule marker, white; calbindin for distal convoluted tubule, green; ERG for endothelial cell nuclei, cyan; nestin for podocytes, yellow); final cell type annotations with colors corresponding to cell type annotation colors in panel c, notably with de-differentiated tubular epithelium in orange. Scale bar: 50μm. Images generated using MCD Viewer (Standard BioTools, v1.0.560.6). **b,** Spatial transcriptomics (ST) workflow for kidney tissue, showing matched AIN tissue from same patient as in panel a. Left to right: H&E morphology of tissue section; cell segmentation algorithm results showing nuclear (blue), interior (pink), and boundary (yellow) detection methods with nuclear (red) segmentation also displayed overlaid on H&E; raw transcript expression (all transcripts) overlaid on H&E; final cell type annotations with colors corresponding to cell type annotation colors in panel d overlaid on H&E. Scale bar: 50μm. Image generated using 10x Genomics Xenium Explorer v4.0.0. **c,** Uniform Manifold Approximation and Projection (UMAP) embedding of 1,331,664 cells from IMC validation cohort (n=65 patients). Numbers indicate major cell populations: (1) PT, proximal tubule; (2) tDL, thin descending limb; (3) TAL, thick ascending limb; (4) DCT, distal convoluted tubule; (5) CNT, connecting tubule; (6) CD, collecting duct; (7) ddTE, dedifferentiated tubular epithelium; (8) EC, endothelial cells; (9) LEC, lymphatic endothelial cells; (10) vSMC, vascular smooth muscle cells; (11) Stromal; (12) Parietal; (13) Glomerular stromal; (14) Podocytes; (15-20) Immune populations. **d,** Uniform Manifold Approximation and Projection (UMAP) embedding of 321,318 cells from ST cohort (n=20 patients) demonstrating concordant cell type identification across platforms. Numbered annotations as in c with addition of (11) Pericyte, (12) Fibroblasts, (13) Myofibroblasts, (15) JGA, (19) Plasma cells, (22) Dendritic cells, (23) APC, antigen presenting cells.

### Disease-specific tubular injury states orchestrate distinct immune neighborhoods

IMC-based analysis of PT cells revealed five distinct states: healthy (PT-H; megalin^high^AQP1^high^), dedifferentiated (PT-dd; megalin^low^AQP1^low^), proliferating (Ki-67^+^), and two injury states: PT-INJ1 (KIM-1^+^VCAM1^−^) and PT-INJ2 (KIM1^±^VCAM1^+^) (**Fig. 2a**). Examination of the adjacent H&E-stained section confirmed that PT-H cells appeared normal whereas PT-INJ1 cells had loss of brush border, cytoplasmic vacuolization, and nuclear drop-out, and PT-INJ2 cells exhibited epithelial flattening in dilated tubules with peritubular scarring (**Fig. 2b**). Blinded pathologist scoring confirmed that PT-INJ1 cells exhibited higher injury scores than PT-H while PT-INJ2 were scored as atrophic (**Extended Data Fig. 4a**).

**Figure 2.**
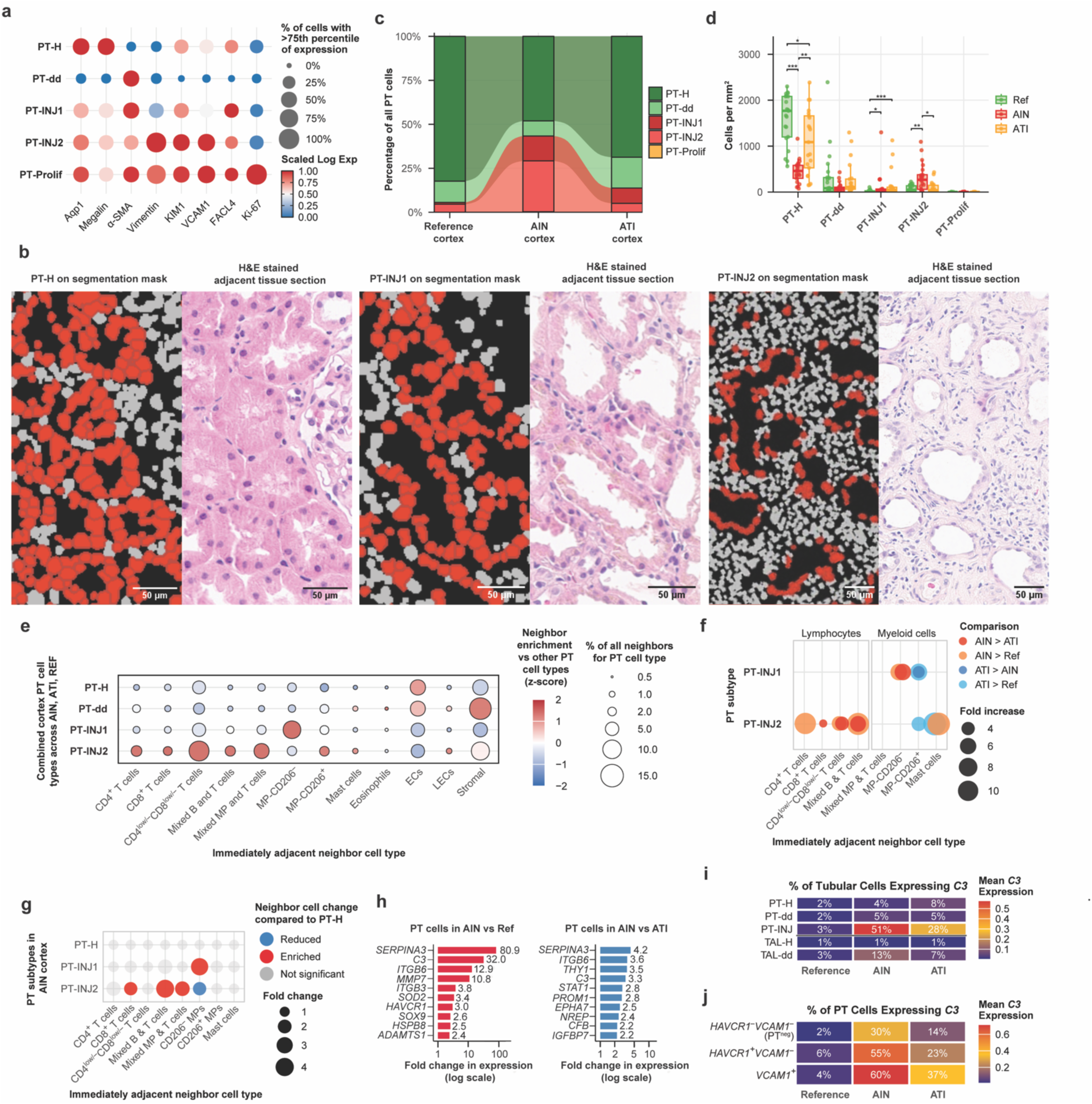
Proximal tubule injury states exhibit distinct cellular neighborhoods and complement expression patterns. **a,** Protein marker expression across PT subtypes identified by imaging mass cytometry (IMC) in cortex tissues in discovery cohort (n=63, AIN=21, ATI=21, Ref=21). Dot size: percentage of cells expressing marker at above the 75th percentile of all tubular epithelium. Color: log-transformed, min-max marker scaled expression. **b,** Morphological correlation of IMC-defined PT states. Cell segmentation masks (left) with PT subtypes highlighted (red) and corresponding H&E sections (right). Scale bars: 50μm. **c,** PT population proportional shifts across disease states in cortex tissues in discovery cohort. Height represents proportion of total PT cells. n=63 cortex biopsies. **d,** PT subtype densities (cells/mm²) across reference (green, n=21), AIN (red, n=21), and ATI (orange, n=21) cortex tissues. Wilcoxon test with Bonferroni correction: *p<0.05, **p<0.01, ***p<0.001. **e,** Cellular composition within 20μm of PT subtypes in cortex tissues. Dot size: percentage of neighbors; color: z-score enrichment relative to mean across subtypes. Analysis combines all cortical regions across tissue types (n=63, AIN=21, ATI=21, Ref=21). **f,** Disease-specific PT subtype neighbor enrichment in cortex tissues (n=63, AIN=21, ATI=21, Ref=21). Only significant interactions shown (q<0.05), statistical significance determined using mixed models.. Dot size: fold change; color: comparison direction. **g,** PT subtype neighbor changes within AIN cortex. PT-INJ1 and PT-INJ2 compared to PT-H, statistical significance determined using mixed models. Red: enriched, blue: reduced, gray: non-significant (q≥0.05). n=21 AIN patients. **h,** Top 10 differentially expressed genes in PT cells from AIN cortex tissue (n=7) versus reference cortex (left, n=2, descriptive comparison only) and versus ATI cortex (right, n=6, adjusted p<0.05, Wilcoxon test with Bonferroni correction) tissues. **i,** Complement *C3* expression across PT and thick ascending limb of loop of Henle (TAL) subtypes in cortex tissues (n=15, AIN=7, ATI=6, Ref=2). Color: mean normalized expression (log1p); percentages: C3-positive cells. TAL stratified by uromodulin: TAL-H (>median *UMOD* expression), TAL-dd (≤median *UMOD* expression). **j,** Complement *C3* expression across PT cells stratified by *HAVCR1*/*VCAM1* transcriptional expression (n=15, AIN=7, ATI=6, Ref=2). Color: mean normalized expression (log1p); percentages: C3-positive cells. **Abbreviations**: PT, proximal tubule; MP, mononuclear phagocyte; EC, blood vascular endothelial cell; LEC, lymphatic endothelial cell.

VCAM1^+^ PT-INJ2 cells comprised 28.9% of PT in AIN (vs 5.1% ATI, 4.6% reference) and PT-INJ1 14.2% (vs 8.7% ATI, 1.0% reference) (**Fig. 2c; Extended Data Fig. 4b**). Overall PT density was reduced in AIN, and these proportional changes translated to 6.5-fold higher PT-INJ2 cells/mm^2^ in AIN compared to ATI (median 267 vs 41 cells/mm²; adj. p=0.013), while PT-INJ1 density showed no significant difference between AIN and ATI (**Fig. 2d; Extended Data Fig. 4c**). These relationships were maintained in the validation cohort except for higher PT-INJ1 cells in reference samples (**Extended Data Fig. 4d-f**).

IMC-based characterization of immediately adjacent cell neighbors revealed unique neighborhood compositions around distinct PT states. PT-H cells had endothelial cells as their most common non-tubular neighbor, whereas PT-dd had more stromal cell neighbors (**Fig. 2e**). PT-INJ2 cells showed enrichment of multiple lymphocyte neighbors, while PT-INJ1 cells showed enrichment of mononuclear phagocyte (including macrophage) neighbors with disease-specific polarization patterns (**Fig. 2f, Extended Data Fig. 5a**). Analysis of immediate cell neighbors within AIN cortex alone showed increased spatial proximity of distinct immune populations was specific to the two PT injury states relative to healthy PT cells (**Fig. 2g, Extended Data Fig. 5b**).

To better characterize these PT injury states, we subclustered 31,707 cortical PT cells from ST data into healthy (PT-H), dedifferentiated (PT-dd) and injured (PT-INJ) states (**Supplementary Tables 6,7**), and additionally stratified cells by *HAVCR1* and *VCAM1* expression to parallel IMC-defined injury states. Within AIN cortex alone, the transition from *HAVCR1*^−^*VCAM1*^−^ PT (PT^neg^) to *HAVCR1*^+^*VCAM1*^−^ to *VCAM1*^+^ PT cells revealed stepwise transcriptional changes (**Extended Data Fig. 6**). *HAVCR1*^+^*VCAM1*^−^ cells exhibited 157 differentially expressed genes (DEGs) compared to PT^neg^ cells, including upregulation of injury markers (*VIM*, *C3*, *ITGB3*) and downregulation of transport genes. *VCAM1*^+^ PT cells showed 232 DEGs, with prominent upregulation of complement genes (*C3*, *C4A*, *C4B*, *CFB*), MHC class II-related gene *CD74*, and survival pathways (*THY1, SERPINA3*).

Analysis of injured PT cells revealed a robust AIN-enriched inflammatory signature shared across both *HAVCR1*^+^*VCAM1*^−^ and *VCAM1*^+^ states compared to ATI, with upregulation of *SERPINA3* (7.5- and 7.1-fold respectively), LCN2 (3.7- and 3.3-fold), C3 (3.3- and 2.3-fold), and *STAT1* (2.7- and 2.9-fold, all p<0.0001) (**Extended Data Fig. 6**). While this core inflammatory program was shared, *HAVCR1*^+^*VCAM1*^−^ cells specifically upregulated *GBP1* (3.5-fold), an interferon-induced GTPase, whereas *VCAM1*^+^ cells showed upregulation of *IL1R1* (2.1-fold) and *GADD45B* (2.2-fold), suggesting enhanced cytokine signaling and stress response. *C3* emerged as a defining feature across injury states in AIN, suggesting tubular cell-expressed complement might help orchestrate immune recruitment in AIN.

### Local complement synthesis drives disease-specific immune recruitment

Given the emergence of *C3* as a defining AIN feature, we quantified its cellular sources. Tubular epithelial cells contributed 63.7% of total *C3* mRNA in AIN cortex, with immune cells contributing 21.9% (**Extended Data Fig. 3b**), aligning with histologic observations of C3 protein deposition along tubules in AIN [25–28]. AIN samples showed significantly higher *C3* expression than ATI across all proximal tubule cells (mean normalized expression 0.441 vs 0.160, p<0.0001). Among PT subpopulations in AIN, expression progressively increased from PT^neg^ cells to *HAVCR1*^+^*VCAM1*^−^ to *VCAM1*^+^ cells (**Fig. 2i,j**), with *VCAM1*^+^ PT cells also upregulating *C4A, C4B,* and *CFB* (**Extended Data Fig. 6**). Consistent with recent observations in AKI and CKD [22], AKI samples showed increased numbers of thick ascending limb cells with diminished UMOD expression (dedifferentiated TAL, TAL-dd) [29, 30]. In AIN, TAL-dd comprised 87.1% of TAL (versus 7.6% in reference) with 23-fold higher *C3* expression than healthy TAL and 3.6-fold higher than TAL-dd in ATI.

At the tissue level, tubular *C3* expression correlated with *C3AR1*^+^ immune density (r=0.91, p=0.00483, **Fig. 3a**), with AIN macrophages expressing higher *C3AR1* mRNA than those found in ATI samples (mean normalized expression 0.086 vs 0.060, p<0.0001). Using a 20μm distance threshold, predicted paracrine C3-C3AR1 interactions were 3.4-fold higher in AIN than reference and 1.5-fold higher than ATI (median 74.1 vs 21.6 and 48.6 interactions/mm², respectively; **Fig. 3b**), with tubular, immune, and stromal cells as *C3^+^* sources and *C3AR1*^+^ mononuclear phagocytes (MPs, specifically macrophages) as the primary targets (**Fig. 3c**). *C3AR1*^+^ macrophages adjacent to *C3*^+^ PT were predominantly M1 antigen-presenting macrophages (51% of PT-macrophage C3-C3AR1 interactions), suggesting direct complement-mediated recruitment. This enrichment of M1 activated, *C3AR1*^+^ MPs around injured PT cells aligned with the MP accumulation and activation states found in our IMC analysis, with tissue MP density of 120 cells/mm² in AIN versus 63 cells/mm² in ATI and 26 cells/mm² in reference tissues (p<0.05 and p<0.001, respectively; **Fig. 3d; Extended Data Fig. 4c**), findings confirmed in the validation cohort (**Extended Data Fig. 4g**). Specifically, PT-INJ1 cells in AIN exhibited a 3.5-fold enrichment of MP neighbors lacking the CD206 (mannose receptor) alternative activation marker, while PT-INJ1 cells in ATI showed CD206^+^ MP neighbor enrichment (**Fig. 2f,g; Extended Data Fig. 5a,b**).

**Figure 3.**
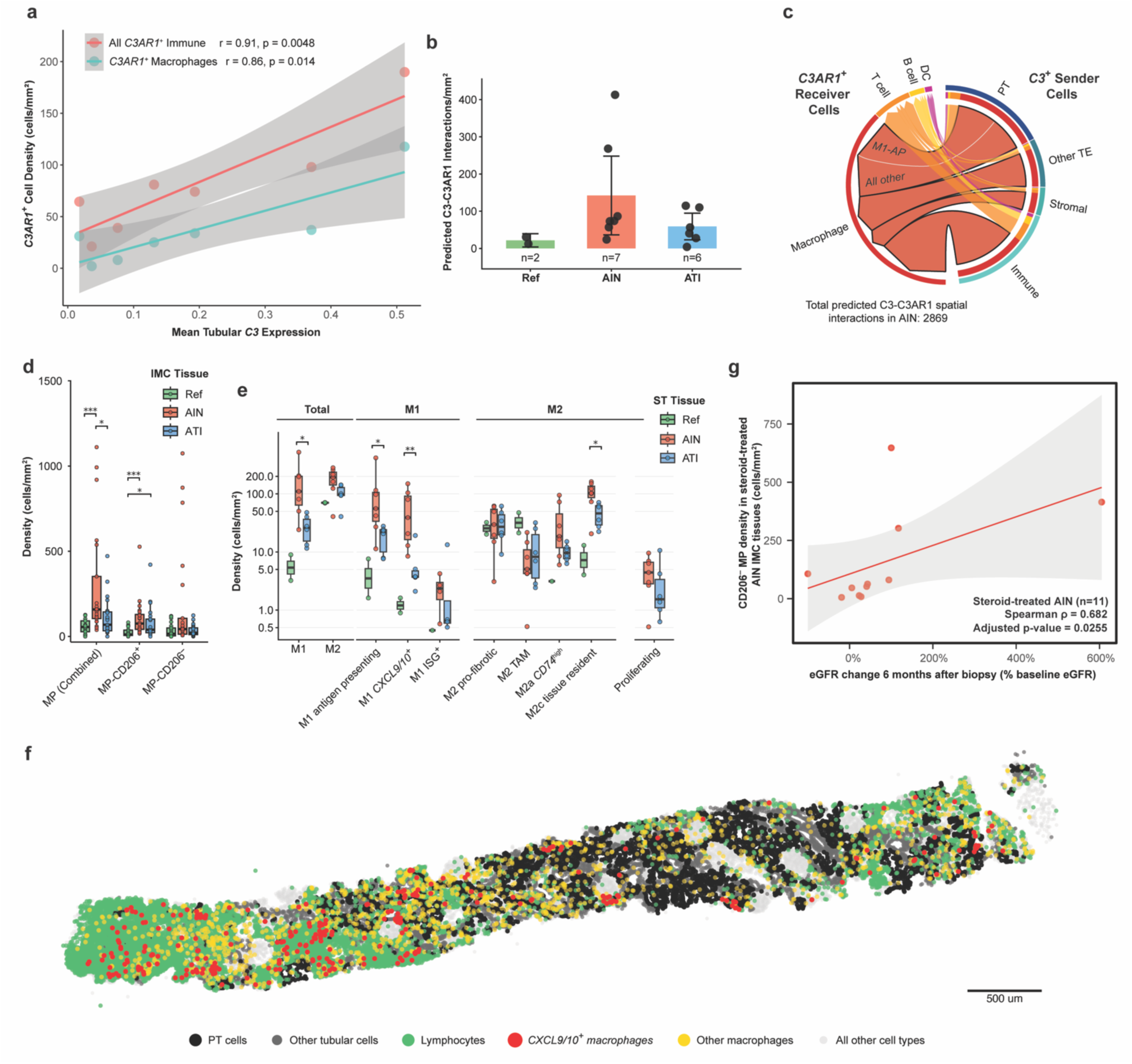
Tubular complement production drives spatially-organized macrophage recruitment with opposing prognostic roles in AIN versus ATI. **a,** Tubular *C3* expression correlates with *C3AR1*⁺ immune infiltration in AIN cortex tissues (n=7). All immune cells (red): r=0.91, p=0.0048; macrophages alone (cyan): r=0.86, p=0.014. Pearson correlation with 95% CI. **b,** Predicted C3-C3AR1 paracrine interaction density (interactions per mm²) in kidney cortex. Cell pairs with detectable *C3* and *C3AR1* expression within 20μm classified as predicted interactions. Expression of both *C3* and *C3AR1* in the same cell (potential autocrine signaling) not included. Data shown as mean ± 95% confidence interval with individual patient values overlaid. Reference (green, n=2): 21.6 interactions/mm² (descriptive only due to limited sample size). ATI (orange, n=6): mean 59.0, median 48.6 interactions/mm². AIN (red, n=7): mean 142.3, median 74.1 interactions/mm². **c,** Chord diagram of predicted C3-C3AR1 spatial interactions (≤20μm) in AIN cortex tissues (n=7). Width proportional to interaction frequency. Arrows from *C3*⁺ senders (right) to *C3AR1*⁺ receivers (left). PT-Macrophage C3-C3AR1 predicted interactions split into PT-M1 antigen-presenting (M1-AP) interactions (51%) and all other PT-Macrophage interactions. Total 2,869 predicted interactions (AIN cortex, n=7). **d,** Imaging mass cytometry quantification of mononuclear phagocyte (MP) populations in cortex tissues in the discovery cohort. Reference (n=21), AIN (n=21), ATI (n=21). *p<0.05, **p<0.01, ***p<0.001; Wilcoxon test with Bonferroni correction. **e,** Spatial transcriptomic profiling of macrophage polarization states in cortex tissues. Total M1/M2 and detailed subtypes in AIN (red, n=7), ATI (blue, n=6), and Reference (green, n=2, descriptive only due to limited sample size). Y-axis log₁₀ scale. Statistics performed only on AIN vs ATI. *p<0.05, **p<0.01; Wilcoxon test. **f,** Representative image of macrophage spatial distribution in AIN cortex tissue. Proximal tubules (black), other tubular cells (dark gray), *CXCL9*/*10*⁺ inflammatory macrophages (red), other macrophages (yellow), lymphocytes (green), all other cell types (light gray). Scale bar: 500μm. **g,** CD206⁻ MP density predicts steroid responsiveness in AIN cortex tissues from discovery cohort (n=11; all patients with AIN who did receive steroid treatment with available baseline eGFR, nadir eGFR, and 6-month eGFR data were included). Spearman ρ=0.682, adjusted p=0.0255. eGFR recovery calculated as percentage of baseline function restored at 6 months. **Abbreviations**: Ref, reference tissue; DC, dendritic cell; TE, tubular epithelium; PT, proximal tubule cells; MP, mononuclear phagocyte; ISG, interferon-stimulated genes; TAM, tumor-associated macrophage; M1-AP, M1 antigen-presenting macrophage.

ST analysis confirmed macrophage enrichment in AIN (**Extended Data Fig. 7a**) and revealed M1 predominance (median 116 vs 27 cells/mm² in ATI, 4.3-fold higher, p=0.014), consisting primarily of antigen-presenting and *CXCL9/10*^+^ M1 subtypes (**Fig. 3e**). Despite 10.6-fold higher overall density of *CXCL9/10*^+^ macrophages in AIN versus ATI, these cells comprised only 7.9% of predicted C3-C3AR1 interactions with PT cells, representing a 2-fold reduction relative to their tissue abundance. Proximity analysis confirmed this spatial avoidance: *CXCL9/10*^+^ macrophages represented 12.6% of macrophages near PT cells versus 15.5% in distant regions (p=0.0016) (**Fig. 3f**), suggesting *CXCL9/10*^+^ macrophage enrichment in a distinct cellular compartment.

ATI samples showed lower overall macrophage density with tissue-resident and pro-fibrotic M2 phenotypes predominating. These distinct macrophage polarization patterns in AIN and ATI predicted opposite clinical outcomes by disease. In steroid-treated AIN, CD206^−^ MP density predicted improved recovery (n=11, ρ=0.682, p=0.026, **Fig. 3g**), while overall MP density predicted worse outcomes in patients with ATI (ρ=-0.564, p=0.020, **Extended Data Fig. 7b**).

### Microenvironmental signatures reveal progression from inflammatory to fibrotic programs

To characterize PT injury-associated microenvironments, we applied SORBET [31] to identify transcripts within a two cell distance radius most predictive of PT cell phenotypes, complemented by DEG analysis within 50 μm (**Fig. 4a; Extended Data Fig. 8a-f; Supplementary Fig. 1-4**). In AIN cortex, *HAVCR1*^+^*VCAM1*^−^ PT microenvironments were best predicted by transcripts coordinating acute inflammation: *IL7R* (SORBET association score 1.00, 1.6-fold upregulation compared to PT^neg^ microenvironment), *SOCS3* (score 0.93, 1.3-fold upregulated), and *CXCL9* (score 0.89, 1.3-fold upregulated). With progression to a *VCAM1*^+^ state, microenvironments showed metabolic-fibrotic transition, with *NAMPT* (score 1.00, 1.9-fold upregulated) and *COL4A1* (score 0.60, 1.6-fold upregulated) expression. Consistent with the IMC finding that *VCAM1*^+^ cells closely associated with lymphocyte populations in AIN (**Fig. 2f,g**), *MS4A1* (CD20) and *CD8A* expression were also highly enriched in and predictive of the *VCAM1*^+^ microenvironment (**Fig. 4a**).

**Figure 4.**
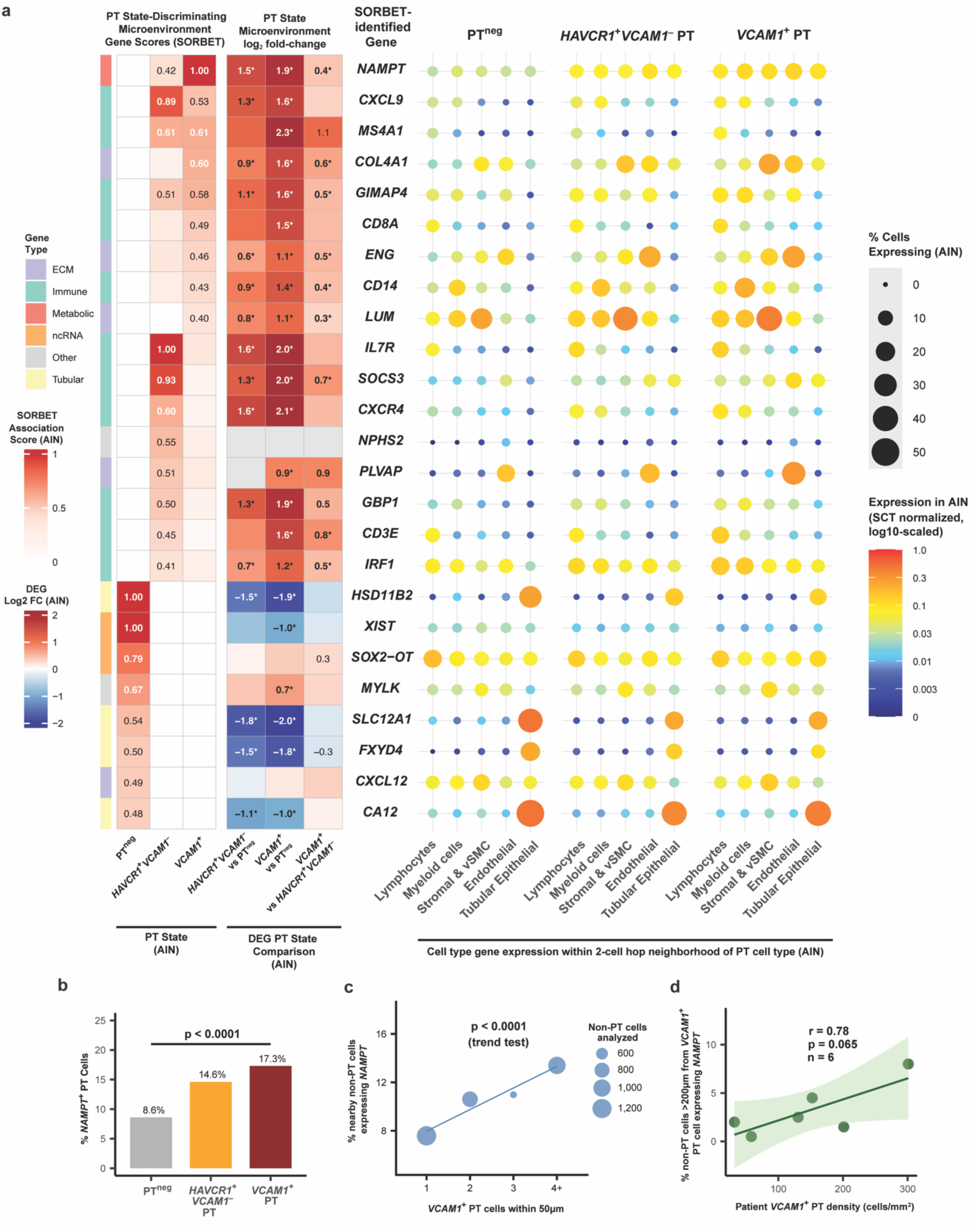
SORBET machine learning-guided spatial analysis reveals hierarchical inflammatory programs in kidney injury microenvironments in AIN cortex tissues. **a,** Analysis of PT injury microenvironments in AIN cortex tissues (n=7) for the top SORBET-identified genes (middle) using SORBET-derived spatial association scores (left) and an analysis of differential gene expression for cellular neighbors within PT injury microenvironments (right). Left panels: SORBET association scores (≥0.40 threshold) identify genes (middle) predictive of specific PT states (*HAVCR1*^−^ *VCAM1*^−^ (PT^neg^), *HAVCR1*⁺*VCAM1*⁻, or *VCAM1*⁺), paired with log₂ fold-change values from differential expression (DEG) analysis of non-PT cells within 50μm microenvironments surrounding PT cell subtypes. Statistical significance of DEGs is indicated by text formatting of log₂ fold-change values using Bonferroni-adjusted p-values: bold with asterisk (*) for p<0.001, bold for p<0.01, and plain text for p<0.05. Log₂ fold-change value not shown where non-significant. Genes are organized by ranked combined SORBET association score between PT injury states, with genes scoring highest for *VCAM1*⁺ PT on top, followed by *HAVCR1*⁺*VCAM1*^−^ and PT^neg^. Right panels: Grouped cell type-specific expression patterns within each PT subtype microenvironment show progressive inflammatory reprogramming from PT^neg^ to *VCAM1*⁺ states, with coordinate upregulation of immune mediators and extracellular matrix components. Dot size indicates the percentage of each grouped cell type expressing the gene within each microenvironment, and color indicates normalized log10-scaled expression of gene for given grouped cell type within PT subtype microenvironment. **b,** Cell-autonomous *NAMPT* expression in cortical PT cells stratified by *HAVCR1*/*VCAM1* expression in AIN (n=7). *NAMPT* positivity progressively rises from PT^neg^ (8.6%) through *HAVCR1*⁺*VCAM1*⁻ (14.6%) to *VCAM1*⁺ PT cells (17.3%; p<0.0001, chi-square test). **c,** Local paracrine regulation of *NAMPT* in AIN cortex tissues (n=7). Non-PT cells show dose-dependent higher *NAMPT* expression based on number of neighboring *VCAM1*⁺ PT cells within 50μm. Dot size proportional to number of non-PT cells analyzed per density category. P<0.0001, Cochran-Armitage trend test for increasing *NAMPT* expression with *VCAM1*⁺ PT cell density. **d,** Regional tissue-level *NAMPT* regulation in AIN cortex tissues. Percentage of *NAMPT*⁺ cells >200μm from any *VCAM1*⁺ PT correlates with whole tissue-level *VCAM1*⁺ PT density (r=0.78, p=0.065, Pearson correlation; n=6 patients), suggesting tissue-wide *NAMPT* upregulation scales with *VCAM1*^+^ PT burden. 1 patient with AIN cortex tissue excluded due to insufficient (<100) cells >200μm from any *VCAM1*⁺ PT (high tissue *VCAM1*⁺ PT burden). **Abbreviations**: PT, proximal tubule; DEG, differentially expressed gene; ECM, extracellular matrix; ncRNA, non-coding RNA; vSMC, vascular smooth muscle cells; SCT, SCTransform.

Expression of the NAD+ biosynthetic enzyme and inflammatory cytokine *NAMPT* was the strongest predictor of *VCAM1*^+^ PT cell microenvironments in AIN (**Fig 4a**), with stepwise transcriptional upregulation with injury severity (microenvironment DEG 1.5 for *HAVCR*^+^*VCAM1*^−^, 1.9 for *VCAM1*^+^) across virtually all neighbor cell types. PT cells also demonstrated stepwise *NAMPT* upregulation with injury (8.6% → 14.6% → 17.3% expression from PT^neg^ to *VCAM1*^+^, p<0.0001, **Fig. 4b**). The level of *NAMPT* expression showed a correlation with *VCAM1*^+^ PT cells at both a local (**Fig. 4c**) as well as regional level (**Fig. 4d**), indicating field effects extending beyond local microenvironments. *CXCL9* emerged as a consistent inflammatory mediator across lymphocyte, myeloid, and stromal populations in AIN, with progressive upregulation in the microenvironment of injured PT (1.3-fold near *HAVCR*^+^*VCAM1*^−^ PT and 1.6-fold near *VCAM1*^+^ PT, **Fig 4a**). This association of distinct T cell signaling genes with *VCAM1*^+^ versus *HAVCR1*^+^*VCAM1*^−^ microenvironments prompted examination of associated lymphocyte subsets.

### T cell infiltration and CXCL9-CXCR3 networks define AIN inflammation

IMC analysis showed T cell populations were markedly elevated in AIN versus ATI and reference tissues (**Fig. 5a; Extended Data Fig. 4c,g**). These cells localized around VCAM1^+^ tubules, with PT-INJ2 cells in AIN showing dramatic lymphocyte recruitment: mixed B and T cell aggregates (5.7-fold vs PT-INJ2 in ATI, p=0.001) and CD8^+^ T cells (2.7-fold vs PT-INJ2 in ATI, p=0.002) as immediate neighbors (**Fig. 2f,g; Extended Data Fig. 5a,b**). PT-INJ1 cells showed no significant lymphocyte enrichment, suggesting VCAM1 expression specifically aligns with lymphocyte recruitment.

**Figure 5.**
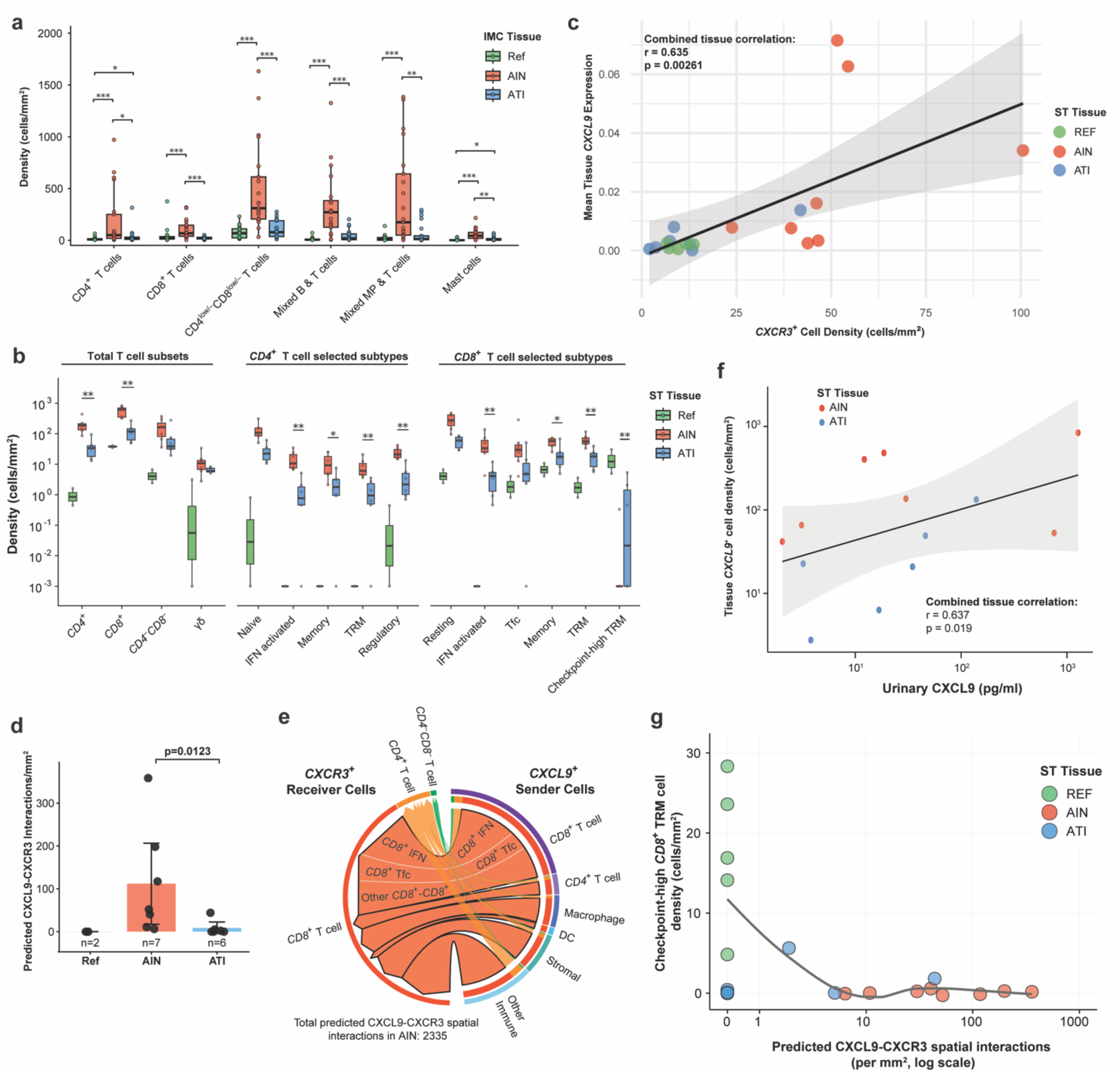
Spatial profiling reveals CXCL9-CXCR3 inflammatory networks and immune cell dynamics in acute kidney injury. **a,** Imaging mass cytometry quantification of cortical immune cell densities across 63 kidney biopsies with cortex tissue. Cell densities (cells/mm²) shown for CD4⁺ T cells, CD8⁺ T cells, CD4 ^low/⁻^CD8^low/⁻^ T cells, mixed B and T cell aggregates, mixed mononuclear phagocyte (MP) and T cell infiltrates, and mast cells comparing reference (Ref, green, n=21), acute interstitial nephritis (AIN, red, n=21), and acute tubular injury (ATI, blue, n=21) tissues. ***p<0.001, **p<0.01, *p<0.05; Wilcoxon test with Bonferroni correction. **b,** Spatial transcriptomic profiling of T cell populations and phenotypic subtypes across 15 patients with cortex tissue. Left: Total T cell subset densities showing 5.6-fold expansion in AIN versus ATI. Middle: *CD4*⁺ T cell subtypes including naive, IFN-activated, memory, tissue-resident memory (TRM), and regulatory populations. Right: *CD8*⁺ T cell subtypes including resting (45% of *CD8*⁺), IFN-activated, T follicular cytotoxic (Tfc, 18%), memory, TRM, and checkpoint-high TRM populations. n=2 Ref, n=7 AIN, n=6 ATI. Y-axis log₁₀ scale. **p<0.01, *p<0.05; Wilcoxon test. Reference tissue populations included for descriptive comparison only without statistical analysis given limited sample size. **c,** Tissue *CXCL9* expression correlates with *CXCR3*⁺ cell density across all disease categories (n=20, Pearson r=0.635, p=0.00261). Gray shading indicates 95% confidence interval. **d,** Predicted CXCL9-CXCR3 paracrine interaction density (interactions per mm²) in kidney cortex. Cell pairs with detectable *CXCL9* and *CXCR3* expression within 20μm classified as predicted interactions. Expression of both *CXCL9* and *CXCR3* in same cell (potential autocrine signaling) not included. Data shown as mean ± 95% confidence interval with individual patient values overlaid. Reference (green, n=2, descriptive only): mean 0.0, median 0.0 interactions/mm². ATI (orange, n=6): mean 8.6, median 1.2 interactions/mm², IQR: 0.1-4.3. AIN (red, n=7): mean 111.9, median 52.1 interactions/mm², IQR: 25.6-157.6. Statistical comparison performed between AIN and ATI cortex tissues only using Wilcoxon rank-sum test. **e,** Chord diagram of predicted CXCL9-CXCR3 spatial interactions for spatially proximate cells (within 20μm) in AIN cortex tissues (44-fold increased vs ATI). Width proportional to interaction frequency. Arrows from *CXCL9*⁺ senders (right) to *CXCR3*⁺ receivers (left). *CD8*^+^ T cell to *CD8*^+^ T cell CXCL9-CXCR3 interactions split into *CD8*^+^ IFN activated-*CD8*^+^ IFN activated (49.2%), *CD8*^+^ Tfc-*CD8*^+^ Tfc (19.5%), and all other *CD8*^+^ T cell-*CD8*^+^ T cell interactions. Total 2,335 interactions (AIN cortex, n=7). **f,** Urinary CXCL9 correlates with cortex tissue *CXCL9*⁺ cell density in cortex tissues (Pearson r=0.637, p=0.019) in paired samples from AIN (red, n=7) and ATI (blue, n=6) patients. Reference samples not included as urinary CXCL9 not available. Both axes log₁₀ scale. **g,** Relationship between CXCL9-CXCR3 interaction density and checkpoint-high *CD8*^+^ TRM cell presence (n=20). Reference tissues (green, n=5) maintain checkpoint-high *CD8*^+^ TRM cells with minimal inflammatory interactions. AIN (n=8) and ATI (n=7) show inverse correlation, with complete checkpoint-high *CD8*^+^ TRM cell absence in all patients exceeding 50 interactions/mm². LOESS curve fitted to non-zero values. X-axis log scale. **Abbreviations**: MP, mononuclear phagocyte; Ref, reference tissue; IFN, interferon; TRM, tissue-resident memory; Tfc, follicular cytotoxic T cell; DC, dendritic cell.

ST analysis confirmed T cells as the dominant immune cell in AIN, with 5.6-fold elevation in median cell density versus ATI (1271 vs 227 cells/mm²) (**Fig 5b**; **Extended Data Fig. 7a**). *CD8*^+^ T cells comprised 62% of the T cell infiltrate (median 465 cells/mm²), predominantly a resting subset (54% of *CD8*^+^ T cells, **Extended Data Fig. 3d**), tissue-resident memory (TRM, 9.3%), and memory (8.8%) *CD8*^+^ T cells. Interferon (IFN)-activated *CD8*^+^ T cells showed the highest enrichment in density versus ATI (8.1-fold, p<0.01). Follicular cytotoxic *CD8*^+^ T cells (Tfc, 6.5%) were also prominent. The *CD4*^+^ compartment included IFN-activated *CD4*^+^ T cells (12.3-fold higher cell density than ATI, p<0.01), and *FOXP3*^+^ regulatory T cells (8.7-fold higher cell density than ATI, p<0.01) (**Extended Data Fig. 3c**). Despite their relative enrichment in AIN, *CD4*^+^ regulatory T (*FOXP3*^+^) cells expressing *TIGIT* [32] comprised only 2.3% of total T cells—well below the 5-10% expected in healthy tissue.

Since urinary CXCL9 is an AIN-specific biomarker [33], we examined cellular *CXCL9* sources and their proximity to CXCL9 receptor (*CXCR3*)-expressing cells. There was a significant correlation between tissue *CXCL9* expression and *CXCR3*^+^ cell density (r=0.646, p=0.001, **Fig. 5c**), with *CD8*^+^ T cells and macrophages contributing equally as *CXCL9* sources. IFN-activated *CD4*^+^ T cells showed highest *CXCR3* expression (11.9%) while TRM cells showed highest *CXCR3* expression among *CD8*^+^ subsets (7.5%). Spatially-constrained (20 μm) paracrine ligand-receptor analysis revealed a 44-fold enrichment in predicted CXCL9-CXCR3 interactions in AIN versus ATI (median 52.1 vs 1.2/mm², p=0.01, **Fig. 5d**), with *CD8*^+^ T cell-to-*CD8*^+^ T cell interactions comprising 41.2% of all predicted interactions (**Fig. 5e**). These were dominated by homotypic amplification loops between follicular cytotoxic (Tfc-Tfc, 19.5% of *CD8*^+^-*CD8*^+^) and IFN-activated *CD8*^+^ T cells (49.2%). Predicted CXCL9-CXCR3 interaction density correlated with *HAVCR1***^+^***VCAM***^−^**PT density (r=0.803, p=0.030), linking tubular injury to inflammatory networks. Urinary CXCL9 correlated with overall *CXCL9*^+^ cell density (r=0.637, p=0.019) rather than interaction density (r=0.113, p=0.714, **Fig. 5f**).

### Absence of checkpoint-high CD8^+^ TRM subset coincides with unchecked inflammatory amplification

Homotypic CXCL9-CXCR3 amplification networks occurred in the setting of a profound reduction of checkpoint-high *CD8*^+^ tissue resident memory (TRM) cells. This subset, characterized by co-expression of multiple inhibitory receptors (*LAG3, TIGIT, PDCD1*) with retained cytotoxic capacity (*PRF1*^+^) but minimal cytotoxic activity (*GZMB*^−^), represents a memory population under multilayered inhibitory control (**Extended Data Fig. 3d**). These cells were present in all 5 reference tissues (median 17.0 cells/mm²) but absent in 88% (7/8) of AIN samples (p=0.005), and their absence correlated with increased inflammatory signaling intensity (**Fig. 5g**). Samples lacking checkpoint-high *CD8*^+^ TRM cells (n=11: 7 AIN, 4 ATI) showed significantly more predicted CXCL9-CXCR3 signaling interactions (median 10.8 interactions/mm²) compared to samples retaining these cells (median 0 interactions/mm², n=9: 1 AIN, 3 ATI, 5 Ref, p=0.05). All patients exceeding 50 CXCL9-CXCR3 predicted interactions/mm² showed complete absence of checkpoint-high *CD8*^+^ TRM cells (4/4 patients, all AIN).

### Lymphoid aggregates spatially propagate inflammatory circuits

The spatial organization of inflammatory signaling in AIN revealed distinct compartmentalization around compact lymphoid aggregates. Using criteria adapted from Liu et al. [34] for identifying tertiary lymphoid structures (TLS), we identified 28 compact, organized lymphoid aggregate regions across all tissues, with disease-specific enrichment: 21 aggregates in 7/8 AIN patients versus 7 aggregates in 2/7 ATI patients (33.3%), and no aggregates in reference tissues (**Fig. 6a**). These aggregates contained B cells, T cells, dendritic cells, and plasma cells, with universal expression of TLS-organizing chemokines *CCL19* and *CCL21*, and the B cell-recruiting chemokine *CXCL13* in 57%. Checkpoint-high *CD8*^+^ TRM cells were absent from 27 of 28 lymphoid aggregates (96.4%), while *CD4*^+^ regulatory T cells were detected in 17 of 28 aggregates (60.7%).

**Figure 6.**
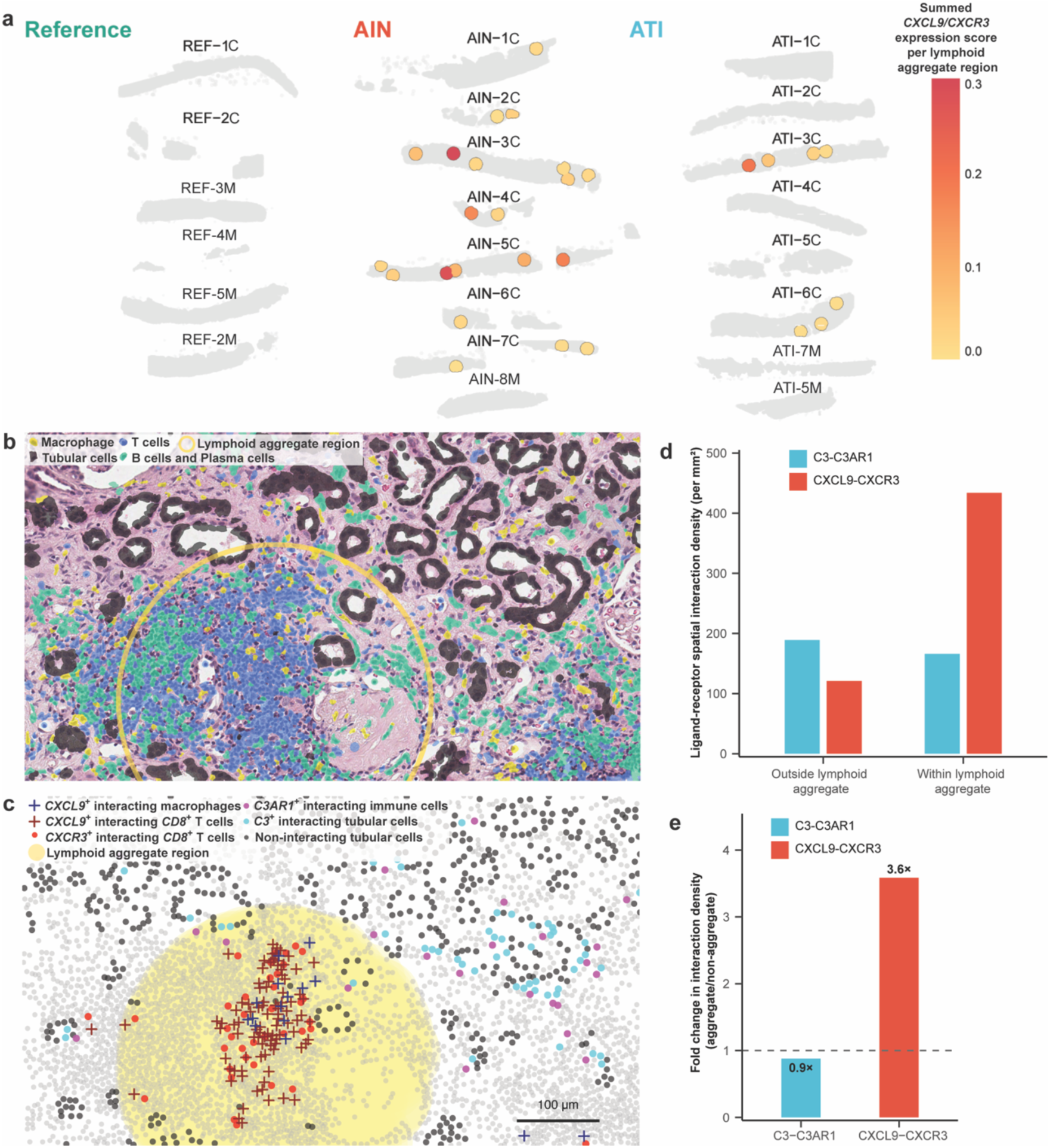
Lymphoid aggregates spatially compartmentalize inflammatory signaling in acute interstitial nephritis. **a,** Tissue-wide mapping of lymphoid aggregates resembling tertiary lymphoid structures (TLS) across 20 kidney biopsies which underwent ST. Reference tissues (green header, n=5) show complete absence of lymphoid aggregates. AIN tissues (red header, n=8) demonstrate 21 aggregates regions in 7/8 patients (87.5%), while ATI (blue header, n=7) shows 7 aggregates in 2/7 patients (28.6%). Colored circles indicate lymphoid aggregate regions with gradient representing summed *CXCL9*/*CXCR3* expression score per region (yellow=low, red=high). Gray areas represent kidney parenchyma. Numbers indicate unique patients within a given tissue type, with M and C designations for medullary and cortical regions; patient tissues REF-2 and ATI-5 had both cortical and medullary regions. **b,** Representative H&E-stained AIN cortex section with cellular segmentation mask with cell type annotation coloring overlay for selected cell types. Color coding: macrophages (yellow), T cells (green), B cells and plasma cells (blue), tubular cells (black). Lymphoid aggregate boundary outlined in yellow demonstrates organized lymphoid architecture with discrete B cell zones, T cell areas, and surrounding tubular structures. Image generated using 10x Genomics Xenium Explorer v4.0.0. **c,** High-magnification spatial mapping of predicted ligand-receptor interactions for spatially proximate cells (within 20μm) within and surrounding a representative lymphoid aggregate (yellow region). Symbols indicate cells participating in predicted spatial interactions within 20μm: CXCL9⁺ macrophages (dark blue crosses), *CXCL9*⁺ *CD8*⁺ T cells (red crosses), *CXCR3*⁺ *CD8*⁺ T cells (red circles), *C3⁺* tubular cells (cyan circles), *C3AR1*⁺ immune cells (magenta circles). Non-interacting cells shown in gray. Note concentration of CXCL9-CXCR3 interactions within lymphoid aggregate boundary versus C3-C3AR1 interactions in surrounding parenchyma. Scale bar: 100μm. **d,** Quantification of predicted ligand-receptor interaction densities (per mm²) comparing regions within versus outside lymphoid aggregate boundaries in AIN cortex tissues (n=7). C3-C3AR1 interactions show no enrichment (189 outside vs 166 within), while CXCL9-CXCR3 interactions demonstrate marked concentration within lymphoid aggregate (121 outside vs 434 within). **e,** Fold change in predicted interaction density comparing lymphoid aggregate to non-aggregate regions in AIN cortex tissues (n=7). CXCL9-CXCR3 shows 3.6-fold enrichment within lymphoid aggregates, while C3-C3AR1 shows 0.9-fold reduction, demonstrating spatial segregation of adaptive (CXCL9-CXCR3) versus innate (C3-C3AR1) inflammatory programs.

Quantitative analysis of predicted ligand-receptor interactions revealed reciprocal patterns for adaptive versus innate immune signaling. Predicted CXCL9-CXCR3 interactions were 3.6-fold higher within lymphoid aggregates (median 433.8 vs 121.0/mm²) (**Fig. 6b-e**), driven by enriched *CXCL9*^+^ cells (5.42% vs 2.62% non-aggregate, p<0.0001) and *CXCR3*^+^ cells (1.14% vs 0.57% non-aggregate, p<0.0001). Per-cell interactions were enriched for *CXCL9*^+^ cells within lymphoid aggregate regions (median 0.80 vs 0.46), with *CD8*^+^ Tfc and *CD8*^+^ IFN-activated T cells dominating. In contrast, predicted C3-C3AR1 interactions were fewer in lymphoid aggregate regions (median 166.0 vs 189.0/mm² in non-aggregate regions). *CXCL9/10*^+^ M1 macrophages that avoided injured tubules (**Fig. 3f**) showed compensatory enrichment within aggregate regions (21.7% vs 17.0% overall macrophage lymphoid aggregate percentage, 1.28-fold enrichment, p<0.001), where they mediated 76.6% of all macrophage-driven CXCL9-CXCR3 interactions. This reciprocal organization–CXCL9-CXCR3 interactions enriched in aggregates, C3-C3AR1 interactions enriched around tubules–suggests lymphoid aggregates act as specialized niches prioritizing T cell-mediated adaptive immunity while complement-driven innate immune recruitment occurs around tubular parenchyma.

## Discussion

Our spatial analysis of 106 kidney biopsies reveals AIN as a spatially organized inflammatory disease driven by distinct cellular circuits that distinguish it from non-immune mediated kidney injury. Three major discoveries emerge: First, *VCAM1*^+^ PT cells are central to inflammatory microenvironments by providing local complement production to recruit innate immune cells and supporting metabolic-inflammatory coupling via *NAMPT*. Second, the CXCL9-CXCR3 axis generates homotypic T cell amplification networks dominated by *CD8*^+^ T cells. Third, lymphoid aggregates with TLS features compartmentalize these CXCL9-CXCR3 circuits into organized units that lack normal checkpoint regulatory elements. These discoveries provide therapeutic targets urgently needed given current reliance on non-specific immunosuppression and the increasing incidence of drug-induced AIN [35].

Our identification of 6.5-fold enrichment of VCAM1^+^ PT cells specifically in AIN extends prior identification of “failed repair” PT cells as drivers of kidney disease progression [20, 36]. In AIN, *VCAM1*^+^ PT cells express protective markers (*SERPINA3*, *THY1*) [37] alongside complement components promoting inflammatory recruitment, contrasting with profibrotic signatures in ATI. *NAMPT* emerged as the strongest discriminator of *VCAM1*^+^ PT microenvironments in AIN, representing the first demonstration of its role in immune-mediated kidney injury. PT cells showed stepwise *NAMPT* upregulation paralleling injury severity, while surrounding cells demonstrated 1.9-fold higher expression—suggesting metabolic-inflammatory niches surround *VCAM1*^+^ tubules. NAMPT functions as an intracellular rate-limiting NAD+ biosynthetic enzyme and extracellular inflammatory cytokine (eNAMPT) [29]. While NAD+ biosynthetic pathway alterations are established features of AKI [38–40], SORBET analysis indicates that enriched *NAMPT* expression specifically defines the injury-to-atrophy microenvironment, potentially reflecting both tubular metabolic dysfunction with loss of fatty acid oxidation [41, 42] and high metabolic demands of infiltrating immune cells. These findings suggest that NAD+ pathway augmentation, recently shown safe and beneficial with nicotinamide for AKI prevention in cardiac surgery [43], and eNAMPT neutralization [44] could provide complementary AIN treatment strategies.

VCAM1^+^ PT cells in AIN specifically associate with lymphocytes through unique spatial relationships. Within AIN tissues themselves, only VCAM1^+^ PT cells showed enrichment of adjacent CD8^+^ T cell and mixed B and T cell aggregates, whereas KIM1^+^VCAM1^−^ (PT-INJ1) cells and healthy tubules showed no lymphocyte enrichment in the same biopsies. This selective lymphocyte tropism demonstrates that VCAM1^+^ PT cells must specifically express lymphocyte chemotactic or adhesive signals. Melchinger et al. showed VCAM1 overexpression on cultured PT cells increased lymphocyte adhesion while CRISPR-based knockout abolished TNF-α-induced adhesion [45], providing one mechanism for this T cell interaction effect.

*C3* and other complement factor expression by injured tubular cells extends work demonstrating local kidney complement synthesis [46, 47] but reveals disease specificity: AIN shows 2-3 fold higher *C3* expression across all segments versus ATI, with injured PT and particularly *VCAM1*^+^ PT cells showing the highest expression. This hierarchy creates complement gradients orchestrating spatial immune organization, with a strong correlation between tubular *C3* and immune infiltration consistent with complement-mediated chemotaxis [48, 49]. The predominance of M1 antigen-presenting macrophages in tubular C3-C3AR1 interactions enables them to serve dual roles of presenting tubular antigens and perpetuating injury through inflammatory mediators. The spatial segregation of macrophage functions, with *C3AR1*-responsive cells at tubular interfaces versus *CXCL9*-producing populations predominating in the interstitium, creates distinct inflammatory microdomains that sustain both direct tubular damage and broader tissue inflammation.

The marked enrichment of CXCL9-CXCR3 interactions in AIN provides the strongest signature of primary immune-mediated injury versus secondary immune responses in ATI. We show that *CD8*^+^ T cells match macrophages as *CXCL9* producers, expanding the prior macrophage-centric paradigm of CXCL9 expression [50, 51]. While CXCL9-CXCR3 interactions provide the molecular signature of AIN, organized lymphoid aggregates provide the spatial compartmentalization of that signature, forming in 87.5% of AIN versus 28.6% of ATI cases. These aggregates create specialized microenvironments that concentrate inflammatory signaling. The 3.6-fold CXCL9-CXCR3 enrichment within aggregate regions identifies them as inflammatory amplification centers, sustaining immune responses independent of secondary lymphoid organs.

These lymphoid aggregates share features with tertiary lymphoid structures (TLS), including organized B and T cell zones, canonical organizing chemokines (*CCL19/CCL21*), and local antibody production. However, they lack classical stromal infrastructure—high endothelial venules, follicular dendritic cells, and fibroblastic reticular cells—with *CXCL13* expression instead deriving from immune cells themselves, suggesting self-organizing inflammatory circuits. Similar structures have been characterized in other acute inflammatory conditions [52–54], reflecting a continuum of lymphoid organization where structures lacking full stromal specialization still coordinate meaningful immune responses.

The lack of specialized stromal infrastructure could limit aggregate persistence, potentially allowing resolution once the inflammatory stimulus is removed. While these aggregates were consistent throughout our AIN cohort, their functional significance for clinical outcomes remains unclear. Given that 20-40% of AIN patients progress to CKD despite treatment [5–10], understanding what determines aggregate resolution versus persistence—and whether specific organizational features predict treatment response—represents an important area for future investigation.

Multiple regulatory checkpoints that normally suppress inflammation fail in AIN. Despite 8.7-fold higher *CD4*^+^ regulatory T cells versus ATI, these comprised only 2.3% of total T cells, suggesting inadequate compensation for the massive lymphocyte infiltration. Checkpoint-high *CD8*^+^ TRM cells co-expressing *LAG3*, *TIGIT*, and *PDCD1*—present in all reference kidneys—were absent in 88% of AIN cases. The loss of this checkpoint-high population may remove a brake on tissue inflammation, as exhausted T cells can occupy tissue niches while maintaining functionally restrained states, and in their absence more active memory populations may contribute to persistent inflammation through unchecked effector responses.

Finally, the apparent paradox that macrophage infiltration predicts functional recovery in steroid-treated AIN but progression in ATI reflects fundamentally different innate programs in response to primary versus secondary immune injury, as well as the steroid-responsive nature of AIN versus the steroid-refractory nature of ATI. Samples were obtained at similar timepoints after AKI initiation (6 days for AIN, 8 for ATI), yet M1/M2 distribution was nearly equal in AIN and heavily M2-skewed in ATI. This M2 predominance in ATI is well described in mouse models [21, 55] and correlates with eventual matrix deposition and fibrosis that steroids cannot reverse [56–58]. Critically, the association between CD206^−^ macrophage abundance and better functional recovery was observed only in steroid-treated AIN patients, suggesting that M1-skewed infiltration marks a steroid-responsive inflammatory state or that these macrophages participate in repair programs activated by immunosuppression. M1 macrophage presence may indicate reversible inflammation where tubular survival pathways (e.g., THY1, autophagy) remain intact and can be leveraged therapeutically, whereas the M2-dominated response in ATI represents a trajectory toward fibrosis refractory to immunosuppressive intervention.

Strengths of our study include a large cohort size (106 tissues), use of two complementary spatial technologies, a validation cohort from a separate institution, and comparison with both ATI and reference tissues. However, important limitations merit consideration. The cross-sectional design prevents determination of causality—we cannot establish whether VCAM1^+^ PT cells drive inflammation or result from it, nor can we track evolution of cellular states over time. The 77% steroid exposure in our AIN cohort confounds outcome associations. Limited reference cortex samples for ST (n=2) preclude formal statistical comparisons for some analyses and the heterogeneous etiologies of AIN cases (drug-induced, checkpoint inhibitor-associated, idiopathic) prevent evaluation of subtype-specific mechanisms, though consistent spatial patterns suggest shared pathophysiology.

Multiple potentially clinically relevant targetable nodes emerge from our findings, including CXCR3 antagonism [59], complement inhibition [60, 61], and NAD+ pathway augmentation [41, 43] or eNAMPT inhibition [44]. Recent studies demonstrating distinct metabolic programs in adaptive versus maladaptive tubular repair [62] reinforce the importance of identifying transition points for therapeutic intervention. Future priorities include determining how distinct AIN triggers impact these pathogenic mechanisms, longitudinal sampling if possible to capture disease evolution and identify when reversible inflammation transitions to autonomous injury, and ultimately clinical trials stratifying patients by pathway biomarkers rather than clinical AIN diagnosis alone.

## Online Methods

### Study approval and participants

For the discovery cohort, we included participants of the Yale kidney biobank enrolled between 2015-2018, which prospectively enrolled participants who underwent clinically indicated kidney biopsies at Yale New Haven Hospital and St. Raphael’s Hospital (both in New Haven, Connecticut, USA). This cohort has been described previously [33, 63]. The study excluded participants undergoing biopsy for evaluation of renal mass and allograft biopsies. All participants provided written informed consent and this study was approved by the Yale Human Investigation Committee (Protocol 11110009286).

The validation cohort of this study was approved by the Johns Hopkins Institutional Review Board under two protocols: IRB00221958 for the overall project and IRB00090103 for the use of FFPE tissue samples. Identical definitions and criteria were used to obtain clinical data.

### Establishing diagnosis and clinical cohorts

In the discovery cohort, three renal pathologists independently evaluated biopsy slides or whole slide scans from the Yale Kidney Biobank to establish AIN diagnosis for this study. The pathologists were blinded to clinical history and official biopsy report. For this study, we selected all cases that were unanimously adjudicated as AIN (n=32) or ATI (n=34), of which, 23 participants with AIN and 23 with ATI had FFPE archival tissue available for analysis and included in the discovery cohort.

For the validation cohort from Johns Hopkins University School of Medicine, cases with top-line diagnoses of AIN (n = 15) and ATI (n = 16) were identified from archival tissues spanning 2012-2024 and screened for both tissue adequacy and clinical diagnosis of AKI. These samples were analyzed using identical pathologist adjudication criteria. Identical definitions and criteria were used to obtain clinical data.

### Reference tissues

To accurately characterize the importance of findings in injured kidneys and as a quality control assessment of expected signals of resident cell antibodies for each batch, we sought to identify histopathologically normal kidney tissues for inclusion on each slide [64]. Reference tissues consisted of previously prospectively accrued living donor samples [65], obtained by back-bench biopsy at the time of organ donation, as well as highly selected prospectively obtained and banked samples of nephrectomy tissues taken from tumor-remote regions. Each kidney biopsy used as reference tissue was scored by a pathologist at 100x magnification using a raster grid consisting of a matrix of 10 rows and 10 columns of equal sized squares (100 total squares per grid). Squares overlying the renal cortex were counted for histological features of interstitial fibrosis and tubular atrophy (IFTA), interstitial cell infiltrate (ICI) and acute tubular injury (ATI). Tubular injury was defined by the presence of 3 features: loss of brush border, tubular lumen dilatation and apical blebbing. Squares containing each type of injury were divided by the total number of squares reviewed and converted into a scoring scheme as follows: none (<5% of area involved), mild (6-25%), moderate (26-50%), and severe (>50%). The overwhelming majority of reference samples used scored as “less than five percent” in all categories and were therefore included in our analysis (**Supplementary Table 1**).

For the validation cohort, reference tissues were obtained from tumor-remote regions of nephrectomy specimens performed in 2024 and underwent the same histopathological scoring protocol used for the Yale cohort to ensure minimal baseline pathology (**Supplementary Table 2**).

### Clinical data collection

In discovery cohort, clinical data were collected in the parent protocol through review of individual patient chart review using the Epic electronic health record system and stored in secured web-based data entry system (**Extended Data Fig 1a,b**). We collected over 200 variables including demographics, medical history, medications, vital signs, and laboratory data. Baseline serum creatinine values were defined based on 2 physician chart review of serum creatinine trend over the 24-month period prior to onset of the AKI event prompting kidney biopsy. Where only one value was available, this value was used as baseline. Baseline serum creatinine values were unable to be established in 8 of the 43 patients with AKI in the discovery cohort and 2 of the 31 patients with AKI in the validation cohort due to a lack of serum creatinine measurements within the defined period prior to AKI.

Onset of AKI was defined as an absolute increase in serum creatinine of 0.3 mg/dL or more (within 48 hours if occurred during an inpatient stay, no time limitation if outpatient), or a 50% or greater increase in serum creatinine (within 7 days if occurred during an inpatient stay, no time limitation if outpatient). Follow up serum creatinine measurements at 6 months were taken from values obtained from 4 to 40 weeks after the date of biopsy, with the value closest to the 6-month time point being the value that was used. There was no significant difference between the length of time from the biopsy to follow up serum creatinine measurement between the two groups (discovery cohort mean 173 days for AIN, 162 days for ATI, p = 0.74; validation cohort mean 163 days for AIN, mean 155 days for ATI, p = 0.67). Recovery of kidney function was calculated as percentage of baseline estimated glomerular filtration rate restored at 6 months using the CKD-Epi equation. Serologic and urinary data associated with individual biopsies were available through the Yale Kidney Biobank for the discovery cohort.

### Sample size determination and blinding

Our power calculation indicated that we needed 29 cases with AIN to detect a 1.75-fold difference in cell numbers between AIN and ATI with at least 90% power with alpha of 0.05. Sample sizes were determined based on tissue availability and quality requirements for spatial analysis, with power calculations indicating >80% power to detect 2-fold differences in cell densities between groups. No statistical methods were used to predetermine sample size for the discovery cohort. Three samples in the discovery cohort (1 AIN, 2 ATI) were not included in the analysis due to missing data from technical disruptions in CyTOF software causing loss of data (**Supplementary Table 1**), one sample was excluded in the validation cohort due to accidental duplication of patient sample and tissue region (**Supplementary Table 2**), and 3 reference tissues and 1 ATI tissue are missing data from ST analysis due to inadequate tissue fixation to the slide and loss of tissue during Xenium processing (**Supplementary Table 5**). Analysts performing antibody hybridization, IMC analysis, and initial clustering were blinded to diagnosis of AIN vs. ATI until after completion of cell phenotyping. For spatial transcriptomics analysis, investigators were not blinded but cell segmentation and initial clustering took place on the entire cell population aggregate, without respect to tissue type.

## Imaging Mass Cytometry

### Antibody selection, validation, and conjugation

The antibody panel used included metal-conjugated antibodies previously validated for IMC use in kidney tissue using our protocol [64, 65], as well as antibodies against new targets (**Supplementary Table 3**). New targets were selected against protein products of cell-type-specific differentially expressed genes and of signaling, state, and injury pathways from published single-cell RNA sequencing datasets. Only BSA-free and carrier-free commercial antibodies against selected targets were suitable for inclusion to be compatible with heavy metal conjugation, and monoclonal antibodies with supplier knock-out validation were given preference in our testing.

Antibody validation and IMC immunolabeling were performed as previously described [64]. Antibodies were validated using tissues in which target antigen would be expected to be found. For example, antibodies against immune cells were validated in a combination of human tonsil and chronic severe interstitial nephritis tissues. Antibodies against kidney-resident cells were validated in normal kidney tissues. Antibodies in which positive staining was detected in expected cells with a maximal signal-to-noise ratio >3 and in which significant ectopic or background staining was not observed in immunofluorescence staining underwent heavy metal conjugation. These antibodies were conjugated using the Maxpar conjugation kit (Standard BioTools) following the manufacturer’s protocol, scaled for 100 μg of carrier-free antibody. Metals were selected based on expected strength and to minimize overlap due to isotopic impurity. Newly conjugated antibodies were validated using IMC along with the full panel of previously validated antibodies, to ensure both that binding specificity had not changed through the conjugation process, and that the addition to the panel did not interfere with the signal of previously validated antibodies. The final panel consisted of 31 antibodies targeting structural markers, immune cells, injury markers, and functional states.

### Slide generation and antibody hybridization

Each slide contained one AIN, one ATI, one reference healthy kidney tissue, and one lymph node, which collectively constituted one “batch” (**Supplementary Tables 1,2**). While including one healthy reference kidney tissue on each slide allowed for a quality assessment of antibody targets of resident kidney cells, inclusion of lymph node on each slide allowed for a quality assessment of immune cell markers.

For each biopsy, the first 3-μm thick section was stained for H&E and an adjacent 5-μm thick unstained section was used for IMC. H&E slides underwent Aperio scanning for quality control of each biopsy and the images were analyzed on Aperio ImageScope (v12.4.6). Unstained tissue slides allocated for IMC were labeled using a cocktail of metal-conjugated antibodies within 2 weeks of tissue fixation to slide to minimize tissue oxidation. For IMC, the unstained tissue sections were deparaffinized, rehydrated, antigen retrieved at pH 9 and then hybridized with a cocktail of metal-conjugated antibodies. Staining was performed within 2 weeks of sectioning to minimize tissue oxidation. By design, all tissues on a given slide underwent antibody hybridization and IMC ablation at the same time. After antibody hybridization, tissue slides were then ablated and imaged using the ND:YAG 213nm laser in the Hyperion™ Imaging System (Standard BioTools) with 1-μm pixel size, with each aerosolized plume analyzed in a Standard BioTools CyTOF mass cytometer, within 4 days of antibody hybridization to minimize deterioration of antibody signal and tissue oxidation.

For the validation cohort, tissue sections from FFPE tissue blocks were processed at John Hopkins University (under supervision of author AR), and shipped overnight to our site for further IMC analysis as described and underwent antibody hybridization and IMC using the same hybridized antibodies, identical protocols and tools, and by the same author (MLB) prior to undergoing IMC at the CyTOF Core at Yale.

### Image acquisition

The stained sections were ablated using the ND:YAG 213nm laser in the Hyperion™ Imaging System (Standard BioTools) with 1 μm^2^ resolution, and the released metals were analyzed in a tandem CyTOF mass cytometer (Standard BioTools). Regions of interest for ablation were selected to span the largest tissue areas possible and to ablate cortex and medulla separately, covering the entire AIN and ATI tissue biopsy. Larger regions of interest were ablated for all reference tissues where additional tissue was available. For tumor-remote nephrectomy reference samples reconstructed into tissue microarrays and for lymph node samples, a region of interest of 1500 μm by 700 μm was selected.

### Image quality control and generation

Images were initially viewed and antibody hybridization was validated using MCD Viewer (Standard BioTools, v1.0.560.6). For all images shown, thresholding was performed in MCD Viewer with gamma set to 1, minimum color range set to zero, and maximum color range set to 100. Threshold minimum was set within the range of 0 to 2 to remove background signal, as assessed morphologically. For all comparative images, signal thresholds (minimum and maximum) were set identically for each marker between samples to allow for unbiased comparison. Analyses of cortex and medulla were performed separately, with corticomedullary junction being analyzed with cortical tissues. Cortex and medulla were defined morphologically.

### Pre-processing

IMC images were pre-processed using a python-based algorithm ‘Steinbock’ as described [66]. Pre-processing included configuring the antibody panel and a hot pixel-filtering step for raw image files. Single cell segmentation was performed using custom optimized Mesmer [67] as described below. Generated segmentation masks were used within steinbock to measure cell intensities, cell morphology, cell neighbors and ultimately export single-cell data for downstream analysis.

### Object Segmentation

Cell segmentation was augmented by implementing the Maxpar® IMC™ Cell Segmentation kit in the antibody panel and pixel expansion method on Mesmer [67], a deep-learning-enabled segmentation algorithm. While trained on a dataset containing more than 1 million manually labeled cells, Mesmer algorithm default parameters were modified for optimal cell segmentation of kidney tissue. Specifically, the modifications included resolution from 1 to 0.8, small object threshold modified from 15 to 53, maxima threshold modified from 0.1 to 0.3, and interior threshold modified from 0.2 to 0.47. Cell segmentation was performed by expansion method by expanding by 1 µm around the nucleus. Performance of optimized cell segmentation method was validated using “the-segmentation-game” plugin on napari, a fast, interactive viewer for multi-dimensional images [68]. F1-score [69] was determined on 15 different tissue samples for which segmentation masks were generated using custom modified Mesmer parameters: 3 chronic kidney disease samples, 3 acute kidney injury samples, 3 diabetic kidney disease samples, 3 tumor remote nephrectomy reference samples, and 3 living donor biopsy reference samples. Across all these samples the mean F1-score was 0.838 with a standard deviation of 0.0274 (**Supplementary Table 8**).

### Signal spillover correction

While the use of mass spectroscopy of rare earth metals conjugated to antibodies to distinguish unique antibody binding greatly decreases the extent of crosstalk between channels relative to dye-based methods, spillover still exists in mass cytometry data. We employed a previously described bead-based compensation workflow and R-based software [70] to estimate and compensate for interference between channels.

### Image and cell-level quality control and batch effect correction

All downstream analysis was performed within the imcRtools workflow [66]. We performed image-level quality control through calculating the signal-to-noise ratio for individual channels and found that each of our channels had an acceptable ratio >3 using the otsu thresholding approach. Batch effect correction was performed between different batches (slides) using the fast mutual nearest neighbors function within the Batchelor package [71].

### Cell phenotyping

The resultant low dimensional embedding of integrated cells after batch effect correction was used for unsupervised cell phenotyping. We employed the RPhenoGraph clustering approach, which includes calculation of a graph by detecting the K nearest neighbors based on Euclidean distance in expression space, weighting of edges between nodes by overlap in nearest neighbor sets using the jaccard index, and Louvain modularity optimization to detect connected communities and partition the graph into clusters of cells [72]. Initial unsupervised clustering results were analyzed through mapping back onto original tissues to assess cell locations as well as analysis of marker expression using violin plots of raw expression patterns. Subclustering was attempted on all unsupervised clusters using all markers appearing within the cluster to be expressed more highly than the average across all unsupervised clusters. Ultimate cell annotations were validated through marker expression patterns and spatial localization, identifying 49 detailed cell populations (including mixed populations, **Supplementary Table 4**), which were distilled to 18 core cell populations and two core mixed cell populations (**Fig. 2c**). All core cell populations identifiable through our RPhenoGraph clustering and subclustering approach were reproduced in the validation cohort (**Extended Data Fig. 1d**)

### Spatial neighborhood analysis

Cell-cell interactions allow cells to exist in communities, perform collective functions, and respond to tissue-level stimuli in a coordinated fashion. Using a k-nearest neighbors-based approach calibrated to capture only the direct, immediate neighboring cells of each cell across different kidney tissues, we were able to define the frequency at which defined interstitial cell types were in immediate spatial proximity to defined tubular cell types. We were interested in capturing each cell’s immediately adjacent neighboring cells, the cells most likely to be directly interacting with a given cell. Using 2 living donor biopsy tissues, 1 AKI biopsy tissue, and 1 CKD biopsy tissue, we manually visually counted the number of immediately adjacent neighboring cells for a subset of 100 random cells per tissue. Across 400 cells and 3 tissue types, we found that individual cells in our kidney tissues have 5 immediately adjacent neighboring cells, on average, with a mean centroid-to-centroid distance of 16 µm. Using a k-nearest neighbors approach and a threshold of 5, we generated a spatial interaction graph which was used to determine the identity of each cell’s immediately adjacent neighbors across the entire project. For tubular-interstitial interaction analysis, we used tubular epithelial clusters as index cell types and immune, endothelial, vascular, and stromal clusters as neighboring cell types.

### Statistical analysis of cell densities and spatial neighbors for IMC data

Cell type densities (cells/mm²) in IMC data were calculated by dividing total cell count by tissue area. Tissue area was measured manually using Fiji/ImageJ. Densities were compared between AIN, ATI, and reference tissue groups using pairwise Wilcoxon rank-sum tests with Bonferroni correction for multiple comparisons. Proportional distributions were compared using Fisher’s exact tests followed by Bonferroni correction.

For analysis testing the association of diagnosis with number of index and surrounding cell pairs, we allowed each type of cell to be surrounded by up to 5 nearest neighbors. To test the association of each diagnosis with index cell-surrounding cell pair, we fit a generalized linear mixed model with outcome as number of index cell surrounding cell pairs and predictor as diagnosis using log link. We clustered the analysis at the level of participant. We present fold difference in number of cells between various diagnoses (β coefficients) as well as multiple comparison adjusted q-values using Benjamini Hochberg procedure (**Fig. 2f; Extended Data Fig. 5a).**

To examine whether different PT injury states within AIN tissues exhibited distinct patterns of immune cell recruitment, we analyzed the immediate cell neighbors of PT-H, PT-INJ1, and PT-INJ2 cells exclusively within AIN cortex samples (**Fig. 2g; Extended Data Fig. 5b**). For each PT cell, we identified up to 5 nearest neighbors. We fit a generalized linear mixed model with log link for each surrounding cell type, with the number of index-surrounding cell pairs as the outcome and PT cell type as the predictor, using PT-H as the reference category. These analyses were conducted in Stata Statistical Software: Release 18, College Station, TX: StataCorp LLC.

## Spatial transcriptomics

### Tissue cohort and gene panel

Single cell spatial transcriptomic analysis was performed on 20 patients included in the Yale cohort IMC analysis. This included 18 unique cortex tissue samples, 4 unique medulla tissue samples, 8 unique AIN patient samples, 7 unique ATI patient samples, and 5 unique healthy reference living donor biopsy samples (**Supplementary Table 5**). 10x Genomics Xenium Analyzer was chosen as the platform to perform spatial transcriptomic analysis because of its single cell resolution, relatively high sensitivity without sacrificing specificity in FFPE tissues [73, 74], and multimodal cell segmentation algorithm. The analysis employed the Xenium Prime 5K Human Pan Tissue & Pathways Panel (5,001 genes) plus a custom 100-gene panel (**Supplementary Table 9**) targeting kidney-resident cells, immune cells, and cell state-specific markers. The custom panel included genes targeting: kidney tubular segment markers (*SLC12A1*, *SLC12A3*, *AQP2*, *SLC34A1*, *SLC22A6*), immune-related markers (*PTPRC*, *CD74*, *CD69*, *CCL2*), injury markers (*LCN2*, *SPP1*), proliferation markers (*PCLAF*, *PRC1*), and fibrosis markers (*COL1A1*, *COL1A2*, *VIM*, *TAGLN*).

### Cell segmentation

Cell segmentation was performed using Xenium’s multimodal algorithm, which employs deep learning to analyze nuclear, cell interior, and cell boundary staining patterns. To validate segmentation accuracy and spatial fidelity, we compared the ST results with post-workflow H&E staining and cell segmentation target images of the same tissue sections. Visual inspection using Xenium Explorer confirmed accurate transcript localization for cell-specific markers (e.g., *LRP2* in PT cells) and proper cell boundary identification across all samples.

### Data integration and clustering

Single-cell spatial transcriptomics data were processed using Seurat v5. Following quality control to remove cells expressing fewer than 10 unique genes, we created a unified Seurat object containing patient identifiers, region, and category information. After quality control, our dataset had a median of 139 (IQR: 53-374) transcripts per cell and a median cell diameter of 8.33 μm. Data layers were split by patient groups to enable sample-specific processing before integration. Expression matrices underwent SCTransform normalization, which uses regularized negative binomial regression to simultaneously normalize the data and identify variable features while accounting for technical variation in sequencing depth. Gene expression values were scaled to standardize the mean expression and variance across cells. To address batch effects while preserving biological variation, we applied Harmony integration through Seurat’s IntegrateLayers function, creating a shared embedding space for cross-patient comparison. Shared nearest neighbor graphs were constructed using the first 30 Harmony dimensions, followed by Louvain clustering. Optimal clustering resolution was determined by evaluating silhouette scores across multiple parameters (0.4-1.2), with resolution 0.8 providing the highest score and optimal biological separation. Cell type annotation combined differential gene expression analysis with spatial localization patterns and histological correlation, particularly for thick ascending limb and parietal cells, resulting in identification of 26 core cell types and one mixed population (**Supplementary Table 6**).

To capture cellular heterogeneity, we performed targeted subclustering of T cells (**Extended Data Fig. 3c,d**) using lineage markers (*CD4, CD8A, FOXP3, CCR7, CD69*) and macrophages using inflammatory (*IL1B, TNF, CXCL8*) and regulatory (*MRC1, MSR1, CD163*) signatures. Proximal tubule cells underwent subclustering to identify healthy (PT-H; *LRP2*^+^, *CUBN*^+^), injured (PT-INJ; *HAVCR1*^+^, *VCAM1*^+^, *LCN2*^+^), and dedifferentiated (PT-dd; reduced canonical and injury marker expression) states based on established differentiation and injury markers (**Supplementary Table 7**).

### Statistical analysis of cell densities and spatial neighbors for spatial transcriptomic data

Cell type densities (cells/mm²) in spatial transcriptomic data were calculated by dividing total cell count by tissue area, measured using the shoelace formula after boundary delineation in R Shiny v1.7.4 with Plotly v4.10.1. Cell inclusion was determined by ray-casting algorithm. Densities were compared between AIN, ATI, and reference tissue groups using pairwise Wilcoxon rank-sum tests with Bonferroni correction for multiple comparisons. Proportional distributions were compared using Fisher’s exact tests followed by Bonferroni correction. Cell type annotations were validated by comparing densities between ST and IMC data from 20 overlapping patient samples (**Supplementary Fig. 5**). Equivalent cell types were mapped between platforms, encompassing tubular epithelial, endothelial, stromal, and immune cell populations. Cell type proportions were consistent between IMC and ST platforms.

### Predicted ligand-receptor interaction analysis

To validate molecular communication mechanisms identified by IMC, we performed spatially-constrained ligand-receptor interaction analysis on ST data. We analyzed two key ligand-receptor pairs based on differential expression results: CXCL9-CXCR3 and C3-C3AR1. Cells were classified as ligand- or receptor-positive based on detectable expression (>0 normalized counts) of respective genes. We quantified spatial interactions by calculating Euclidean distances between all ligand-expressing and receptor-expressing cells using transformed spatial coordinates. Cell pairs within 20 μm were considered interacting, consistent with our k-nearest neighbor analysis showing mean intercellular distances of 16 μm of immediately adjacent neighbors. Possible autocrine interactions, or expression of receptor and ligand within the same cell, were not included in these analyses.

Interaction densities (interactions/mm²) were calculated by normalizing total interactions to tissue area. For each ligand-receptor pair and disease category, we characterized interaction composition by identifying participating cell types, classified as “ligand_cell_type → receptor_cell_type”. Mean minimum distances were calculated as the average distance from each ligand-positive cell to its nearest receptor-positive cell, regardless of whether that nearest cell was within the 20 μm interaction threshold, providing a measure of overall spatial clustering between cell types. Statistical comparisons used Poisson tests for interaction counts with patient-level clustering.

### Lymphoid aggregate region identification

Following established criteria [34], we identified organized lymphoid aggregates through a two-step process. First, we detected B cell aggregates (≥10 B cells within 100 μm radius) using spatial clustering with the dbscan algorithm. Second, we refined aggregate centers using density-based analysis to identify the highest concentration of B cells and plasma cells within a 250 μm search radius. For each refined center, we evaluated cellular composition within a 150 μm radius, classifying regions as organized lymphoid aggregates if they contained: (1) B cells (≥10), (2) T cells (≥5), (3) dendritic cells (≥2) or plasma cells (≥5), and (4) expression of lymphoid organizing chemokines (*CXCL13*, *CCL19*, or *CCL21*). This density-based refinement ensured accurate capture of lymphoid aggregate cores while maintaining standardized criteria for comparison across samples.

To characterize the organizational features of these aggregates, we assessed markers of specialized lymphoid stromal cells found in classical tertiary lymphoid structures. Specifically, we evaluated: high endothelial venule markers (*GLYCAM1*, *MADCAM1*), follicular dendritic cell markers (*CR2*), and fibroblastic reticular cell markers (co-expression of *PDPN* and *CCL19*). We did not identify cells expressing these specialized markers within the aggregates. Additionally, we quantified cellular sources of *CXCL13* expression and found it was predominantly derived from immune cells (B cells: 289/793, T cells: 257/793 *CXCL13*^+^ cells) with minimal stromal contribution (fibroblasts: 13/793). Despite lacking these specialized stromal components typically associated with mature tertiary lymphoid structures, these aggregates met organizational criteria used in recent studies including Liu et al. [34], who similarly characterized structures lacking classical FDCs as TLS in nasopharyngeal carcinoma.

### SORBET Analysis

We adapted Spatial Omics Framework for Biomarker Extraction and Tissue classification (SORBET) [31] — a supervised, geometric deep learning model — to predict PT cell state from ST data. SORBET requires two inputs: tissue samples profiled with spatial ‘omics technologies and a label (PT class) associated with each subregion (subgraph). Analyses were restricted to the renal cortex and only considered the AIN and ATI pathologies. AIN and ATI were modeled separately.

PT cells were classified as PT^neg^ (*HAVCR1*^−^*VCAM1*^−^), *HAVCR1*^+^*VCAM1*^−^ PT or *VCAM1*^+^ PT. A cell was considered positive for *HAVCR1* or *VCAM1* if its expression of the specified marker exceeded zero.

Each tissue sample was modeled as a graph *G_K_* = (*V_K_*, *E_K_*), where *V_K_* is the set of cells and *E_K_* a set of edges encoding spatial adjacency. A single distance threshold was set for each graph to identify edges. Briefly, let *d*_10_(*c_i_*) denote the distance from *c_I_* to its 10^th^ nearest neighbor. Then the edge \**c_i_*, *c_i_*+ ∈ *E_k_* if *d*\**c*_10_, *c*_1i_+ ≤ Median({d_10_(c_i_)|c_i_ ∈ V_k_}).

Next, cell niches were extracted around each PT cell; the niche was labeled according to that PT cell’s classification. SORBET’s automated subgraph extraction algorithm was to extract niches with the parameters *k* = 2 (niche radius / hop-distance) and *MS* = 5 (minimum subgraph size). To avoid confounding, all PT cells and their incident edges were excluded from extracted cell niches.

SORBET’s graph convolutional network (GCN)-based model was modified to account for multi-class prediction. Rather than a sigmoid function, 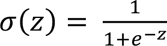, the final layer is replaced with a soft-max function, 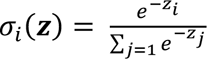, over three outputs, ***z*** ∈ ℝ^+^, corresponding to the three potential PT classes. The output loss function is modified to compute the (multi-class) cross-entropy loss, 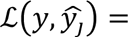 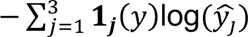, where **1**_*j*_(*y*), is the indicator that the true label is j and 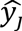 is the predicted likelihood that the sample is of class *j*. Models were otherwise trained using the same methods as described in [31]; hyperparameters were optimized to maximize the F1-score.

Models of PT cell state in the AIN and ATI populations were interpreted using cell phenotype similarity (CPS) score and canonical correlation analysis (CCA)-based analyses to identify cell- and marker-level patterns, as described in [31]. These methods were originally designed for interpreting binary classification tasks. Each disease (AIN and ATI) was analyzed in three separate comparisons: PT^neg^ vs *HAVCR1^+^VCAM1*^−^ PT, PT^neg^ vs *VCAM1*^+^ PT, and *HAVCR1^+^VCAM1*^−^ PT vs *VCAM1*^+^ PT.

For any of the above pairwise comparisons, the CPS method determines which cells are most relevant for pairwise discrimination between of the corresponding two types of PT cells in their respective neighborhood. Briefly, let *z_i_* represent the model’s representation of a cell’s spatial context and *s_j_* the representation of the *j*-th PT cell’s niche. Moreover, let *k* ∈ {*PT^neg^, HAVCR*1^+^*VCAM*1^−^ *PT, VCAM*1^+^ *PT*. Then, the score for cell *c_i_*, with model representation, *z_i_*, is

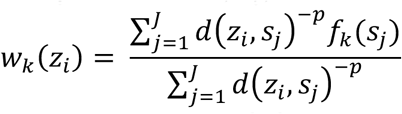

where *J* denotes the number of PT cell niches and *f_k(_ s_j_*_)_ is a function that is 1 if the *j*-th PT cell is in state *k* and -1 if it is not in the *k* state.

Similarly, we fit CCA models to the input data, *c_i_*, and the representation of the PT cell’s niche, *s_j_*. Let *c_k_* = [*c_i_*] denote a matrix of the cells input data (*C*∈ ℝ^*N* × *d*^, where *N* is the number of cells in cell niches and *d* = 5101, the number of markers profiled). Then *S*_2_ = [*s_j_*] is a matrix of niche representations (*S* ∈ ℝ^*N* × *h*^, where *h* is the size of the PT cell niche representation), where cell *i* is in PT niche *j*. Here we modify the CCA analysis in SORBET each hop distance SORBET approach, by discarding the CCA computation between each PT cell and its associated subgraph representation.

### Differential expression analyses

Differential expression analysis was performed using Seurat’s FindMarkers function with Wilcoxon rank-sum test. Three pairwise comparisons were conducted: AIN-cortex versus Ref-cortex, ATI-cortex versus Ref-cortex, and AIN-cortex versus ATI-cortex. Testing was restricted to cell types containing at least 3 cells per condition group. Parameters included minimum gene detection threshold of 10% in either group, minimum average log2 fold-change of 0.25, and Bonferroni correction for multiple testing. Genes were considered significantly differentially expressed at adjusted p-value < 0.05. For visualization, volcano plots were generated with EnhancedVolcano package showing -log10(adjusted p-value) versus log2 fold-change.

### Statistical analysis

All statistical analyses were performed in R (v.4.3.0) unless otherwise specified. Continuous variables were compared between groups using Wilcoxon rank-sum tests for two-group comparisons or Kruskal-Wallis tests for multiple groups. Categorical variables were compared using Fisher’s exact tests. Correlations employed Spearman’s method with Fisher’s z-transformation for comparing correlation coefficients between groups: z = 0.5×ln((1+r)/(1-r)) with standard error = 1/√(n-3).

For analyzing recovery outcomes, we used weighted Spearman correlation where weights were defined as the number of index tubular cells per patient. Generalized linear mixed models with log link accounted for patient-level clustering when analyzing cell-cell interactions. Multiple testing correction used Bonferroni adjustment for predefined comparisons (cell densities, spatial interactions) and Benjamini-Hochberg false discovery rate for exploratory analyses (differential expression, correlation networks).

For spatial transcriptomics analyses of cortex tissues only, reference tissue comparisons (n=2 cortex tissues) are presented as descriptive observations only, without formal statistical testing. Statistical comparisons are limited to groups with adequate sample sizes (cortex, AIN n=7, ATI n=6; cortex and medulla, AIN n=8, ATI n=7, Ref n=5).

All tests were two-sided with significance threshold of P < 0.05 after adjustment. Random seed was set for all stochastic processes to ensure reproducibility. Effect sizes were calculated using Cohen’s d for continuous variables and Cramér’s V for categorical variables. Software versions: R v.4.3.0, Seurat v5.0, Harmony v0.1.1, batchelor v1.12.3, imcRtools v1.14.0, and Python v3.9 with scanpy v1.9.3 for specific analyses.

## Data availability

Raw imaging mass cytometry and spatial transcriptomic data will be deposited to Gene Expression Omnibus upon acceptance. Analysis code will be made available at our github.com site upon publication. Clinical metadata will be available at Datadryad.org upon acceptance. Additional data supporting the findings are available from the corresponding author upon reasonable request.

## Acknowledgments

We thank the participants who contributed samples to the Yale Kidney Biobank (ClinicalTrials.gov Identifier: NCT04343417) and who contributed samples available at Johns Hopkins University, without whom this study would not have been possible. We thank the Yale Kidney Biobank and the Yale Clinical and Translational Research Accelerator (CTRA), which houses and manages the biobank, for sample collection and processing. We thank Lori A. Charette and Deirdre H. Salemme from Yale Pathology Tissue Services for assistance with tissue fixation for IMC and spatial transcriptomic analyses. We thank Ruth Montgomery (former Director), Liza Konnikova (Director), and Shelly Ren from the Yale CyTOF Core Facility for technical support. We thank William Renock, Bony De Kumar, and Ashish Shelar from the Yale Center for Genome Analysis (YCGA) for assistance with Xenium spatial transcriptomic processing and bioinformatics support. We thank the Yale Center for Research Computing (YCRC) and their staff for computational resources and technical support.

This work was supported by NIH grants R01DK126815 (LGC), R01DK128087 (DGM), T32DK007276 (MLB), U01DK133768 (VRK and LGC), P30DK045735 (Yale Diabetes Research Center Pilot and Feasibility Grant, MLB and LGC), and U54DK137331 (CRP); American Diabetes Association Postdoctoral Fellowship 11-23-PDF-63 (MLB). Development of SORBET was supported by NIH grants R01DA063148, U01DA051410, U54AG076043, U54AG079759, and P50CA121974 (JMC and YK).

## Figures

**Extended Data Figure 1.**
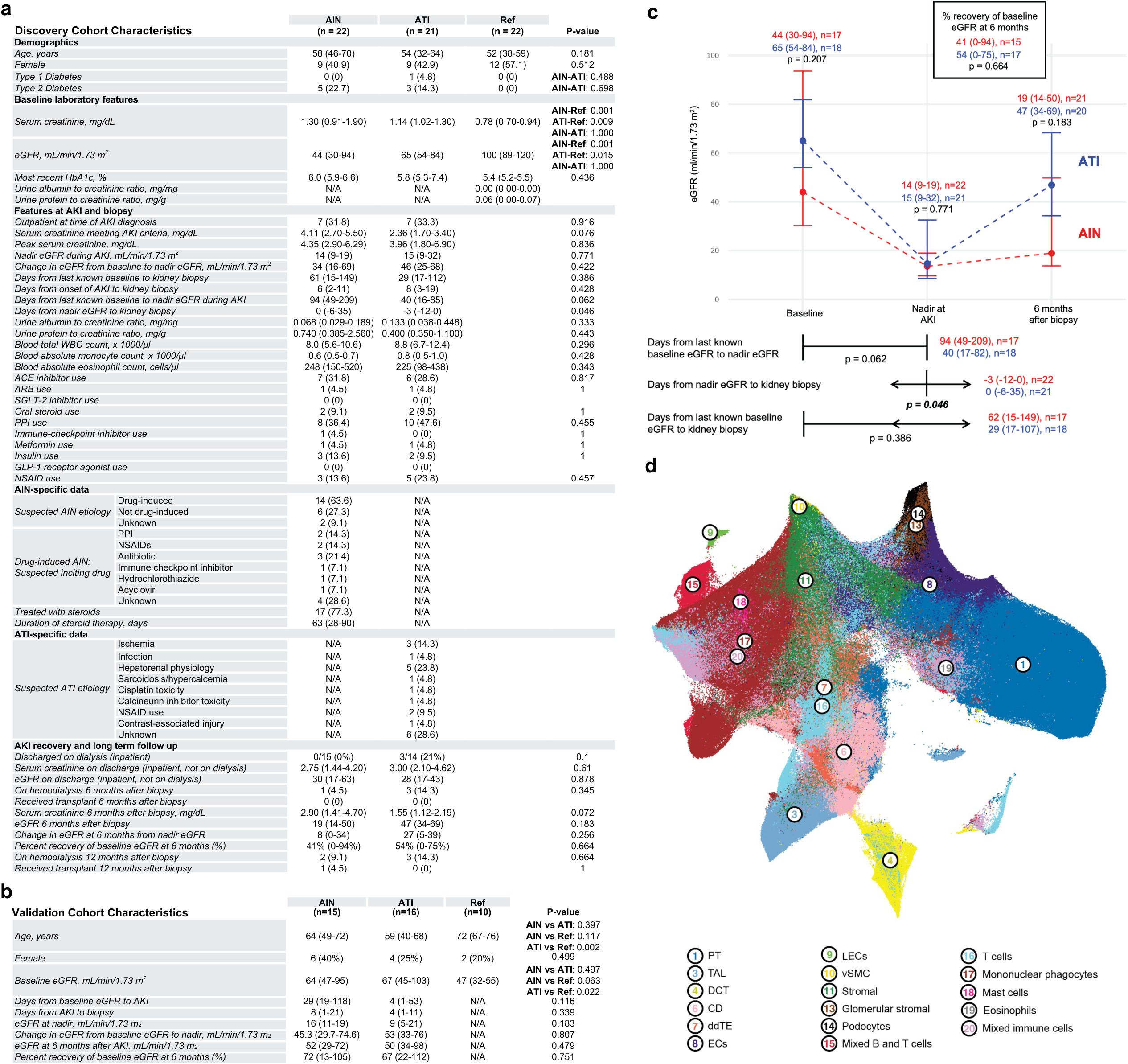
Clinical characteristics and cellular landscapes of discovery and validation cohorts in acute interstitial nephritis.

**Extended Data Figure 2.**
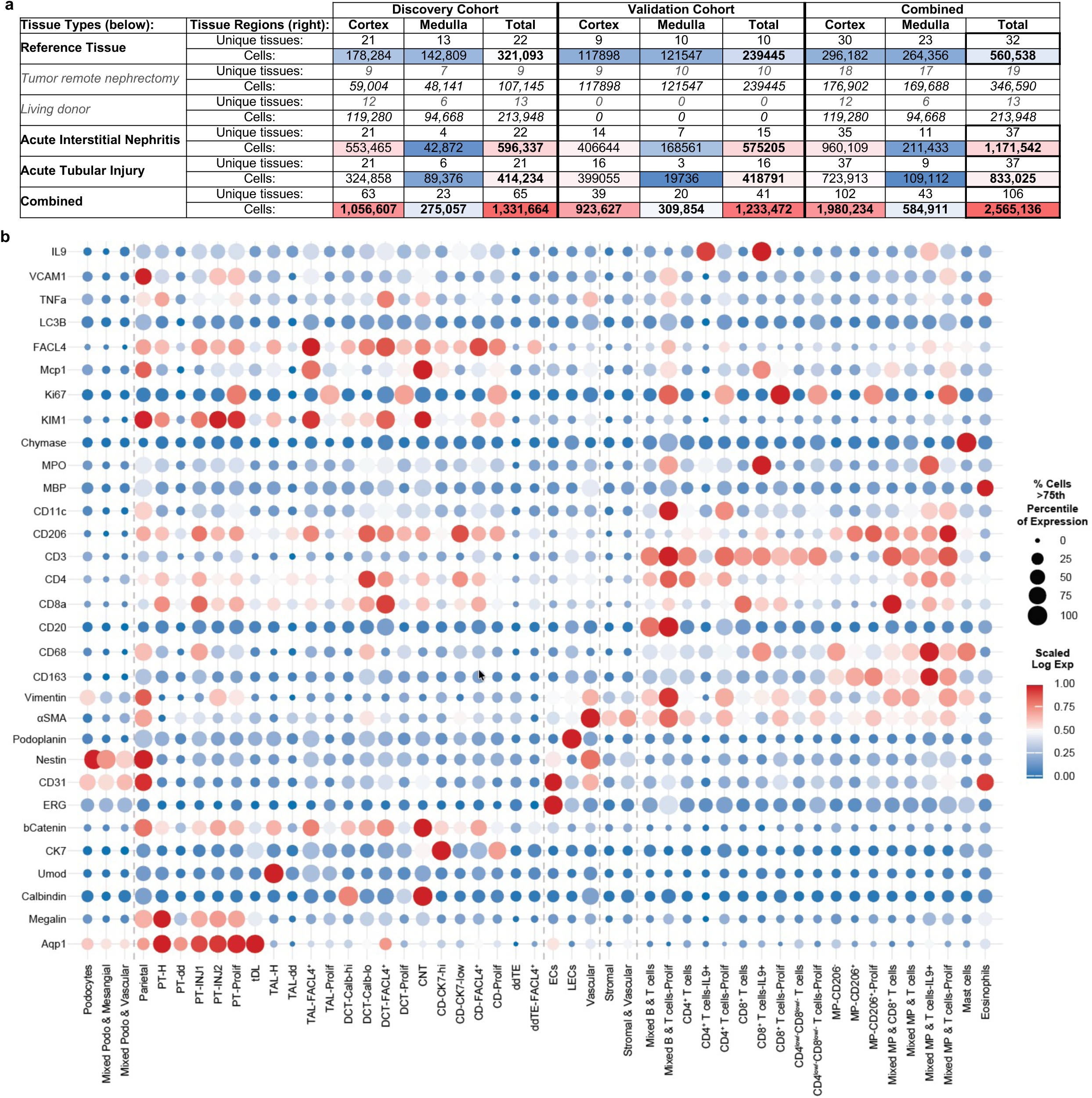
Tissue distribution and marker expression profiles of IMC-defined cell populations.

**Extended Data Figure 3.**
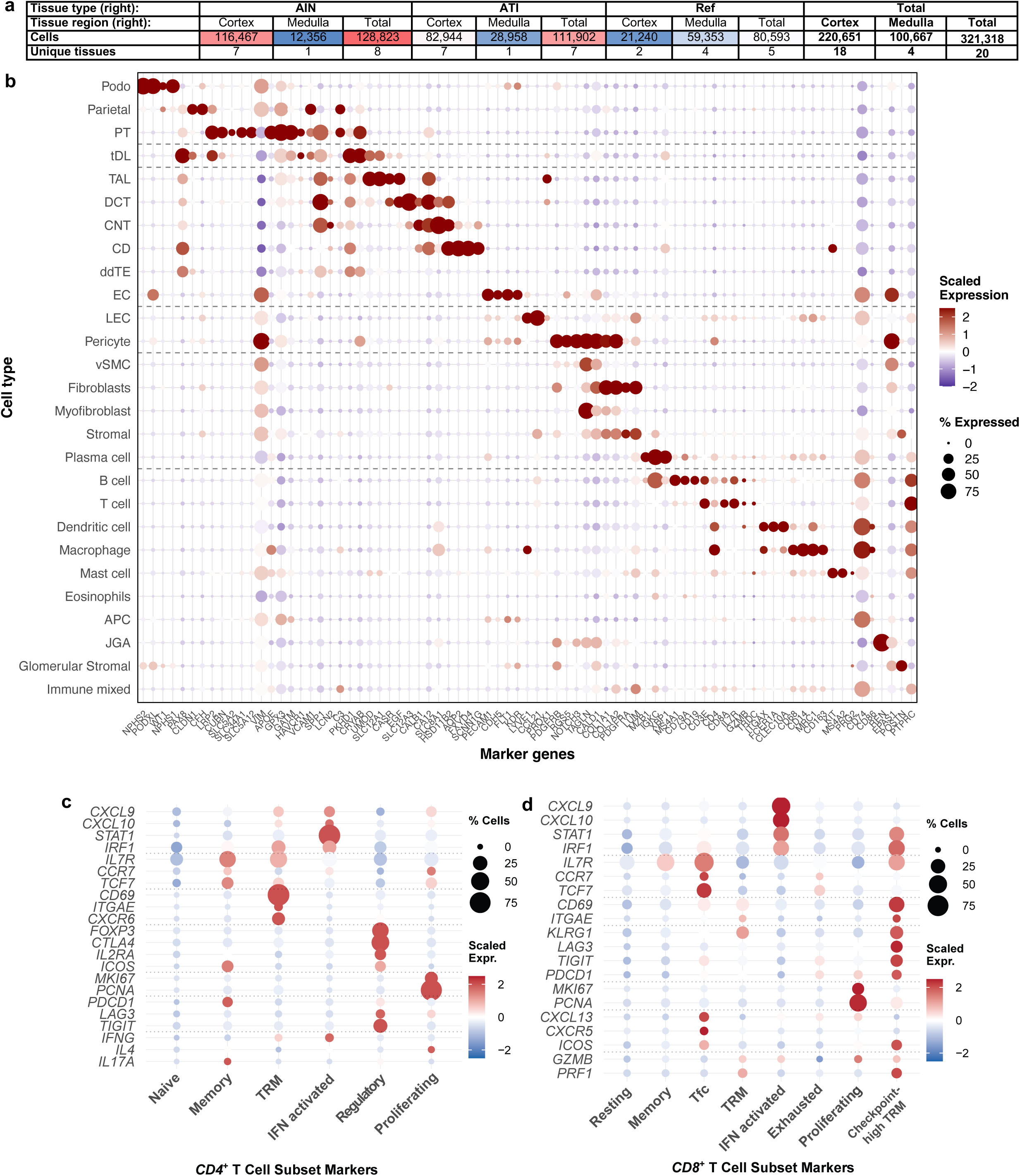
Spatial transcriptomics cellular landscapes and T cell subset characterization.

**Extended Data Figure 4.**
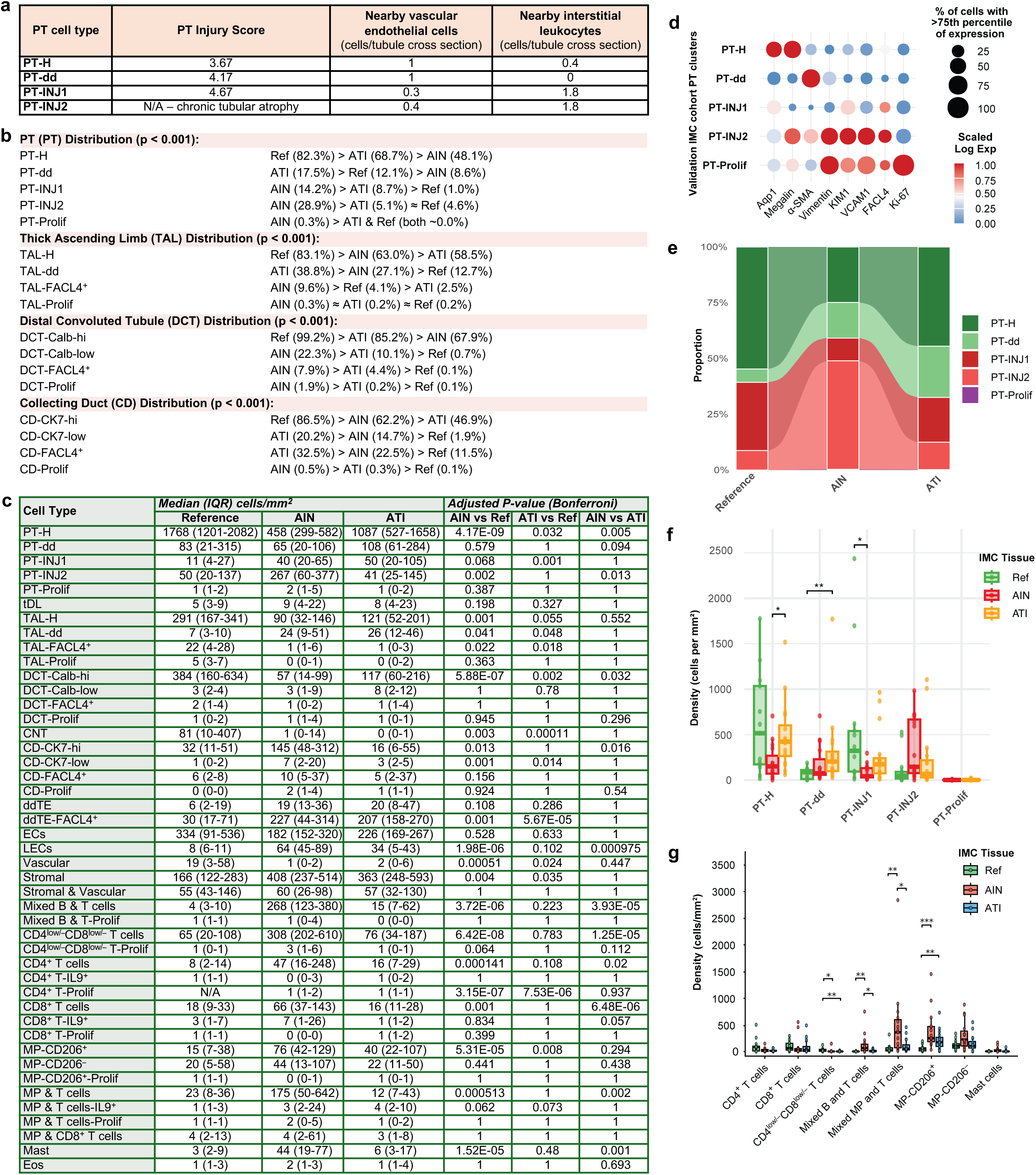
Proximal tubule injury phenotyping and cellular distribution analysis across discovery and validation IMC cohorts.

**Extended Data Figure 5.**
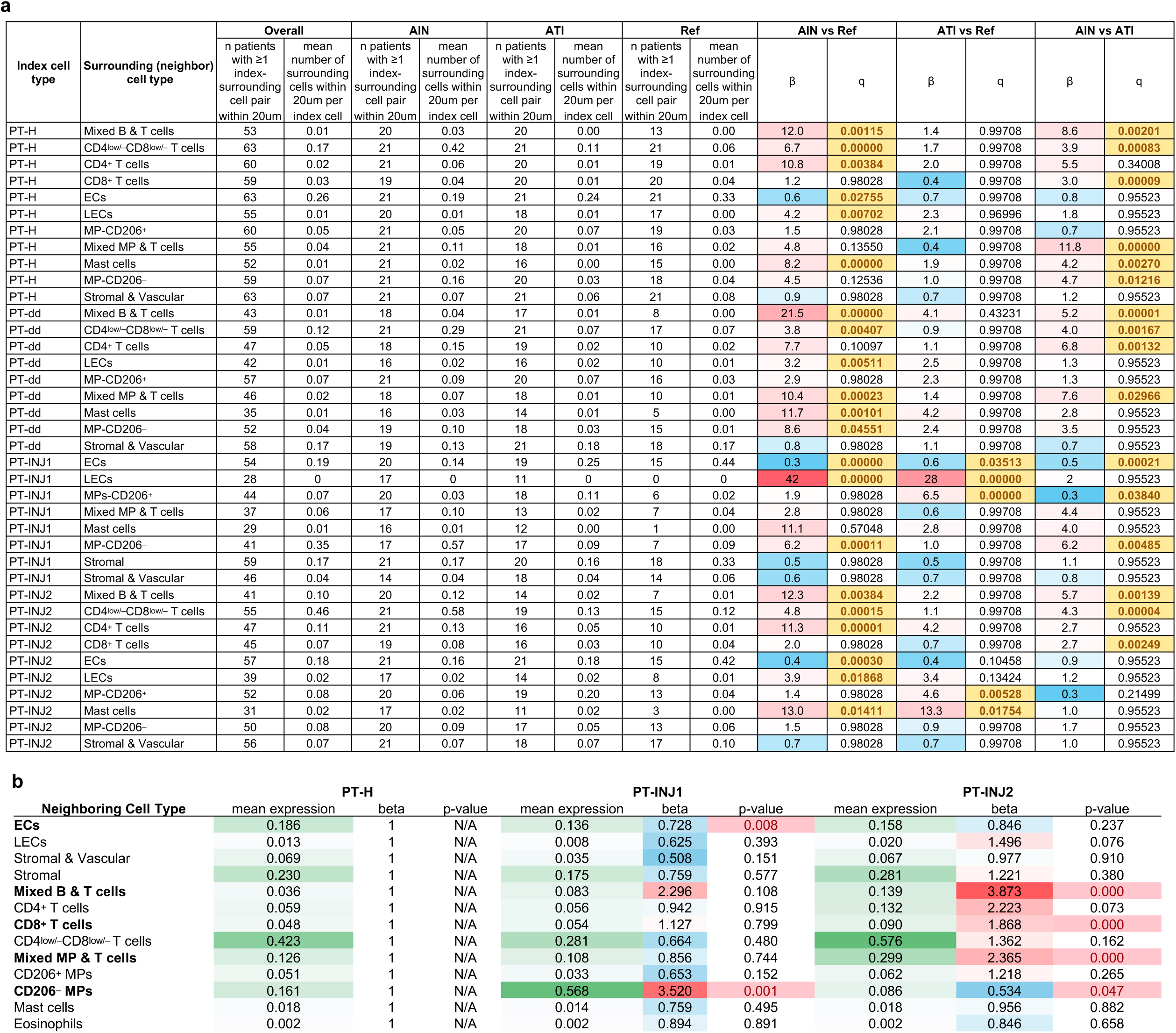
Spatial analysis of immune cell neighborhoods surrounding proximal tubule injury states.

**Extended Data Figure 6.**
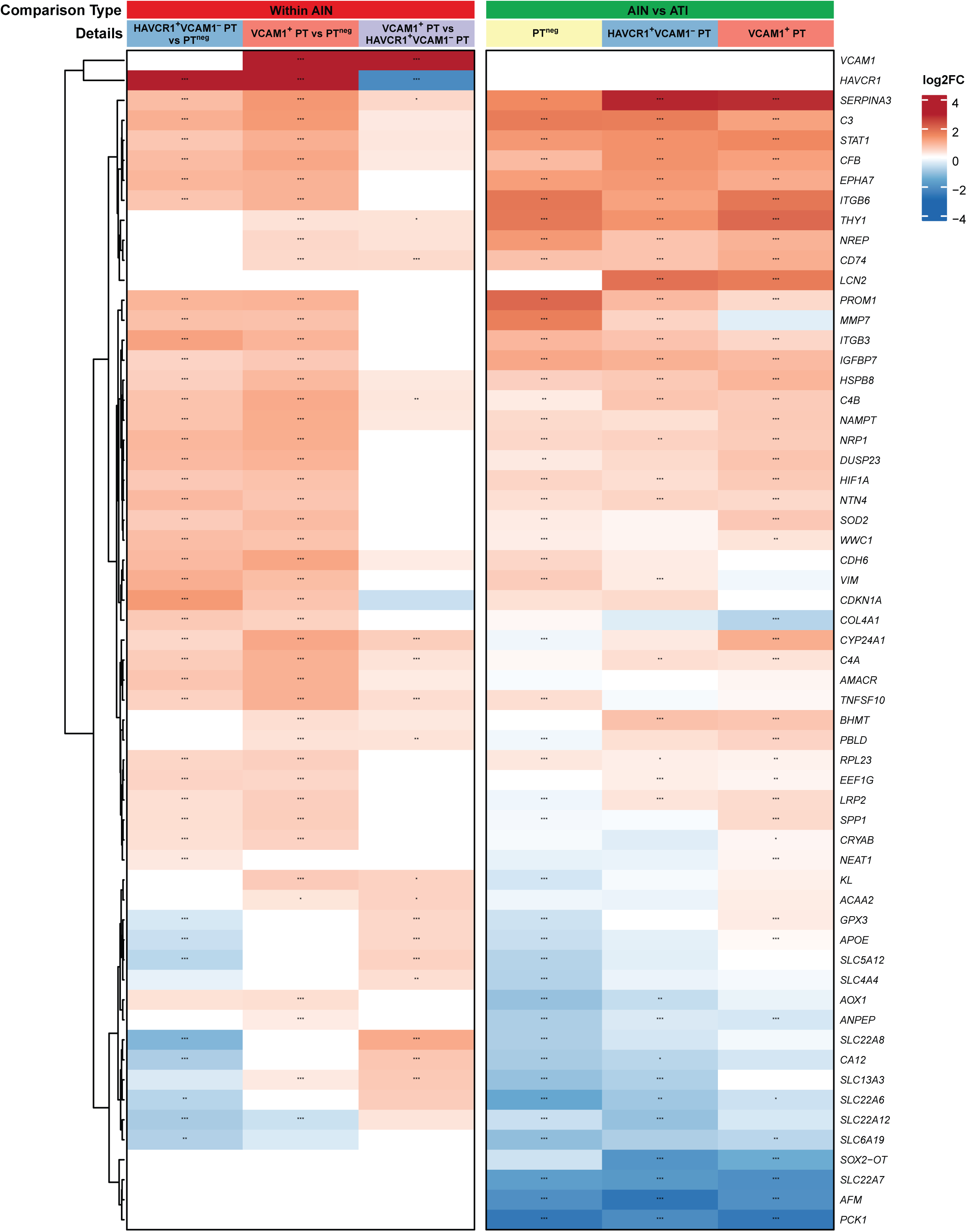
Differential gene expression analysis of proximal tubule injury states within AIN cortex and between AIN and ATI cortex
tissues.

**Extended Data Figure 7.**
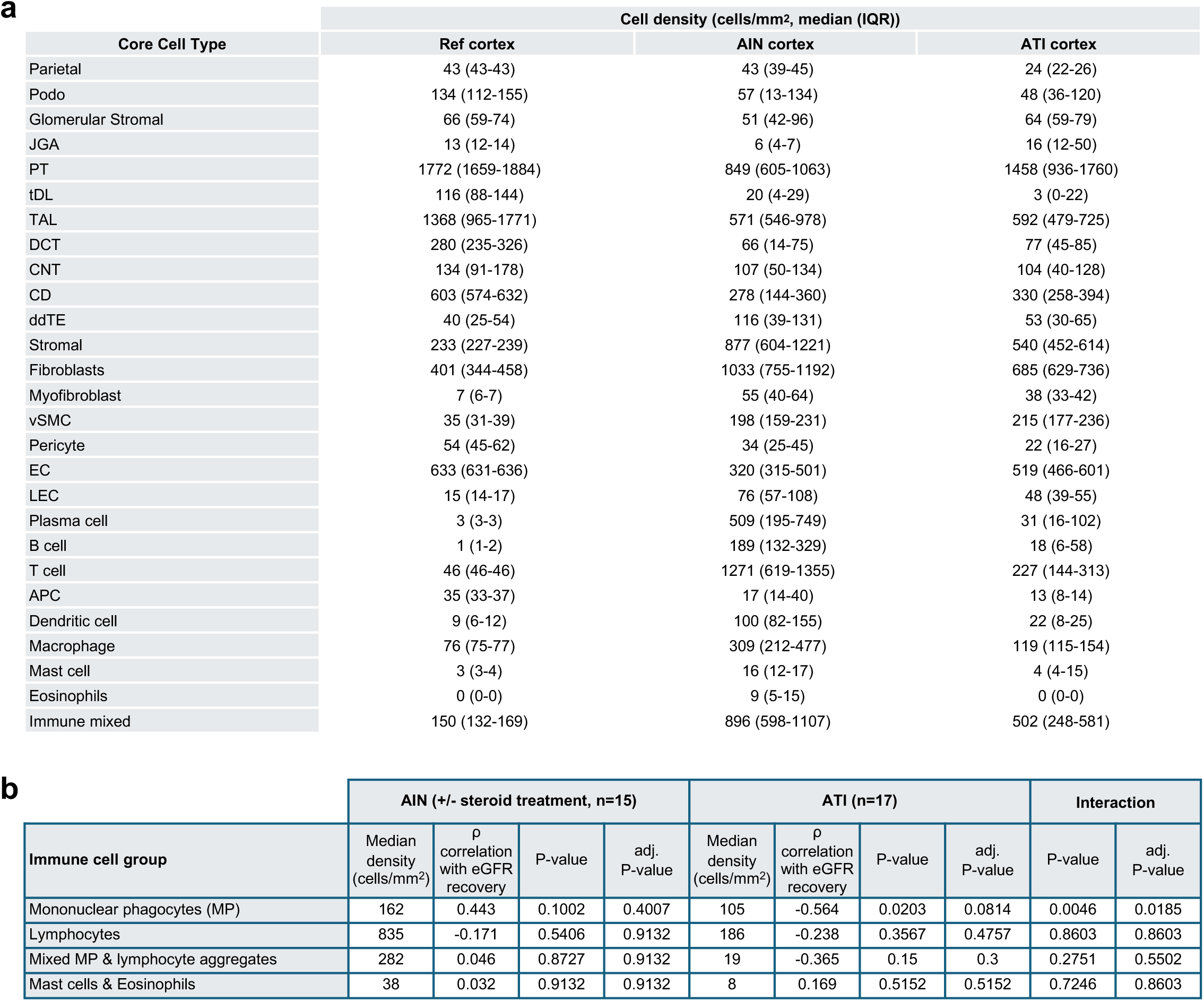
Spatial transcriptomics cell densities and immune infiltrate correlation with kidney function recovery.

**Extended Data Figure 8.**
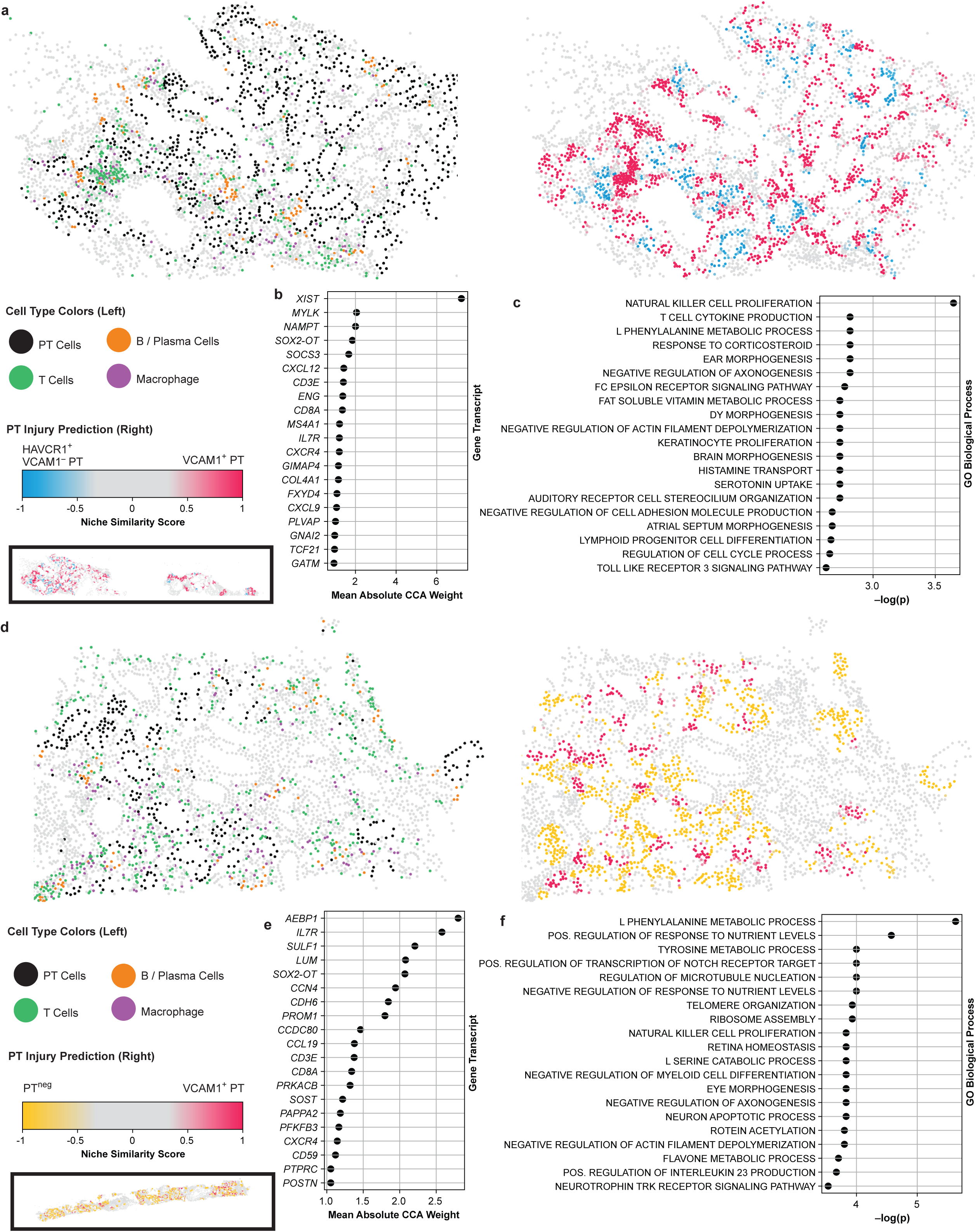
Comprehensive SORBET models for prediction of injured PT microenvironments in AIN and ATI cortex tissues.

## References

1. Ronco, C., R. Bellomo, and J.A. Kellum, Acute kidney injury. Lancet, 2019. 394(10212): p. 1949–1964.

2. Hoste, E.A.J., et al., Global epidemiology and outcomes of acute kidney injury. Nat Rev Nephrol, 2018. 14(10): p. 607–625.

3. Silver, S.A. and G.M. Chertow, The Economic Consequences of Acute Kidney Injury. Nephron, 2017. 137(4): p. 297–301.

4. Kellum, J.A., et al., Acute kidney injury. Nat Rev Dis Primers, 2021. 7(1): p. 52.

5. Baker, M.L. and L.G. Cantley, Adding insult to injury: the spectrum of tubulointerstitial responses in acute kidney injury. J Clin Invest, 2025. 135(6).

6. Muriithi, A.K., et al., Biopsy-proven acute interstitial nephritis, 1993-2011: a case series. Am J Kidney Dis, 2014. 64(4): p. 558–66.

7. Praga, M. and E. Gonzalez, Acute interstitial nephritis. Kidney Int, 2010. 77(11): p. 956–61.

8. Raghavan, R. and G. Eknoyan, Acute interstitial nephritis - a reappraisal and update. Clin Nephrol, 2014. 82(3): p. 149–62.

9. Sanchez-Alamo, B., C. Cases-Corona, and G. Fernandez-Juarez, Facing the Challenge of Drug-Induced Acute Interstitial Nephritis. Nephron, 2023. 147(2): p. 78–90.

10. Mose, F.H., et al., Prednisolone treatment in acute interstitial nephritis (PRAISE) - protocol for the randomized controlled trial. BMC Nephrol, 2021. 22(1): p. 161.

11. Wilson, C.B., Nephritogenic tubulointerstitial antigens. Kidney Int, 1991. 39(3): p. 501–17.

12. Fogo, A.B., et al., AJKD Atlas of Renal Pathology: Acute Interstitial Nephritis. Am J Kidney Dis, 2016. 67(6): p. e35–6.

13. Muriithi, A.K., S.H. Nasr, and N. Leung, Utility of urine eosinophils in the diagnosis of acute interstitial nephritis. Clin J Am Soc Nephrol, 2013. 8(11): p. 1857–62.

14. Prendecki, M., et al., Long-term outcome in biopsy-proven acute interstitial nephritis treated with steroids. Clin Kidney J, 2017. 10(2): p. 233–239.

15. Gonzalez, E., et al., Early steroid treatment improves the recovery of renal function in patients with drug-induced acute interstitial nephritis. Kidney Int, 2008. 73(8): p. 940–6.

16. Cheng, M., X. Gu, and G.A. Herrera, Dendritic cells in renal biopsies of patients with acute tubulointerstitial nephritis. Hum Pathol, 2016. 54: p. 113–20.

17. Ivanyi, B., et al., Acute tubulointerstitial nephritis: phenotype of infiltrating cells and prognostic impact of tubulitis. Virchows Arch, 1996. 428(1): p. 5–12.

18. Spanou, Z., et al., Involvement of drug-specific T cells in acute drug-induced interstitial nephritis. J Am Soc Nephrol, 2006. 17(10): p. 2919–27.

19. Muto, Y., et al., Epigenetic reprogramming driving successful and failed repair in acute kidney injury. Sci Adv, 2024. 10(32): p. eado2849.

20. Kirita, Y., et al., Cell profiling of mouse acute kidney injury reveals conserved cellular responses to injury. Proc Natl Acad Sci U S A, 2020. 117(27): p. 15874–15883.

21. Xu, L., et al., Immune-mediated tubule atrophy promotes acute kidney injury to chronic kidney disease transition. Nat Commun, 2022. 13(1): p. 4892.

22. Lake, B.B., et al., An atlas of healthy and injured cell states and niches in the human kidney. Nature, 2023. 619(7970): p. 585–594.

23. Dixon, E.E., et al., Spatially Resolved Transcriptomic Analysis of Acute Kidney Injury in a Female Murine Model. J Am Soc Nephrol, 2022. 33(2): p. 279–289.

24. Abedini, A., et al., Single-cell multi-omic and spatial profiling of human kidneys implicates the fibrotic microenvironment in kidney disease progression. Nat Genet, 2024. 56(8): p. 1712–1724.

25. Ikuta, H., et al., Idiopathic immune complex-mediated tubulointerstitial nephritis with hypocomplementemia and neutrophil-rich interstitial infiltrates. Clin Nephrol, 2018. 90(5): p. 357–362.

26. Kambham, N., et al., Idiopathic hypocomplementemic interstitial nephritis with extensive tubulointerstitial deposits. Am J Kidney Dis, 2001. 37(2): p. 388–99.

27. Nomura, A., et al., Role of complement in acute tubulointerstitial injury of rats with aminonucleoside nephrosis. Am J Pathol, 1997. 151(2): p. 539–47.

28. Thurman, J.M., et al., Acute tubular necrosis is characterized by activation of the alternative pathway of complement. Kidney Int, 2005. 67(2): p. 524–30.

29. Puthumana, J., et al., Biomarkers of inflammation and repair in kidney disease progression. J Clin Invest, 2021. 131(3).

30. Wen, Y., et al., Longitudinal biomarkers and kidney disease progression after acute kidney injury. JCI Insight, 2023. 8(9).

31. Shimonov, S., et al., SORBET: Automated cell-neighborhood analysis of spatial transcriptomics or proteomics for interpretable sample classification via GNN. bioRxiv, 2024.

32. Noel, S., et al., Immune Checkpoint Molecule TIGIT Regulates Kidney T Cell Functions and Contributes to AKI. J Am Soc Nephrol, 2023. 34(5): p. 755–771.

33. Moledina, D.G., et al., Identification and validation of urinary CXCL9 as a biomarker for diagnosis of acute interstitial nephritis. J Clin Invest, 2023. 133(13).

34. Liu, Y., et al., Single-cell and spatial transcriptome analyses reveal tertiary lymphoid structures linked to tumour progression and immunotherapy response in nasopharyngeal carcinoma. Nat Commun, 2024. 15(1): p. 7713.

35. Cortazar, F.B., et al., Clinical Features and Outcomes of Immune Checkpoint Inhibitor-Associated AKI: A Multicenter Study. J Am Soc Nephrol, 2020. 31(2): p. 435–446.

36. Gerhardt, L.M.S., et al., Single-nuclear transcriptomics reveals diversity of proximal tubule cell states in a dynamic response to acute kidney injury. Proc Natl Acad Sci U S A, 2021. 118(27).

37. Asghari, M., et al., Integration of spatial protein imaging and transcriptomics in the human kidney tracks the regenerative potential of proximal tubules. Sci Adv, 2025. 11(33): p. eadv8918.

38. Clark, A.J., M.C. Saade, and S.M. Parikh, The Significance of NAD+ Biosynthesis Alterations in Acute Kidney Injury. Semin Nephrol, 2022. 42(3): p. 151287.

39. Doke, T., et al., NAD(+) precursor supplementation prevents mtRNA/RIG-I-dependent inflammation during kidney injury. Nat Metab, 2023. 5(3): p. 414–430.

40. Nadour, Z., et al., Validation of a liquid chromatography coupled to tandem mass spectrometry method for simultaneous quantification of tryptophan and 10 key metabolites of the kynurenine pathway in plasma and urine: Application to a cohort of acute kidney injury patients. Clin Chim Acta, 2022. 534: p. 115–127.

41. Tran, M.T., et al., PGC1alpha drives NAD biosynthesis linking oxidative metabolism to renal protection. Nature, 2016. 531(7595): p. 528–32.

42. Kang, H.M., et al., Defective fatty acid oxidation in renal tubular epithelial cells has a key role in kidney fibrosis development. Nat Med, 2015. 21(1): p. 37–46.

43. Poyan Mehr, A., et al., De novo NAD(+) biosynthetic impairment in acute kidney injury in humans. Nat Med, 2018. 24(9): p. 1351–1359.

44. Sun, B.L., et al., Involvement of eNAMPT/TLR4 inflammatory signaling in progression of non-alcoholic fatty liver disease, steatohepatitis, and fibrosis. FASEB J, 2023. 37(3): p. e22825.

45. Melchinger, I., et al., VCAM-1 mediates proximal tubule-immune cell cross talk in failed tubule recovery during AKI-to-CKD transition. Am J Physiol Renal Physiol, 2024. 327(4): p. F610–F622.

46. Sacks, S.H. and W. Zhou, The role of complement in the early immune response to transplantation. Nat Rev Immunol, 2012. 12(6): p. 431–42.

47. Farrar, C.A., et al., Local extravascular pool of C3 is a determinant of postischemic acute renal failure. FASEB J, 2006. 20(2): p. 217–26.

48. Strainic, M.G., et al., Locally produced complement fragments C5a and C3a provide both costimulatory and survival signals to naive CD4+ T cells. Immunity, 2008. 28(3): p. 425–35.

49. Peng, Q., et al., C3a and C5a promote renal ischemia-reperfusion injury. J Am Soc Nephrol, 2012. 23(9): p. 1474–85.

50. Menke, J., et al., CXCL9, but not CXCL10, promotes CXCR3-dependent immune-mediated kidney disease. J Am Soc Nephrol, 2008. 19(6): p. 1177–89.

51. House, I.G., et al., Macrophage-Derived CXCL9 and CXCL10 Are Required for Antitumor Immune Responses Following Immune Checkpoint Blockade. Clin Cancer Res, 2020. 26(2): p. 487–504.

52. Dieu-Nosjean, M.C., et al., Tertiary lymphoid structures in cancer and beyond. Trends Immunol, 2014. 35(11): p. 571–80.

53. Manzo, A. and C. Pitzalis, Lymphoid tissue reactions in rheumatoid arthritis. Autoimmun Rev, 2007. 7(1): p. 30–34.

54. Schumacher, T.N. and D.S. Thommen, Tertiary lymphoid structures in cancer. Science, 2022. 375(6576): p. eabf9419.

55. Lee, S., et al., Distinct macrophage phenotypes contribute to kidney injury and repair. J Am Soc Nephrol, 2011. 22(2): p. 317–26.

56. Han, H.I., et al., The role of macrophages during acute kidney injury: destruction and repair. Pediatr Nephrol, 2019. 34(4): p. 561–569.

57. Kim, M.G., et al., The Role of M2 Macrophages in the Progression of Chronic Kidney Disease following Acute Kidney Injury. PLoS One, 2015. 10(12): p. e0143961.

58. Huen, S.C. and L.G. Cantley, Macrophages in Renal Injury and Repair. Annu Rev Physiol, 2017. 79: p. 449–469.

59. Boof, M.L., et al., Pharmacokinetics, pharmacodynamics and safety of the novel C-X-C chemokine receptor 3 antagonist ACT-777991: Results from the first-in-human study in healthy adults. Br J Clin Pharmacol, 2024. 90(2): p. 588–599.

60. Nester, C., et al., Developing Therapies for C3 Glomerulopathy: Report of the Kidney Health Initiative C3 Glomerulopathy Trial Endpoints Work Group. Clin J Am Soc Nephrol, 2024. 19(9): p. 1201–1208.

61. Dixon, B.P., et al., Clinical Safety and Efficacy of Pegcetacoplan in a Phase 2 Study of Patients with C3 Glomerulopathy and Other Complement-Mediated Glomerular Diseases. Kidney Int Rep, 2023. 8(11): p. 2284–2293.

62. Balzer, M.S., et al., Single-cell analysis highlights differences in druggable pathways underlying adaptive or fibrotic kidney regeneration. Nat Commun, 2022. 13(1): p. 4018.

63. Moledina, D.G., et al., Urine TNF-alpha and IL-9 for clinical diagnosis of acute interstitial nephritis. JCI Insight, 2019. 4(10).

64. Singh, N., et al., Development of a 2-dimensional atlas of the human kidney with imaging mass cytometry. JCI Insight, 2019. 4(12).

65. Avigan, Z.M., et al., Tubular Cell Dropout in Preimplantation Deceased Donor Biopsies as a Predictor of Delayed Graft Function. Transplant Direct, 2021. 7(7): p. e716.

66. Windhager, J., et al., An end-to-end workflow for multiplexed image processing and analysis. Nat Protoc, 2023. 18(11): p. 3565–3613.

67. Greenwald, N.F., et al., Whole-cell segmentation of tissue images with human-level performance using large-scale data annotation and deep learning. Nat Biotechnol, 2022. 40(4): p. 555–565.

68. contributors, n., napari: a multi-dimensional image viewer for python. 2019.

69. Hicks, S.A., et al., On evaluation metrics for medical applications of artificial intelligence. Sci Rep, 2022. 12(1): p. 5979.

70. Chevrier, S., et al., Compensation of Signal Spillover in Suspension and Imaging Mass Cytometry. Cell Syst, 2018. 6(5): p. 612–620 e5.

71. Haghverdi, L., et al., Batch effects in single-cell RNA-sequencing data are corrected by matching mutual nearest neighbors. Nat Biotechnol, 2018. 36(5): p. 421–427.

72. Levine, J.H., et al., Data-Driven Phenotypic Dissection of AML Reveals Progenitor-like Cells that Correlate with Prognosis. Cell, 2015. 162(1): p. 184–97.

73. Cook, D.P., et al., A Comparative Analysis of Imaging-Based Spatial Transcriptomics Platforms. bioRxiv, 2023: p. 2023.12.13.571385.

74. Wang, H., et al., Systematic benchmarking of imaging spatial transcriptomics platforms in FFPE tissues. bioRxiv, 2023.

