## Supplementary Figures and Legends for "Spatial analysis reveals the cellular microenvironments and mechanisms of inflammation and kidney injury in acute interstitial nephritis"

**Index of Supplementary Information**

**Extended Data:**

**Extended Data Figure 1 | Clinical characteristics and cellular landscapes of discovery and validation cohorts in acute interstitial nephritis.**

**a,** Demographics, laboratory features, and AKI-specific characteristics of the IMC discovery cohort (n=22 AIN, n=21 ATI, n=22 Reference). Data shown as median (interquartile range) for continuous variables or n (%) for categorical variables. P-values from Kruskal-Wallis test for overall comparison across three groups for continuous variables, with post-hoc Bonferroni-corrected pairwise comparisons for significant results. Pearson Chi-square or Fisher's Exact test used for categorical variables. For AIN vs ATI comparison, Mann-Whitney U test used for continuous variables and Chi-square or Fisher's Exact test for categorical variables.

**b,** Clinical characteristics of the spatial transcriptomics validation cohort. Data shown as median (IQR) or n (%) with statistical analysis as described in panel a. Kruskal-Wallis test performed for Age, Baseline eGFR, Days from baseline eGFR to AKI, Days from AKI to biopsy, eGFR at nadir, Change in eGFR from baseline to nadir, eGFR at 6 months, and Percent recovery at 6 months, with post-hoc Mann-Whitney U tests for pairwise comparisons.

**c,** Longitudinal eGFR trajectories showing baseline, nadir, and 6-month recovery in the discovery cohort. Data shown as median (IQR) with number of patients with available data. P-values from two-sided Mann-Whitney U test; exact significance displayed except for nadir eGFR, 6- and 12-month eGFR, and days from nadir to biopsy (asymptotic significance). Six-month follow-up measurements obtained 4-40 weeks post-biopsy, using values closest to 6-month timepoint.

**d,** Uniform Manifold Approximation and Projection (UMAP) visualization of core cell populations from IMC validation cohort (n=41 patients: 15 AIN, 16 ATI, 10 reference) showing 1,233,472 cells, using identical antibody panel, hybridization process, CyTOF core, and data analysis pipeline.

**Abbreviations**: PT, proximal tubule; TAL, thick ascending limb; DCT, distal convoluted tubule; CD, collecting duct; ddTE, dedifferentiated tubular epithelium; ECs, endothelial cells; LECs, lymphatic endothelial cells; vSMC, vascular smooth muscle cells; Mast cells; Eosinophils; Mixed immune cells as indicated. Statistical analyses performed using SPSS software and R. AIN, acute interstitial nephritis; ATI, acute tubular injury; Ref, reference.

**Extended Data Figure 2 | Tissue distribution and marker expression profiles of IMC-defined cell populations.**

**a,** Summary of tissue and cell counts across discovery, validation, and combined IMC cohorts. Reference tissues are stratified by source (tumor remote nephrectomy vs living donor). Numbers represent unique patient tissues and total cells analyzed. Discovery cohort: n=22 AIN, n=21 ATI, n=22 reference (9 tumor remote, 12 living donor) comprising 1,331,664 total cells. Validation cohort: n=15 AIN, n=16 ATI, n=10 reference (all tumor remote) comprising 1,233,472 total cells. Combined analysis includes 1,980,234 cells from cortical regions and 584,911 cells from medullary regions across both cohorts.

**b,** Dot plot showing protein expression patterns across 49 IMC-defined detailed cellular clusters from discovery cohort. Marker expression quantified as percentage of cells exceeding the 75th percentile of expression for each marker across all cells (dot size, range 0-100%) and mean log-transformed expression scaled per marker (color intensity, red = high, blue = low). Cell types ordered by anatomical compartment: glomerular (podocytes and associated cells), tubular (parietal cells through dedifferentiated tubular epithelium), endothelial, stromal, and immune populations. Vertical dashed lines delineate major compartments. Expression profiles distinguish functional states within cell lineages, including: healthy (H) vs dedifferentiated (dd) states in proximal tubule (PT) and thick ascending limb (TAL) cells; injury phenotypes (PT-INJ1 with KIM-1$^+$, PT-INJ2 with VCAM-1$^+$); proliferating subsets (Ki-67$^+$); ferroptosis-associated states (FACL4$^+$); and immune activation states (IL-9+ in T cells, CD206+ in mononuclear phagocytes). Marker panel includes 23 cell type-defining antibodies and 8 functional state markers (KIM-1, VCAM-1, Ki-67, MCP-1, FACL4, LC3B, TNF-α, IL-9).

**Abbreviations**: PT, proximal tubule; tDL, thin descending limb; TAL, thick ascending limb; DCT, distal convoluted tubule; CNT, connecting tubule; CD, collecting duct; ddTE, dedifferentiated tubular epithelium; Podo, podocytes; ECs, endothelial cells; LECs, lymphatic endothelial cells; MP, mononuclear phagocytes; Eos, eosinophils; UI, unidentified; uIS, undefined interstitial; uTE, undefined tubular epithelium; Prolif, proliferating.

**Extended Data Figure 3 | Spatial transcriptomics cellular landscapes and T cell subset characterization.**

**a,** Distribution of cells and unique tissue sections analyzed by spatial transcriptomics across disease categories and anatomical regions. Data from 20 tissues (8 AIN, 7 ATI, 5 reference) generating 321,318 total cells. Size of numbers visually represented by color gradient where red is highest, blue is lowest.

**b,** Dot plot showing gene expression patterns across 26 core cell types and one mixed population identified through unsupervised clustering of spatial transcriptomics data. Expression quantified as percentage of cells expressing each marker (dot size, 0-75%) and scaled mean expression (color intensity, red = high, blue = low). Cell types ordered anatomically from glomerular through tubular epithelium, stromal, vascular, and immune populations. Horizontal dashed lines separate major anatomical compartments. Gene panel includes 5,101 genes from 10x Genomics Xenium Prime 5K Human Pan Tissue & Pathways Panel plus 100 custom kidney-specific genes. Data aggregated from all 20 tissues used in ST excluding cells classified as low quality (<10 genes expressed).

**c,** *CD4*$^{+}$ T cell subset analysis showing marker expression across six functionally distinct populations. Subsets identified through targeted re-clustering using lineage and activation markers: Naive, Memory (*IL7R*$^{+}$/*CCR7*$^{+}$), TRM (tissue-resident memory; *CD69*$^{+}$/*ITGAE*$^{+}$), interferon (IFN)-activated (*CXCL9*$^{+}$/*CXCL10*$^{+}$/*STAT1*$^{+}$), regulatory ($T_{reg}$, *FOXP3*$^{+}$/*CTLA4*$^{+}$), and Proliferating (*MKI67*$^{+}$). Expression shown as percentage of cells positive (dot size) and scaled expression z-score (color). Horizontal dotted lines separate functional marker groups**.**

**d,** CD8$^{+}$ T cell subset analysis revealing eight distinct populations through expression of lineage, activation, and exhaustion markers. Subsets include: Resting, Memory (*IL7R*$^{+}$/*CCR7*$^{+}$), Tfc (follicular cytotoxic; *CXCL13*$^{+}$/*ICOS*$^{+}$), TRM (tissue-resident memory; *CD69*$^{+}$/*ITGAE*$^{+}$), interferon (IFN)-activated (*CXCL9*$^{+}$/*CXCL10*$^{+}$), exhausted (*LAG3*$^{+}$*TIGIT*$^{+}$*CD69*$^{-}$*ITGAE*$^{-}$), proliferating (*MKI67*$^{+}$), and checkpoint-high TRM *(LAG3*$^{+}$*/TIGIT*$^{+}$*/PDCD1*$^{+}$*)*.

**Extended Data Figure 4 | Proximal tubule injury phenotyping and cellular distribution analysis across discovery and validation IMC cohorts.**

**a,** Discovery cohort: Pathologist-adjudicated injury scores and cellular neighborhood analysis for proximal tubule (PT) cell states. Injury scores assessed on H&E-stained adjacent sections (scale 1-5, with 5 representing severe injury) based on presence. PT-INJ2 cells exhibited chronic tubular atrophy morphology precluding conventional injury scoring (N/A). Nearby vascular endothelial cells and interstitial leukocytes quantified within 20µm radius of each tubular cross-section.

**b,** Discovery cohort: Distribution of tubular epithelial cell states in cortex tissues across tissue types. Percentages indicate proportion of each cell state within total cells of that lineage. P-values from Kruskal-Wallis test for overall comparison across three tissue types.

**c,** Discovery cohort: Comprehensive analysis of cortical cell densities (cells/mm²) across 45 cell types, excluding glomerular cell types. Data shown as median (IQR). Statistical analysis by Kruskal-Wallis test with post-hoc pairwise Wilcoxon rank-sum tests using Bonferroni correction.

**d,** Validation cohort: Marker expression profiles of PT cell states. Dot size represents percentage of cells exceeding the 75th percentile of expression across all tubular epithelial cells. Color indicates scaled log-transformed expression (red = high, blue = low). Key markers distinguish functional states: megalin/AQP1 (healthy), vimentin (dedifferentiated), KIM-1/VCAM-1 (injury), FACL4 (ferroptosis), Ki-67 (proliferation).

**e,** Validation cohort: Alluvial diagram visualizing the proportional shifts in PT cell populations in cortex tissues across tissue types. Height of streams represent relative abundance.

**f,** Validation cohort: Quantitative comparison of PT cell densities in cortex tissues by tissue type. Box plots show median and IQR with individual patient values overlaid. Statistical analysis by pairwise Wilcoxon rank-sum tests with Bonferroni correction. Asterisks indicate significance: *$p<0.05$, **$p<0.01$, ***$p<0.001$.

**g,** Validation cohort: Immune cell densities in cortex tissues stratified by tissue type. Analysis focused on lymphocyte subsets (CD4$^{+}$ T cells, CD8$^{+}$ T cells, CD4$^{low/-}$CD8$^{low/-}$ T cells, mixed B and T cell aggregates), mononuclear phagocytes (MP & T cells, MP-CD206$^{+}$, MP-CD206$^{-}$), and mast cells. Box plots with overlaid data points; statistical testing as in panel f.

**Extended Data Figure 5 | Spatial analysis of immune cell neighborhoods surrounding proximal tubule injury states.**
**a,** Quantification of immediately adjacent neighboring cells within 20μm radius of proximal tubule (PT) cell states across cortex tissue types in the IMC discovery cohort (n=62, AIN=21, ATI=21, Ref=21). Table shows the mean number of each neighbor cell type per index PT cell, with the number of patients exhibiting at least one index-neighbor pair. Fold changes (β) represent ratios of mean neighbor counts between tissue types, with q-values adjusted for multiple comparisons using Benjamini-Hochberg correction.
**b,** Comparative analysis of cellular neighborhoods between PT injury states within AIN cortex tissues only (n=21). Mean expression represents average number of neighboring cells within 20μm per index PT cell, with PT-H serving as reference (β=1). Beta values indicate fold change relative to PT-H neighborhoods. P-values from negative binomial regression without adjustment for multiple comparisons.
**Abbreviations**: ECs, endothelial cells; LECs, lymphatic endothelial cells; MPs, mononuclear phagocytes.

**Extended Data Figure 6 | Differential gene expression analysis of proximal tubule injury states within AIN cortex and between AIN and ATI cortex tissues.** Heatmap showing log2 fold changes of differentially expressed genes (DEGs) across proximal tubule cell comparisons from spatial transcriptomics data of cortex tissues (n=16, AIN=7, ATI=6, Ref=2). Analysis includes three within-AIN comparisons (red header): *HAVCR1*$^{+}$*VCAM1*$^{-}$ PT vs *HAVCR1*$^{-}$*VCAM1*$^{-}$ PT (PT$^{neg}$), *VCAM1*$^{+}$ PT vs PT$^{neg}$, and *VCAM1*$^{+}$ PT vs *HAVCR1*$^{+}$*VCAM1*$^{-}$ PT; and three between-tissue comparisons of matched cell states (green header): AIN vs ATI in PT$^{neg}$, *HAVCR1*$^{+}$*VCAM1*$^{-}$ PT, and *VCAM1*$^{+}$ PT populations. Genes displayed include top 20 most significant DEGs from each comparison plus key injury and inflammatory markers (*HAVCR1*, *VCAM1*, *C3*, *LCN2*, *SPP1*, *VIM*, *CD74*, *STAT1*, *CFB*, *ITGB6*, *THY1*, *SERPINA3*). Color scale represents log2 fold change from -4 (blue, downregulated) to +4 (red, upregulated). Significance levels indicated by asterisks: *p<0.05, **p<0.01, ***p<0.001 (adjusted p-values, Wilcoxon rank-sum test with Bonferroni correction). Genes clustered by Ward.D2 hierarchical clustering based on Euclidean distance.

**Extended Data Figure 7 | Spatial transcriptomics cell densities and immune infiltrate correlation with kidney function recovery.**
**a,** Cell densities (cells/mm²) for core cell types identified in spatial transcriptomics analysis of cortical tissues. Data shown as median (IQR) across reference (n=2 tissues), AIN (n=7), and ATI (n=6) cortical samples.
**b,** Correlation analysis between immune cell group densities in cortex tissues in IMC discovery cohort and eGFR recovery at 6 months post-biopsy. Analysis included all AIN patients with available clinical follow-up data (n=15, including both patients who did and did not receive steroid treatment). Immune cell groups comprised: mononuclear phagocytes (all CD206$^{+}$, CD206$^{-}$, and proliferating subsets combined), lymphocytes (all T cell subsets, B cells, and plasma cells combined), mixed mononuclear phagocyte & lymphocyte aggregates, and combined mast cells & eosinophils. Spearman correlation coefficients (ρ) calculated for each group with eGFR recovery (percent of baseline eGFR recovered at 6 months). ATI patients (n=17) shown for comparison. P-values adjusted for multiple comparisons using Benjamini-Hochberg method. Interaction p-values test whether the correlation between immune density and recovery differs significantly between AIN and ATI cortex tissues.

**Extended Data Figure 8 | Comprehensive SORBET models for prediction of injured PT microenvironments in AIN and ATI cortex tissues.**
**a–c,** Overall SORBET model for analysis of AIN cortex for 3-way discriminatory prediction of PT cell type based on microenvironment. **a,** Comparison of cell type labels (left; see legend, lower left) with cell-microenvironment scores (right; see legend, lower left), which estimate the predictive utility of each cell (and its microenvironment) for PT injury state. Blue indicates utility for *HAVCR1*$^{+}$/*VCAM1*$^{-}$ PT cell prediction; red indicates *VCAM1*$^{+}$. Cell type labels were not used as part of model for prediction. (Inset Box) Full tissue sample, colored by cell-niche score. **b,** Top 20 markers ranked by mean absolute weight from canonical correlation analysis (CCA) comparing input data to the inferred representation of each cell. Higher weight indicates stronger predictive utility for 3-way PT cell state discrimination (see also **Supplementary Fig. 3**). **c,** Top 20 Gene Ontology Biological Process (GO BP) gene sets enriched among non-zero markers identified by CCA. Significance was estimated with a hypergeometric test; false discovery rate was controlled at 0.01 (Benjamini–Hochberg).
**d–f,** Equivalent analysis of ATI cortex, with alternate display of PT$^{neg}$ vs *HAVCR1*$^{+}$/*VCAM1*$^{-}$ PT cell microenvironment prediction. **d,** Comparison of cell types (left) with cell-microenvironment scores (right). Mixed cellular patterns characterize the PT$^{neg}$ phenotype. **e,** Top 20 markers ranked by mean absolute CCA weight (see also **Supplementary Fig. 4**). **f,** GO BP gene sets enriched among CCA-identified markers.

**Supplementary Figures:**

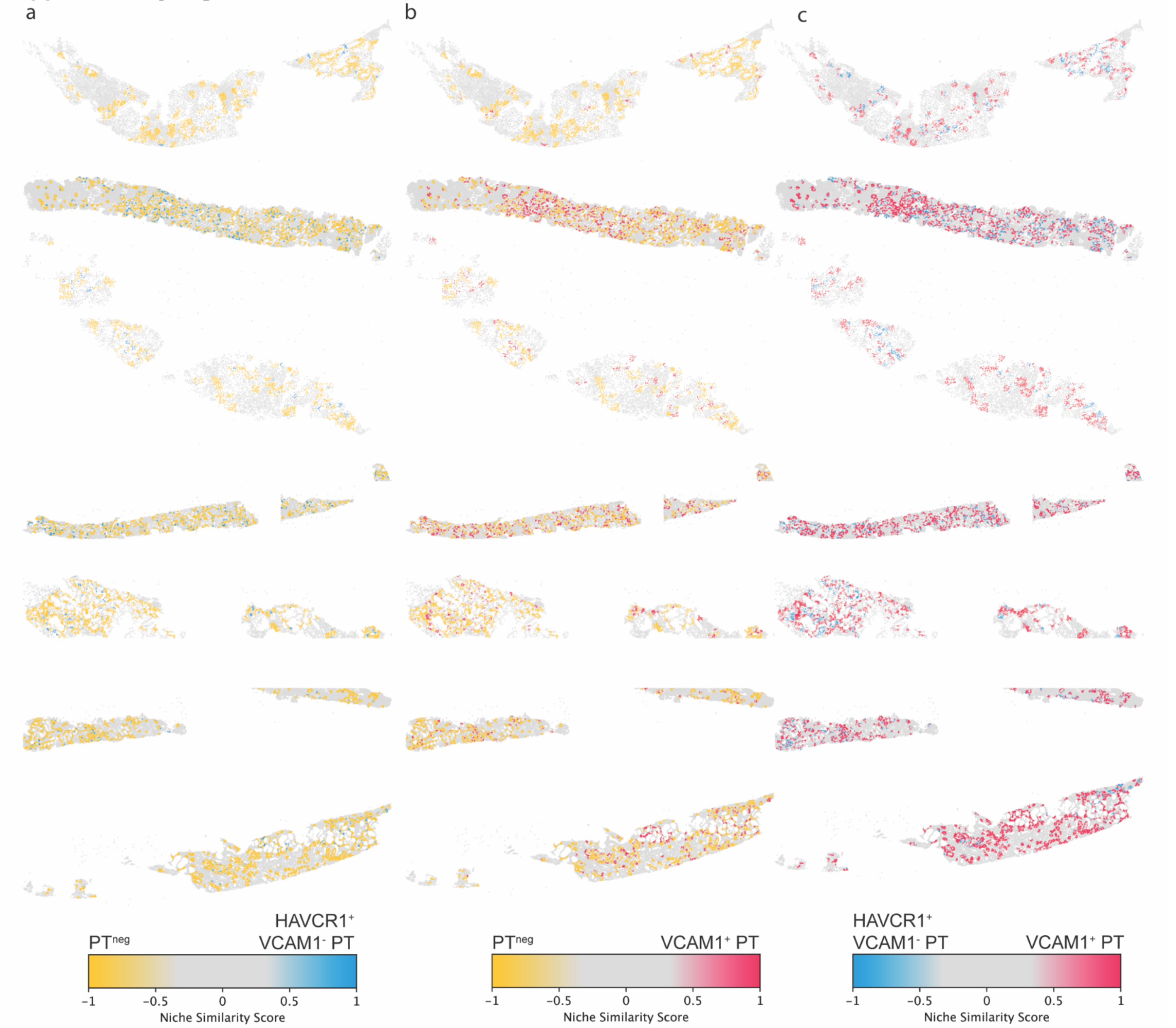


**Supplementary Figure 1 | SORBET-derived cellular niche analysis reveals distinct microenvironmental signatures of proximal tubule injury states in AIN cortex.** Spatial transcriptomics data from AIN kidney cortex samples analyzed using SORBET (Spatial Optimal transport for Representation of Biological Environmental Topology), a geometric deep learning framework that identifies disease-relevant cellular neighborhoods through graph convolutional networks. SORBET extracts subgraphs centered on each cell, generates cell-niche embeddings (CNEs) capturing local cellular composition and spatial organization within 2-hop neighborhoods (~50μm radius), and computes inverse-distance weighted scores (IDWS) to quantify microenvironmental similarity between cell states. Each panel displays identical tissue sections with cells colored by pairwise niche similarity scores: **a,** $PT^{neg}$ (*HAVCR1*$^{-}$*VCAM1*$^{-}$ PT) vs. *HAVCR1*$^{+}$*VCAM1*$^{-}$ PT cells; **b,** $PT^{neg}$ vs. *VCAM1*$^{+}$PT cells; **c,** *HAVCR1*$^{+}$*VCAM1*$^{-}$ PT vs. *VCAM1*$^{+}$ PT cells. Positive scores indicate microenvironments resembling the second cell state in each comparison, while negative scores similarity to the first state. Gray represents cells excluded from analysis. Spatial segregation patterns reveal hierarchical organization of injury microenvironments: $PT^{neg}$ niches (yellow, panels a-b) occupy distinct spatial domains from *HAVCR1*$^{+}$*VCAM1*$^{-}$ states, while *HAVCR1*$^{+}$*VCAM1*$^{-}$ and *VCAM1*$^{+}$ PT niches show substantial spatial overlap (panel c), suggesting shared inflammatory microenvironments. The progressive transition from segregated (healthy vs injured) to overlapping (injured subtypes) niche patterns indicates that microenvironmental remodeling precedes or accompanies phenotypic changes during AIN progression. Scale bars represent normalized niche similarity scores (-1 to 1) computed using SORBET's probabilistic model with max pooling across CNE dimensions.

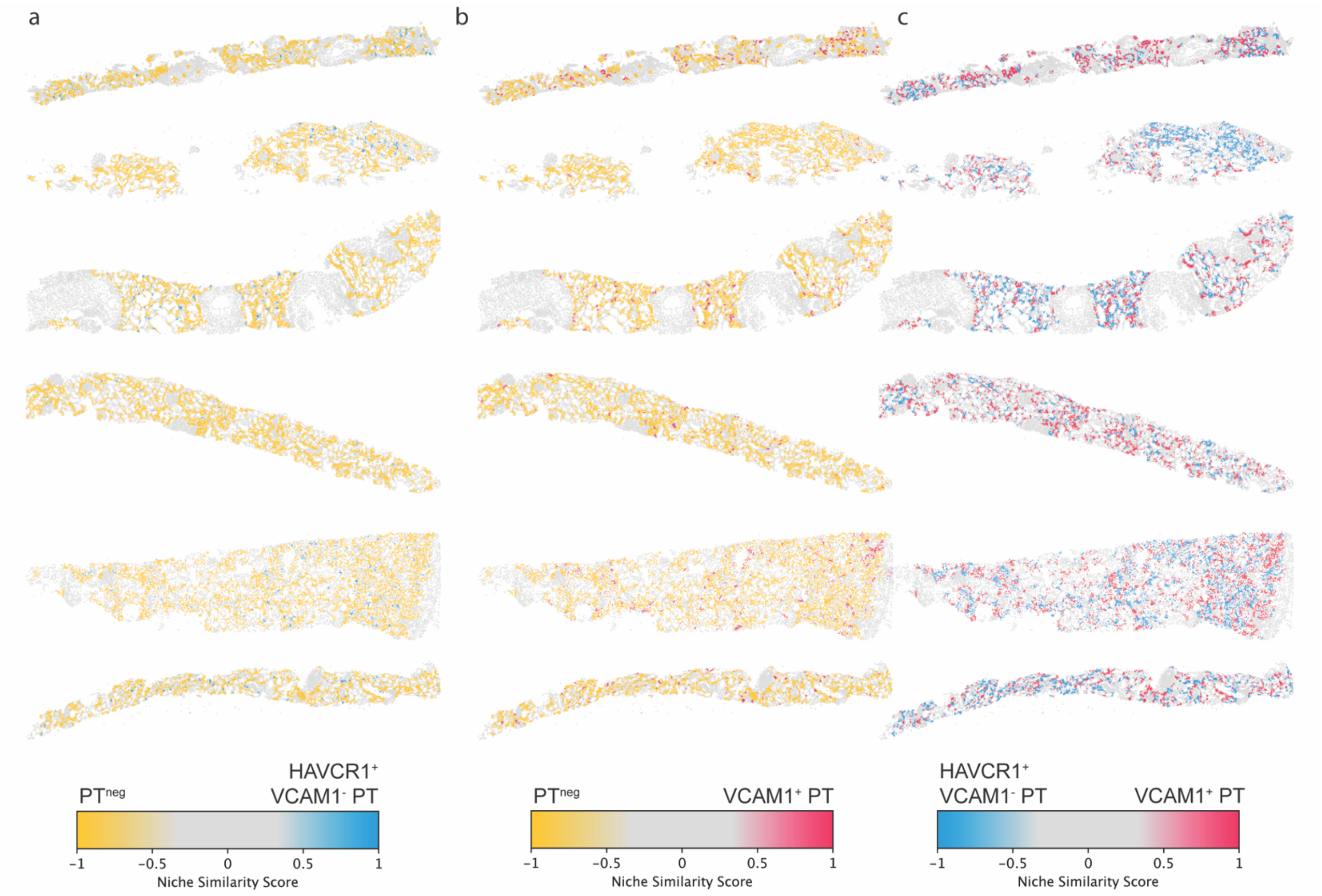


**Supplementary Figure 2 | SORBET-derived cellular niche analysis reveals distinct microenvironmental signatures of proximal tubule injury states in ATI cortex.** Spatial transcriptomics data from ATI kidney cortex samples analyzed using SORBET (Spatial Optimal transport for Representation of Biological Environmental Topology), a geometric deep learning framework that identifies disease-relevant cellular neighborhoods through graph convolutional networks. SORBET extracts subgraphs centered on each cell, generates cell-niche embeddings (CNEs) capturing local cellular composition and spatial organization within 2-hop neighborhoods (~50μm radius), and computes inverse-distance weighted scores (IDWS) to quantify microenvironmental similarity between cell states. Each panel displays identical tissue sections with cells colored by pairwise niche similarity scores: **a,** $PT^{neg}$ ($HAVCR1^{-}VCAM1^{-}$ PT) versus $HAVCR1^{+}VCAM1^{-}$ PT cells; **b,** $PT^{neg}$ versus $VCAM1^{+}$ PT cells; **c,** $HAVCR1^{+}VCAM1^{-}$ PT versus $VCAM1^{+}$ PT cells. Positive scores indicate microenvironments resembling the second cell state in each comparison, while negative scores indicate similarity to the first state. Gray represents cells excluded from analysis. Spatial segregation patterns reveal hierarchical organization of injury microenvironments: $PT^{neg}$ niches (yellow, panels a-b) occupy distinct spatial domains from $HAVCR1^{+}VCAM1^{-}$ states, while $HAVCR1^{+}VCAM1^{-}$ and $VCAM1^{+}$ PT niches show substantial spatial overlap (panel c), suggesting shared inflammatory microenvironments despite distinct injury markers. The progressive transition from segregated ($PT^{neg}$ vs $HAVCR1^{+}VCAM1^{-}$) to overlapping ($HAVCR1^{+}VCAM1^{-}$ vs $VCAM1^{+}$ PT) niche patterns indicates that microenvironmental remodeling precedes or accompanies phenotypic changes during ATI progression. Scale bars represent normalized niche similarity scores (-1 to 1) computed using SORBET's probabilistic model with max pooling across CNE dimensions.

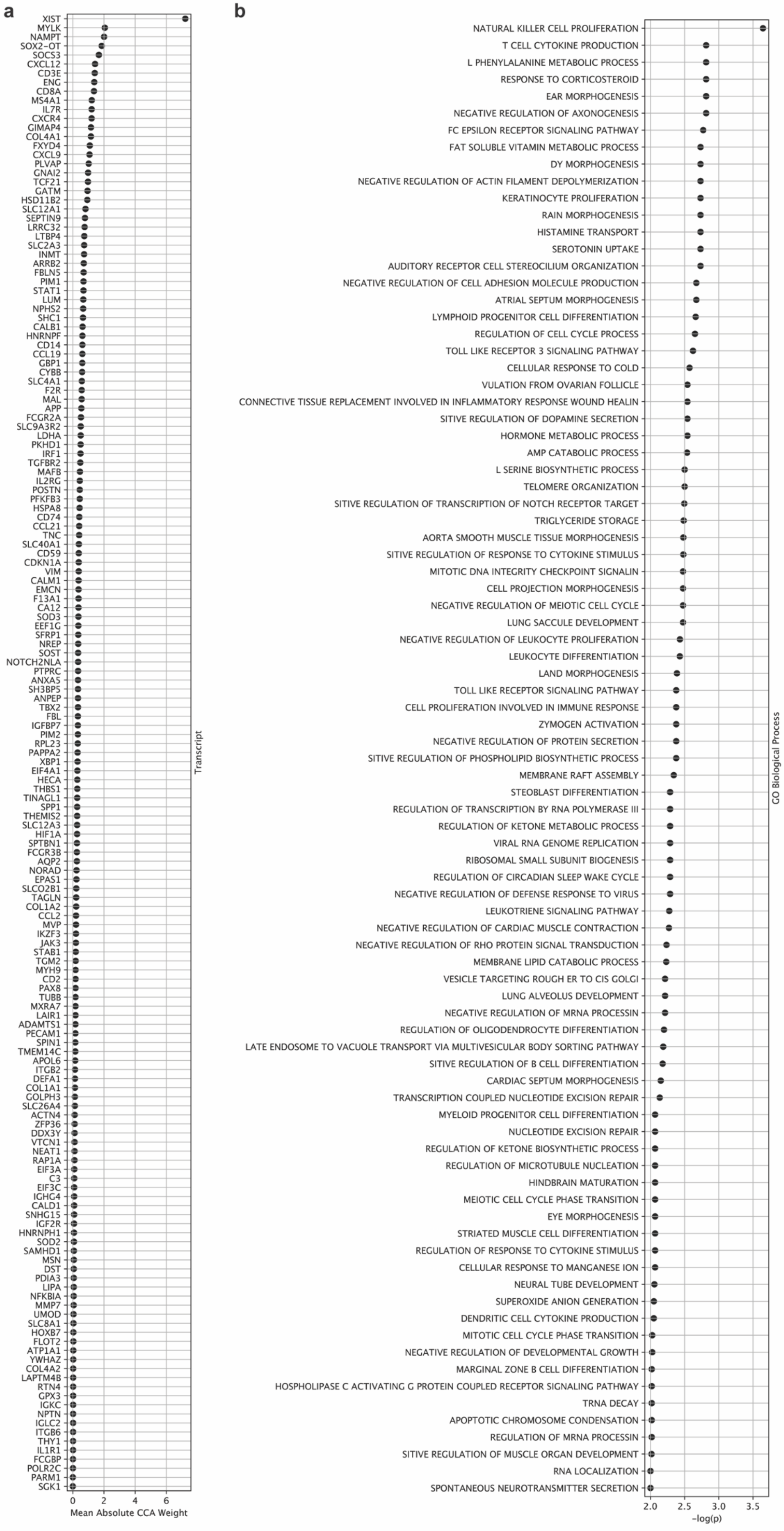
a
XIST
MYLK
NAMPT
SOX2-OT
SOCS3
CXCL12
CD3E
ENG
CD8A
MS4A1
IL7R
CXCR4
GIMAP4
COL4A1
FXYD4
CXCL9
PLVAP
GNAI2
TCF21
GATM
HSD11B2
SLC12A1
SEPTIN9
LRRC32
LTBP4
SLC2A3
INMT
ARRB2
FBLN5
PIM1
STAT1
LUM
NPHS2
SHC1
CALB1
HNRNPF
CD14
CCL19
GBP1
CYBB
SLC4A1
F2R
MAL
APP
FCGR2A
SLC9A3R2
LDHA
PKHD1
IRF1
TGFBR2
MAFB
IL2RG
POSTN
PFKFB3
HSPA8
CD74
CCL21
TNC
SLC40A1
CD59
CDKN1A
VIM
CALM1
EMCN
F13A1
CA12
SOD3
EEF1G
SFRP1
NREP
SOST
NOTCH2NLA
PTPRC
ANXA5
SH3BP5
ANPEP
TBX2
FBL
IGFBP7
PIM2
RPL23
PAPPA2
XBP1
EIF4A1
HECA
THBS1
TINAGL1
SPP1
THEMIS2
SLC12A3
HIF1A
SPTBN1
FCGR3B
AQP2
NORAD
EPAS1
SLCO2B1
TAGLN
COL1A2
CCL2
MVP
IKZF3
JAK3
STAB1
TGM2
MYH9
CD2
PAX8
TUBB
MXRA7
LAIR1
ADAMTS1
PECAM1
SPIN1
TMEM14C
APOL6
ITGB2
DEFA1
COL1A1
GOLPH3
SLC26A4
ACTN4
ZFP36
DDX3Y
VTCN1
NEAT1
RAP1A
EIF3A
C3
EIF3C
IGHG4
CALD1
SNHG15
IGF2R
HNRNPH1
SOD2
SAMHD1
MSN
DST
PDIA3
LIPA
NFKBIA
MMP7
UMOD
SLC8A1
HOXB7
FLOT2
ATP1A1
YWHAZ
COL4A2
LAPTM4B
RTN4
GPX3
IGKC
NPTN
IGLC2
ITGB6
THY1
IL1R1
FCGBP
POLR2C
PARM1
SGK1
0
2
4
6
Mean Absolute CCA Weight
Transcript
b
NATURAL KILLER CELL PROLIFERATION
T CELL CYTOKINE PRODUCTION
L PHENYLALANINE METABOLIC PROCESS
RESPONSE TO CORTICOSTEROID
EAR MORPHOGENESIS
NEGATIVE REGULATION OF AXONOGENESIS
FC EPSILON RECEPTOR SIGNALING PATHWAY
FAT SOLUBLE VITAMIN METABOLIC PROCESS
DY MORPHOGENESIS
NEGATIVE REGULATION OF ACTIN FILAMENT DEPOLYMERIZATION
KERATINOCYTE PROLIFERATION
RAIN MORPHOGENESIS
HISTAMINE TRANSPORT
SEROTONIN UPTAKE
AUDITORY RECEPTOR CELL STEREOCILIUM ORGANIZATION
NEGATIVE REGULATION OF CELL ADHESION MOLECULE PRODUCTION
ATRIAL SEPTUM MORPHOGENESIS
LYMPHOID PROGENITOR CELL DIFFERENTIATION
REGULATION OF CELL CYCLE PROCESS
TOLL LIKE RECEPTOR 3 SIGNALING PATHWAY
CELLULAR RESPONSE TO COLD
VULATION FROM OVARIAN FOLLICLE
CONNECTIVE TISSUE REPLACEMENT INVOLVED IN INFLAMMATORY RESPONSE WOUND HEALIN
SITIVE REGULATION OF DOPAMINE SECRETION
HORMONE METABOLIC PROCESS
AMP CATABOLIC PROCESS
L SERINE BIOSYNTHETIC PROCESS
TELOMERE ORGANIZATION
SITIVE REGULATION OF TRANSCRIPTION OF NOTCH RECEPTOR TARGET
TRIGLYCERIDE STORAGE
AORTA SMOOTH MUSCLE TISSUE MORPHOGENESIS
SITIVE REGULATION OF RESPONSE TO CYTOKINE STIMULUS
MITOTIC DNA INTEGRITY CHECKPOINT SIGNALIN
CELL PROJECTION MORPHOGENESIS
NEGATIVE REGULATION OF MEIOTIC CELL CYCLE
LUNG SACCULE DEVELOPMENT
NEGATIVE REGULATION OF LEUKOCYTE PROLIFERATION
LEUKOCYTE DIFFERENTIATION
LAND MORPHOGENESIS
TOLL LIKE RECEPTOR SIGNALING PATHWAY
CELL PROLIFERATION INVOLVED IN IMMUNE RESPONSE
ZYMOGEN ACTIVATION
NEGATIVE REGULATION OF PROTEIN SECRETION
SITIVE REGULATION OF PHOSPHOLIPID BIOSYNTHETIC PROCESS
MEMBRANE RAFT ASSEMBLY
STEOBLAST DIFFERENTIATION
REGULATION OF TRANSCRIPTION BY RNA POLYMERASE III
REGULATION OF KETONE METABOLIC PROCESS
VIRAL RNA GENOME REPLICATION
RIBOSOMAL SMALL SUBUNIT BIOGENESIS
REGULATION OF CIRCADIAN SLEEP WAKE CYCLE
NEGATIVE REGULATION OF DEFENSE RESPONSE TO VIRUS
LEUKOTRIENE SIGNALING PATHWAY
NEGATIVE REGULATION OF CARDIAC MUSCLE CONTRACTION
NEGATIVE REGULATION OF RHO PROTEIN SIGNAL TRANSDUCTION
MEMBRANE LIPID CATABOLIC PROCESS
VESICLE TARGETING ROUGH ER TO CIS GOLGI
LUNG ALVEOLUS DEVELOPMENT
NEGATIVE REGULATION OF MRNA PROCESSIN
REGULATION OF OLIGODENDROCYTE DIFFERENTIATION
LATE ENDOSOME TO VACUOLE TRANSPORT VIA MULTIVESICULAR BODY SORTING PATHWAY
SITIVE REGULATION OF B CELL DIFFERENTIATION
CARDIAC SEPTUM MORPHOGENESIS
TRANSCRIPTION COUPLED NUCLEOTIDE EXCISION REPAIR
MYELOID PROGENITOR CELL DIFFERENTIATION
NUCLEOTIDE EXCISION REPAIR
REGULATION OF KETONE BIOSYNTHETIC PROCESS
REGULATION OF MICROTUBULE NUCLEATION
HINDBRAIN MATURATION
MEIOTIC CELL CYCLE PHASE TRANSITION
EYE MORPHOGENESIS
STRIATED MUSCLE CELL DIFFERENTIATION
REGULATION OF RESPONSE TO CYTOKINE STIMULUS
CELLULAR RESPONSE TO MANGANESE ION
NEURAL TUBE DEVELOPMENT
SUPEROXIDE ANION GENERATION
DENDRITIC CELL CYTOKINE PRODUCTION
MITOTIC CELL CYCLE PHASE TRANSITION
NEGATIVE REGULATION OF DEVELOPMENTAL GROWTH
MARGINAL ZONE B CELL DIFFERENTIATION
HOSPHOLIPASE C ACTIVATING G PROTEIN COUPLED RECEPTOR SIGNALING PATHWAY
TRNA DECAY
APOPTOTIC CHROMOSOME CONDENSATION
REGULATION OF MRNA PROCESSIN
SITIVE REGULATION OF MUSCLE ORGAN DEVELOPMENT
RNA LOCALIZATION
SPONTANEOUS NEUROTRANSMITTER SECRETION
2.0
2.5
3.0
3.5
-log(p)
GO Biological Process

**Supplementary Figure 3 | Extended canonical correlation analysis identifies transcriptional programs associated with proximal tubule injury microenvironments in AIN cortex. a,** Ranked genes from canonical correlation analysis (CCA) performed by SORBET to identify transcriptional features in cellular neighborhoods that differentiate PT cell states ($PT^{neg}$, *HAVCR1*$^{+}$*VCAM1*$^{-}$ PT, *VCAM1*$^{+}$ PT) in AIN tissues. The sparse CCA algorithm correlates input gene expression with cell-niche embeddings (CNEs) generated through SORBET's graph convolutional network analysis of 2-hop spatial neighborhoods. Genes ranked by mean absolute weight across CCA components, with higher weights indicating stronger contribution to distinguishing among the three PT cell states. **b,** Gene set enrichment analysis of SORBET-identified genes revealing biological processes that distinguish PT cell states. Pathways ranked by enrichment score. The $\ell_0$-based sparse CCA implementation with orthogonal regularization and two-fold cross-validation ensures robust feature selection across the heterogeneous spatial transcriptomics dataset.

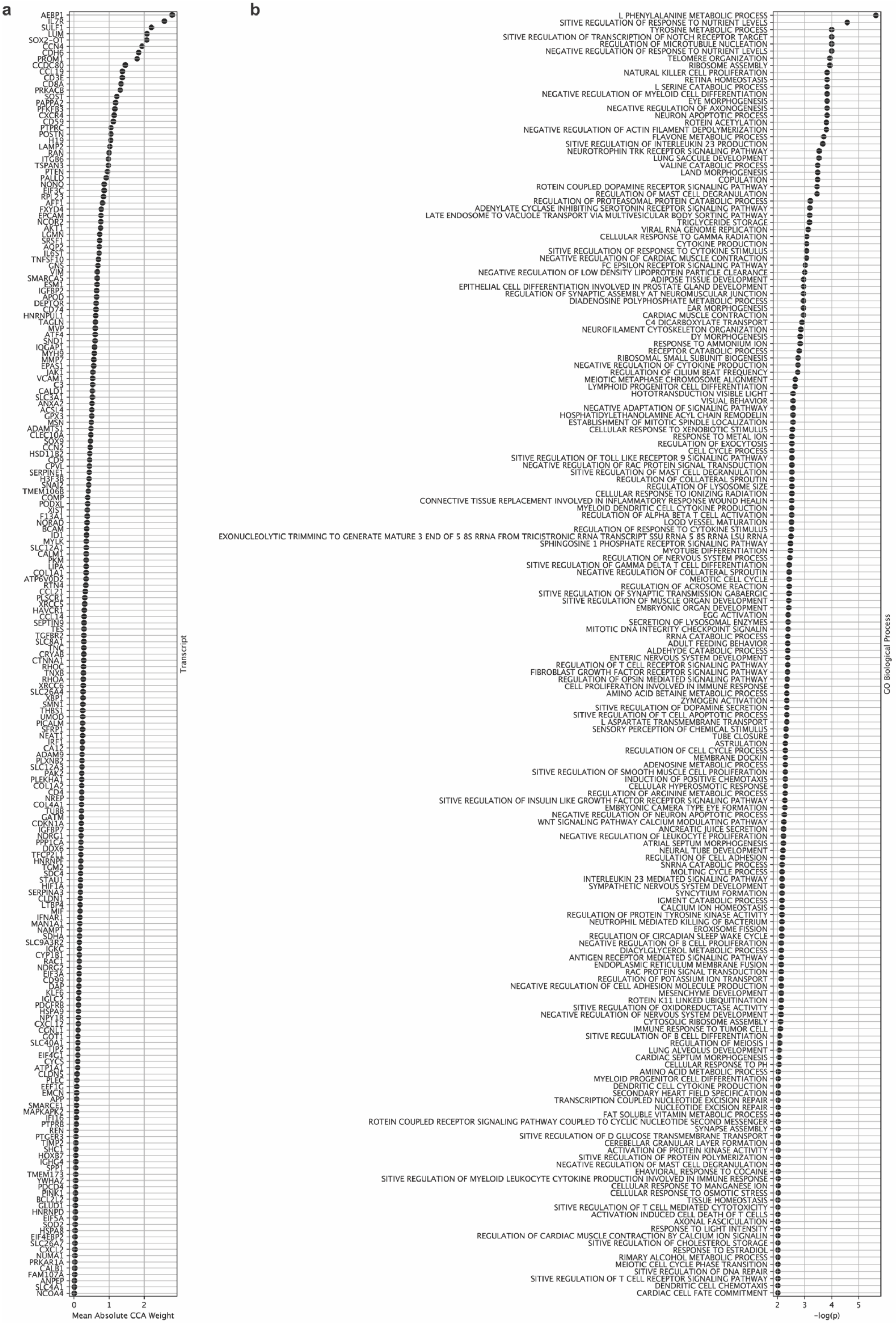
a
Transcript
Mean Absolute CCA Weight
0
1
2
b
GO Biological Process
-log(p)
2
3
4
5

**Supplementary Figure 4 | Extended canonical correlation analysis identifies transcriptional programs associated with proximal tubule injury microenvironments in ATI cortex. a,** Ranked genes from canonical correlation analysis (CCA) performed by SORBET to identify transcriptional features in cellular neighborhoods that differentiate PT cell states ($PT^{neg}$, *HAVCR1*$^{+}$*VCAM1*$^{-}$ PT, *VCAM1*$^{+}$ PT) in ATI tissues. The sparse CCA algorithm correlates input gene expression with cell-niche embeddings (CNEs) generated through SORBET's graph convolutional network analysis of 2-hop spatial neighborhoods (~50μm radius). Genes ranked by mean absolute weight across CCA components, with higher weights indicating stronger contribution to distinguishing among the three PT cell states. **b,** Gene set enrichment analysis of SORBET-identified genes revealing biological processes that distinguish PT cell states. Pathways ranked by enrichment score. The $\ell0$-based sparse CCA implementation with orthogonal regularization and two-fold cross-validation ensures robust feature selection. Comparison with AIN SORBET results (**Supplementary Fig. 4**) reveals both shared injury responses and disease-specific microenvironmental programs.

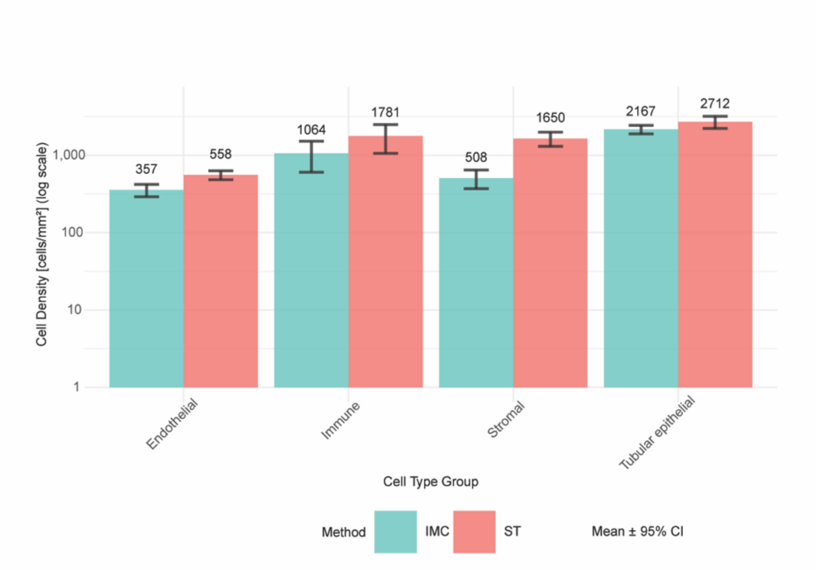


**Supplementary Figure 5 | Cross-platform validation of cell density measurements between spatial transcriptomics and imaging mass cytometry.** Comparison of cell densities (cells/mm²) between spatial transcriptomics (Xenium, red) and imaging mass cytometry (IMC, teal) platforms from 20 overlapping patient samples. Cell types were harmonized between platforms based on shared marker expression profiles: tubular epithelial cells include all PT states (PT-H, PT-dd, PT-INJ), thick ascending limb (TAL), distal convoluted tubule (DCT), collecting duct (CD), connecting tubule (CNT), dedifferentiated tubular epithelium (ddTE); endothelial compartment comprises endothelial cells (EC) and lymphatic endothelial cells (LEC); stromal populations include fibroblasts, myofibroblasts, pericytes, vascular smooth muscle cells, general stromal cells and Mixed_VIM_high populations; immune cells encompass all lymphoid and myeloid populations including T cells, B cells, plasma cells, macrophages, dendritic cells, mast cells, and eosinophils. Data presented as mean cell density ± 95% confidence intervals calculated from patient-level measurements. Y-axis displayed on log10 scale. Numbers above bars indicate mean density values. Strong concordance observed between platforms for endothelial (IMC: 357 vs Xenium: 558 cells/mm²) and immune compartments (IMC: 1064 vs Xenium: 1781 cells/mm²). Tubular epithelial densities showed excellent agreement (IMC: 2167 vs Xenium: 2712 cells/mm²). Platform-specific differences in stromal density (IMC: 508 vs Xenium: 1650 cells/mm²) likely reflect differential performance in cell segmentation of stromal cell populations and differential sensitivity to stromal marker detection between protein-based (IMC) and transcript-based (Xenium) platforms. Overall concordance validates the reliability of cell type annotations and supports cross-platform integration of spatial analyses.

**Supplementary Tables (attached separately):**

**Supplementary Table 1 | Discovery cohort IMC batch breakdown and reference tissue injury scoring.** Imaging mass cytometry (IMC) experimental batches for the discovery cohort showing temporal distribution of sample processing and pathologist-adjudicated injury scores for reference tissues. Each batch contained one patient sample from each disease category (AIN, ATI, and reference tissue) to minimize batch effects, with exceptions in batches #9 and #12 where ATI samples were unavailable. Reference tissues were obtained from tumor nephrectomy specimens distant from tumor site (tumor remote) or from living kidney donors. Tissue injury was assessed by a blinded renal pathologist on adjacent H&E-stained sections using standardized scoring criteria for interstitial fibrosis and tubular atrophy (IFTA), interstitial chronic inflammation (ICI), and acute tubular injury (ATI). Scores were reported as percentage of cortical area affected using categorical ranges (<5%, 6-25%, 26-50%, >50%). For statistical analysis, categorical ranges were converted to continuous variables using range midpoints (2.5%, 15.5%, 38%, and 75%, respectively), with summary statistics presented as mean ± standard deviation. This conversion assumes uniform distribution within each range and may underestimate variance, particularly for open-ended categories. The low injury scores across reference tissues (IFTA: 4.0 ± 5.2%, ICI: 2.5 ± 0.0%, ATI: 4.4 ± 5.3%) confirm minimal background pathology in control samples. Antibody hybridization and IMC data acquisition were performed on consecutive days to ensure optimal signal quality. n=22 AIN, n=21 ATI, n=22 reference tissues across 23 batches. 1 AIN and 2 ATI samples were not included due to technical disruptions in CyTOF software causing loss of data. Reference tissue used in batch 7 was a unique section from the same patient tissue used for reference tissue in batch 1, and thus was not counted as a unique patient.

**Supplementary Table 2 | Validation cohort IMC batch breakdown and reference tissue injury scoring.** Imaging mass cytometry (IMC) experimental batches for the validation cohort showing temporal distribution of sample processing and pathologist-adjudicated injury scores for reference tissues. Batches #1-10 contained one patient sample from each disease category (AIN, ATI, and reference tissue) to minimize batch effects, with exception of batch #8 where AIN sample was unavailable. Batches #11-16 contained AIN and ATI samples paired with additional tissue sections from reference patients previously analyzed in earlier batches (indicated in parentheses) to further control for batch effects and enable cross-batch normalization. All reference tissues were obtained from tumor nephrectomy specimens distant from tumor site. Tissue injury was assessed by a blinded renal pathologist on adjacent H&E-stained sections using standardized scoring criteria for interstitial fibrosis and tubular atrophy (IFTA), interstitial chronic inflammation (ICI), and acute tubular injury (ATI). Scores were reported as percentage of cortical area affected using categorical ranges (<5%, 6-25%, 26-50%, >50%). For statistical analysis, categorical ranges were converted to continuous variables using range midpoints (2.5%, 15.5%, 38%, and 75%, respectively), with summary statistics presented as mean ± standard deviation. This conversion assumes uniform distribution within each range and may underestimate variance, particularly for open-ended categories. The validation reference tissues showed higher baseline injury scores (IFTA: 8.7 ± 11.6%, ICI: 6.4 ± 6.3%, ATI: 29.1 ± 20.7%) compared to discovery cohort reference tissues (Supplementary Table 1), likely reflecting underlying kidney disease in this patient population. Antibody hybridization and IMC data acquisition were performed within 2-5 days to ensure optimal signal quality. n=15 AIN (1 excluded due to sample duplication), n=16 ATI, n=10 unique reference tissue patients across 16 batches.

**Supplementary Table 3 | IMC antibody panel for discovery and validation cohorts**. Complete antibody panel used for imaging mass cytometry (IMC) analysis across discovery and validation cohorts. The 35-antibody panel comprises cell type-defining markers (non-underlined bolded targets), functional markers for cell activation states, injury biomarkers, and signaling molecules (underlined bolded targets), and markers for cell segmentation (regular font). Metal-conjugated antibodies were either purchased pre-conjugated from Standard BioTools (indicated by asterisk on metal ion) or conjugated in-house using lanthanide metals and MaxPar conjugation kits (Standard BioTools) following manufacturer's protocols. Clone information is provided for monoclonal antibodies; polyclonal antibodies are indicated by catalog numbers in brackets. Dilutions represent final working concentrations optimized for kidney tissue staining. Prior IMC validation in kidney tissue is indicated by PubMed identifiers (PMIDs) where these specific metal-conjugated antibodies were previously published. Additional validation methods for antibodies not previously published in kidney IMC include: knockout (KO) validation per supplier specifications, immunofluorescence (IF) validation performed in our laboratory following protocols detailed in Methods, and well-referenced (R) status indicating >20 publications using the specific clone. All antibodies underwent vendor validation prior to purchase with preference given to knockout-validated monoclonal antibodies when available. Post-conjugation validation was performed for all antibodies using IMC on kidney tissue sections containing expected target antigens. The panel enables comprehensive single-cell phenotyping of kidney-resident cells (tubular epithelium segments, podocytes, endothelial cells, fibroblasts), immune cells (T cells, B cells, macrophages, dendritic cells, eosinophils, neutrophils, mast cells), and functional states (injury markers KIM-1 and VCAM-1, proliferation marker Ki-67, cytokines IL-9 and TNF-α, cellular processes including ferroptosis, autophagy, and chemotaxis). DNA intercalators (191Ir, 193Ir) and cell membrane markers (195Pt, 196Pt) enable accurate nuclear and cellular segmentation, respectively. Abbreviations: TE, tubular epithelium; SB, Standard BioTools; BL, BioLegend; EM, EMD Millipore; RD, R&D Systems; TF, ThermoFisher Scientific; NB, Novus Biologicals; CS, Cell Signaling Technology; KO, knockout validated; IF, immunofluorescence validated; R, well-referenced clone (>20 publications).

**Supplementary Table 4 | IMC cluster definitions for discovery and validation cohorts**. Comprehensive cell type definitions derived from unsupervised clustering of imaging mass cytometry (IMC) data from 106 kidney biopsies (discovery cohort n=65, validation cohort n=41). A total of 49 distinct cellular clusters were identified based on protein expression patterns and spatial localization, which were subsequently grouped into 20 core cell types (including two mixed cell types) for UMAP visualization. Cluster assignments were determined through hierarchical clustering using marker expression profiles, with refinement based on spatial context and morphological features. Tubular epithelial cells were classified into specialized segments and functional states: proximal tubule (PT) cells included healthy (PT-H), dedifferentiated (PT-dd), two injury phenotypes (PT-INJ1 characterized by KIM-1 positivity; PT-INJ2 by VCAM-1 positivity), and proliferating (PT-Prolif) states. Similar functional heterogeneity was observed across thick ascending limb (TAL), distal convoluted tubule (DCT), and collecting duct (CD) populations, with additional FACL4$^+$ subsets indicating ferroptosis-associated states. Dedifferentiated tubular epithelial cells (ddTE) represented cells with loss of segment-specific markers while maintaining epithelial identity through β-catenin expression. Immune populations were defined by lineage markers and activation states, including CD4$^+$ and CD8$^+$ T cell subsets with IL-9$^+$ variants, mononuclear phagocytes (MP) stratified by CD206 expression (M2-like polarization), mast cells, and eosinophils. Mixed immune clusters likely represent areas of dense immune infiltration where individual cell boundaries could not be fully resolved at IMC resolution. Stromal compartments included endothelial cells (ECs), lymphatic endothelial cells (LECs), vascular smooth muscle cells, and stromal fibroblasts. Glomerular cells (podocytes, mesangial, parietal) were identified through combined marker expression and spatial localization within glomerular structures. Undefined clusters (UI, uIS, uTE) representing rare cells with ambiguous marker profiles or technical artifacts were excluded from downstream analyses. Distinguishing markers listed represent proteins with differential expression defining each cluster; markers in brackets indicate spatial or morphological features used for cluster refinement. Expression levels noted as "low" indicate reduced but detectable signal compared to corresponding high-expression clusters. Abbreviations: PT, proximal tubule; tDL, thin descending limb; TAL, thick ascending limb; DCT, distal convoluted tubule; CNT, connecting tubule; CD, collecting duct; ddTE, dedifferentiated tubular epithelium; Podo, podocytes; ECs, endothelial cells; LECs, lymphatic endothelial cells; MP, mononuclear phagocytes; Eos, eosinophils; UI, unidentified; uIS, undefined interstitial; uTE, undefined tubular epithelium; Prolif, proliferating.

**Supplementary Table 5 | Spatial transcriptomics slide layout and tissue types included in analysis.** Layout of tissue sections across four 10x Genomics Xenium slides for spatial transcriptomics analysis. Each slide accommodated up to 6 tissue sections arranged side by side. Tissues selected from the IMC discovery cohort (batch numbers cross-referenced with **Supplementary Table 1**) to enable direct spatial proteomic-transcriptomic comparison within the same patient samples. Analysis included both cortical and medullary regions when available. Of 20 planned tissue sections (8 AIN, 7 ATI, 5 reference), 16 were successfully analyzed. Four sections (1 ATI, 3 reference) were lost during Xenium processing due to inadequate tissue adhesion during the multi-day workflow, indicated by 'X' in the table. The ATI sample from batch #19 and reference samples from batches #6, #17, and #18 detached from slides during enzymatic permeabilization or washing steps and could not be recovered for analysis. Final spatial transcriptomics dataset comprised 18 unique cortical sections and 4 medullary sections from 16 patients (8 AIN, 6 ATI, 2 reference), generating 321,333 cells with single-cell resolution gene expression profiles for 5,101 genes. Cortex-medulla tissue pairs were available for 3 patients (AIN batch #9, ATI batch #3, reference batch #6), enabling regional heterogeneity assessment. All tissues underwent quality control validation through H&E staining and cell segmentation verification prior to downstream analysis.

**Supplementary Table 6 | Cell type definitions and marker expression in spatial transcriptomics analysis.** Comprehensive annotation of 27 distinct core cell populations (including one mixed population) identified through unsupervised clustering of 321,318 cells from spatial transcriptomics analysis of 20 kidney biopsies (8 AIN, 7 ATI, 5 reference). Cell types were defined using the 10x Genomics Xenium Prime 5K Human Pan Tissue & Pathways Panel plus 100 custom kidney-specific genes. Percentages indicate proportion of cells within each cluster expressing the marker above background threshold. Categories reflect anatomical compartments and functional groupings: Glomerular (podocytes, parietal epithelial cells), Proximal tubule (healthy, dedifferentiated, and injured states), Loop of Henle, Distal nephron (DCT, CNT, CD), general Tubular (dedifferentiated epithelium), Vascular (endothelial cells, lymphatic endothelium), Stromal (fibroblasts, pericytes, vascular smooth muscle, myofibroblasts), Lymphoid (T cell subsets, B cells, plasma cells), Myeloid (macrophages, dendritic cells, mast cells, eosinophils), Specialized (juxtaglomerular apparatus, glomerular stromal), and Mixed populations representing areas of dense cellular infiltration or intermediate differentiation states. Notable findings include: (1) Three distinct proximal tubule states with PT-INJ showing highest prevalence (20,849 cells) characterized by injury markers HAVCR1/VCAM1/SPP1; (2) Thick ascending limb representing the largest single population (56,887 cells) with preserved UMOD expression; (3) Eosinophils (271 cells) lacking detectable transcripts at single-cell resolution, identified retrospectively through H&E morphology and spatial registration with segmentation masks, highlighting platform limitations for detecting cells with low mRNA content; (4) Mixed immune population (21,586 cells) expressing pan-leukocyte markers (PTPRC/CD74) in areas of dense inflammation where individual cell resolution was challenging. Markers shown represent the most discriminative features for each population, though classification incorporated expression patterns across all 5,101 genes analyzed.

**Supplementary Table 7 | Molecular characterization of proximal tubule cell states through unsupervised subclustering in spatial transcriptomics.** Results of unsupervised clustering analysis of 90,336 pooled proximal tubule cells from spatial transcriptomics data across all tissue types (8 AIN, 7 ATI, 5 reference, as well as 15 diabetic kidney disease samples not included in this project). Twenty-one initial clusters were identified through graph-based clustering and consolidated into three biologically meaningful states based on marker expression patterns. Final annotations: PT-H (healthy; n=30,124 cells) characterized by high expression of functional transporters LRP2 (megalin, 58.7-98.7% positive), CUBN (cubilin, 48.9-99.8% positive), and low injury markers (HAVCR1 <11%, VCAM1 <16%). PT-dd (dedifferentiated; n=5,850 cells) showing intermediate transporter expression (LRP2 27.8-38.6%, CUBN 9.6-25.2%) with minimal injury markers. PT-INJ (injured; n=20,849 cells) exhibiting variable injury marker expression including HAVCR1 (KIM-1, 11.5-37.8% positive), VCAM1 (16.2-42% positive), SPP1 (osteopontin, 53.8-98.8% positive), and inflammatory markers CD74 (22.4-74.7% positive) and C3 (complement C3, 18.3-55% positive). Expression values represent mean normalized counts per cell. Percentage positive indicates proportion of cells with detectable expression above background. The heterogeneity within PT-INJ clusters (subclusters 0, 2, 6, 7, 10, 13, 14, 15, 18, 20) likely reflects a spectrum of injury severity and temporal evolution, with highest injury marker expression in subclusters 10 (VCAM1 42% positive), 14 (HAVCR1 36.4% positive), and 15 (HAVCR1 37.8% positive). Subcluster 20 uniquely showed high LCN2 (lipocalin-2, 17.5% positive) with minimal VCAM1, suggesting a distinct injury phenotype. These molecular profiles validate the IMC-based PT state classifications and provide transcriptional evidence for distinct injury trajectories in acute kidney injury.

**Supplementary Table 8 | Validation of IMC cell segmentation accuracy using napari The Segmentation Game.** F1-scores for cell segmentation quality assessment across different kidney disease states and reference tissue types. Manual ground truth annotations were created and compared to automated Mesmer segmentation masks using napari The Segmentation Game framework. F1-scores calculated as the harmonic mean of precision and recall: 2 × (precision × recall)/(precision + recall), where precision represents correctly identified cells/total identified cells and recall represents correctly identified cells/total true cells. Scores range from 0 (poor) to 1 (perfect) segmentation, with values approaching 1 indicating high concordance between automated segmentation and ground truth. Analysis included 15 tissue samples with modified Mesmer parameters optimized for kidney tissue (resolution: 0.8, small object threshold: 53, maxima threshold: 0.3, interior threshold: 0.47, nuclear segmentation with 1-pixel expansion). Three samples evaluated per disease category: chronic kidney disease (CKD), acute kidney injury (AKI), diabetic kidney disease (DKD), tumor remote nephrectomy reference, and living donor biopsy reference. Each row represents an individual tissue sample F1-score. Mean F1-score of 0.838 ± 0.028 (SD) across all samples indicates robust segmentation performance with minimal variation between disease states, validating the reliability of downstream single-cell analyses. This performance is comparable to Mesmer's reported accuracy in other tissue types and superior to many alternative segmentation algorithms.

**Supplementary Table 9 | Custom gene panel for 10x Genomics Xenium spatial transcriptomics analysis.** List of 100 project-specific genes added to the standard Xenium Prime 5K Human Pan Tissue & Pathways Panel (5,001 genes) for enhanced characterization of kidney cell types, immune infiltrates, and disease states relevant to acute interstitial nephritis. Custom panel designed to capture: (1) kidney segment-specific transporters and markers (SLC transporters, *AQP2, FXYD4, MIOX, DPEP1*), (2) glomerular proteins (*NPHS1, NPHS2, EMCN*), (3) injury and inflammation markers (*SPP1, LCN2, C3, NAMPT, GADD45B*), (4) immune cell markers and chemokines (*CCL2, CCL4, CCL21, CD74, PTPRC, IL7R, CD69, CD9*), (5) extracellular matrix and fibrosis genes (*COL1A1, COL1A2, COL12A1, LUM, LTBP4, TAGLN*), (6) metabolic and stress response genes (*GPX3, GATM, SOD2, CRYAB, MIF*), and (7) cell state and dedifferentiation markers (*VIM, APOE, IGFBP7*). Probesets indicate the number of independent probe designs targeting each transcript, with redundancy providing increased detection sensitivity. Codewords represent unique molecular barcodes assigned to each gene for multiplexed detection. Gene selection based on: differential expression analysis from prior single-cell RNA sequencing studies in kidney disease, IMC antibody panel targets requiring transcriptional validation, and literature-curated markers of kidney injury, repair, and immune-mediated pathology. The combined 5,101-gene panel enabled comprehensive single-cell spatial transcriptomic profiling across 321,318 cells from 16 kidney biopsies, facilitating identification of 27 distinct core cell types (including one mixed population) and disease-specific transcriptional states. Ensembl IDs provided for unambiguous gene identification.
