## Supplementary Tables for "Spatial analysis reveals the cellular microenvironments and mechanisms of inflammation and kidney injury in acute interstitial nephritis"

**Supplementary Table 1:** Discovery cohort IMC batch breakdown and reference tissue injury scoring

| | Ab hybridization | IMC burn | AIN | ATI | Reference | | Reference Tissue Scoring (%) | | |
|---|---|---|---|---|---|---|---|---|---|
| **Batch** | **Date** | **Date** | **# Patients** | **# Patients** | **# Patients** | **Tissue Type** | **IFTA** | **ICI** | **ATI** |
| **#1** | 3/15/23 | 3/17/23 | *0* | 1 | 1 | Tumor remote | <5 | <5 | <5 |
| **#2** | 4/4/23 | 4/5/23 | 1 | 1 | 1 | Living donor | <5 | <5 | <5 |
| **#3** | 4/20/23 | 4/25/23 | 1 | 1 | 1 | Living donor | <5 | <5 | <5 |
| **#4** | 4/27/23 | 4/28/23 | 1 | 1 | 1 | Living donor | <5 | <5 | <5 |
| **#5** | 5/9/23 | 5/10/23 | 1 | 1 | 1 | Living donor | <5 | <5 | <5 |
| **#6** | 5/9/23 | 5/11/23 | 1 | 1 | 1 | Living donor | <5 | <5 | 10 |
| **#7** | 6/6/23 | 6/9/23 | 1 | 1 | *(Batch 1 patient)* | Tumor remote | | | |
| **#8** | 6/6/23 | 6/8/23 | 1 | 1 | 1 | Tumor remote | <5 | <5 | <5 |
| **#9** | 8/8/23 | 8/9/23 | 1 | *0* | 1 | Tumor remote | <5 | <5 | 20 |
| **#10** | 8/8/23 | 8/10/23 | 1 | 1 | 1 | Tumor remote | <5 | <5 | <5 |
| **#11** | 12/7/23 | 12/9/23 | 1 | 1 | 1 | Tumor remote | <5 | <5 | <5 |
| **#12** | 12/7/23 | 12/11/23 | 1 | *0* | 1 | Tumor remote | 13 | <5 | <5 |
| **#13** | 12/12/23 | 12/13/23 | 1 | 1 | 1 | Tumor remote | <5 | <5 | 20 |
| **#14** | 12/12/23 | 12/15/23 | 1 | 1 | 1 | Tumor remote | <5 | <5 | <5 |
| **#15** | 1/11/24 | 1/12/24 | 1 | 1 | 1 | Living donor | 25 | <5 | <5 |
| **#16** | 1/11/24 | 1/13/24 | 1 | 1 | 1 | Living donor | <5 | <5 | <5 |
| **#17** | 1/16/24 | 1/17/24 | 1 | 1 | 1 | Living donor | <5 | <5 | <5 |
| **#18** | 1/16/24 | 1/18/24 | 1 | 1 | 1 | Living donor | <5 | <5 | <5 |
| **#19** | 1/31/24 | 2/1/24 | 1 | 1 | 1 | Living donor | <5 | <5 | <5 |
| **#20** | 2/6/24 | 2/7/24 | 1 | 1 | 1 | Living donor | <5 | <5 | <5 |
| **#21** | 1/31/24 | 2/2/24 | 1 | 1 | 1 | Living donor | <5 | <5 | <5 |
| **#22** | 2/6/24 | 2/8/24 | 1 | 1 | 1 | Living donor | <5 | <5 | <5 |
| **#23** | 1/31/24 | 2/4/24 | 1 | 1 | 1 | Tumor remote | <5 | <5 | <5 |
| **TOTAL** | | | **22** | **21** | **22** | | **4.0 ± 5.2** | **2.5 ± 0.0** | **4.4 ± 5.3** |

**Supplementary Table 2**: Validation cohort IMC batch breakdown and reference tissue injury scoring

| | Ab hybridization | IMC burn | AIN | ATI | Reference | | Reference Tissue Scoring (%) | | |
|---|---|---|---|---|---|---|---|---|---|
| **Batch** | **Date** | **Date** | **# Patients** | **# Patients** | **# Patients** | **Tissue Type** | **IFTA** | **ICI** | **ATI** |
| **#1** | 8/20/24 | 8/22/24 | 1 | 1 | 1 | Tumor remote | <5 | <5 | 26-50 |
| **#2** | 8/20/24 | 8/24/24 | 1 | 1 | 1 | Tumor remote | <5 | <5 | 6-25 |
| **#3** | 9/2/24 | 9/4/24 | 1 | 1 | 1 | Tumor remote | <5 | <5 | <5 |
| **#4** | 9/2/24 | 9/6/24 | 1 | 1 | 1 | Tumor remote | <5 | 6-25 | 26-50 |
| **#5** | 9/5/24 | 9/7/24 | 1 | 1 | 1 | Tumor remote | <5 | <5 | 26-50 |
| **#6** | 9/5/24 | 9/9/24 | 1 | 1 | 1 | Tumor remote | 6-25 | <5 | 6-25 |
| **#7** | 10/2/24 | 10/7/24 | 1 | 1 | 1 | Tumor remote | 6-25 | 6-25 | 6-25 |
| **#8** | 10/2/24 | 10/8/24 | *0* | 1 | 1 | Tumor remote | 26-50 | 6-25 | >50 |
| **#9** | 10/9/24 | 10/11/24 | 1 | 1 | 1 | Tumor remote | <5 | <5 | 6-25 |
| **#10** | 10/9/24 | 10/12/24 | 1 | 1 | 1 | Tumor remote | <5 | <5 | 26-50 |
| **#11** | 10/15/24 | 10/17/24 | 1 | 1 | (additional tissue from sample used in batch #5) | | | | |
| **#12** | 10/15/24 | 10/18/24 | 1 | 1 | (additional tissue from sample used in batch #7) | | | | |
| **#13** | 11/4/24 | 11/6/24 | 1 | 1 | (additional tissue from sample used in batch #10) | | | | |
| **#14** | 11/4/24 | 11/7/24 | 1 | 1 | (additional tissue from sample used in batch #5) | | | | |
| **#15** | 11/11/24 | 11/13/24 | 1 | 1 | (additional tissue from sample used in batch #9) | | | | |
| **#16** | 11/11/24 | 11/14/24 | 1 | 1 | (additional tissue from sample used in batch #7) | | | | |
| **TOTAL:** | | | **15** | **16** | **10** | | **8.7 ± 11.6** | **6.4 ± 6.3** | **29.1 ± 20.7** |

**Supplementary Table 3:** IMC antibody panel for discovery and validation cohorts

| Target | Antigen/probe | Clone | Metal ion | Supplier | Dilution | Uses in IMC (PMIDs of references) | Validation |
|---|---|---|---|---|---|---|---|
| ***general TE*** | Beta catenin | D10A8 | $^{147}$Sm* | SB | 0.389 | 31217358 | |
| ***proximal TE*** | Aquaporin-1 | EPR11588 | $^{173}$Yb | Abcam | 1.778 | 31217358, 34476295 | |
| ***proximal TE*** | Megalin | 10D5.1 | $^{174}$Yb | EM | 0.215 | 31217358, 34476295 | |
| ***thick ascending limb TE*** | Uromodulin | 774056 | $^{151}$Eu | RD | 1.153 | 31217358, 34476295 | |
| ***distal convoluted tubule TE*** | Calbindin | 401025 | $^{142}$Nd | TF | 0.319 | 31217358, 34476295 | |
| ***collecting duct TE*** | Cytokeratin 7 | RCK105 | $^{164}$Dy* | SB | 0.146 | 31217358, 34476295 | |
| ***podocytes*** | Nestin | 196908 | $^{146}$Nd | Abcam | 0.181 | 31217358, 34476295 | |
| ***fibroblasts, mesangial cells, parietal cells, podocytes*** | Vimentin | RV202 | $^{150}$Nd | Abcam | 0.319 | 34476295 | KO, R, IF |
| ***endothelial cells*** | CD31 | JC/70A | $^{149}$Sm | Abcam | 0.111 | 31217358, 34476295 | |
| ***endothelial cells*** | ETS related gene (ERG) | EPR3864 | $^{166}$Er | Abcam | 0.389 | | R, IF |
| ***lymphatic endothelial cells*** | Podoplanin | D2-40 | $^{154}$Sm | BL | 0.020 | | R, IF |
| ***vascular smooth muscle cells, myofibroblasts*** | α-Smooth muscle actin | 1A4 | $^{141}$Pr* | SB | 0.736 | 31217358, 34476295 | |
| ***mononuclear phagocytes*** | CD68 | KP1 | $^{159}$Tb* | SB | 0.597 | 31217358, 34476295 | |
| ***anti-inflammatory macrophages*** | CD163 | EDHu-1 | $^{148}$Nd | Bio-Rad | 0.111 | | R, IF |
| ***alternatively actived macrophages*** | CD206 | [ab64693] | $^{163}$Dy | Abcam | 0.319 | | R, IF |
| ***dendritic cell*** | CD11c | EP1347Y | $^{167}$Er | Abcam | 0.181 | | R, IF |
| ***T cell*** | CD3 | C3e/1308 | $^{170}$Er | NB | 0.215 | | R, IF |
| ***T helper cells*** | CD4 | EPR6855 | $^{156}$Gd* | SB | 0.111 | 31217358, 34476295 | |
| ***cytotoxic T cells*** | CD8a | 144B | $^{162}$Dy* | SB | 0.250 | 31217358, 34476295 | |
| ***B cells*** | CD20 | H1 | $^{161}$Dy* | SB | 0.146 | 31217358, 34476295 | |
| ***eosinophils*** | Major basic protein | BMK-13 | $^{143}$Nd | NB | 0.056 | | R, IF |
| ***neutrophils*** | Myeloperoxidase | EPR17996 | $^{172}$Yb | Abcam | 0.389 | | R, IF |
| ***mast cells*** | Chymase | EPR13136 | $^{165}$Ho | Abcam | 0.319 | | R, IF |
| ***kidney epithelial cell injury*** | Kidney injury molecule-1 | 219211 | $^{160}$Gd | NB | 0.250 | 34476295 | R, IF |
| ***kidney epithelial cell injury*** | Vascular cell adhesion molecule 1 | EPR5047 | $^{152}$Sm | Abcam | 0.181 | | KO, R, IF |
| ***cell cycle, proliferation*** | Ki67 | B56 | $^{168}$Er* | SB | 0.111 | 34476295 | IF |
| ***Th2-type cytokine*** | Interleukin 9 | [ab181397] | $^{153}$Eu | Abcam | 0.215 | | IF |
| ***ferroptosis*** | Fatty acid-CoA ligase 4 | EPR8640 | $^{155}$Gd | Abcam | 0.319 | | R, IF |
| ***macrophage recruitment*** | Monocyte chemoattractant protein-1 | 2D8 | $^{169}$Tm | NB | 0.250 | | R, IF |
| ***inflammatory cytokine*** | Tumor necrosis factor-α | EPR19147 | $^{145}$Nd | Abcam | 0.250 | | KO, R, IF |
| ***autophagy*** | Light chain 3 beta (LC3b) | EPR18709 | $^{158}$Gd | Abcam | 0.181 | | KO, R, IF |
| Nuclei | DNA intercalator | N/A | $^{191}$Ir | SB | 0.736 | | IMC |
| Nuclei | DNA intercalator | N/A | $^{193}$Ir | SB | 0.736 | | IMC |
| Cell membrane phospholipids | Cell segmentation kit | N/A | $^{195}$Pt | SB | 0.319 | | IMC |
| Cell membrane phospholipids | Cell segmentation kit | N/A | $^{196}$Pt | SB | 0.319 | | IMC |

**Supplementary Table 4**: IMC Cluster definitions (discovery and validation cohorts)

| **Abbreviated cluster name** | **Core cell type (UMAP)** | **Full cluster name** | **Distinguishing markers** |
|---|---|---|---|
| PT-H | PT | Healthy PT | Beta-catenin, Megalin, Aquaporin 1 |
| PT-dd | PT | De-differentiated PT | Beta-catenin, Megalin (low), Aquaporin 1 (low) |
| PT-INJ1 | PT | Injured PT 1 | Beta-catenin, Megalin, Aquaporin 1, KIM-1 |
| PT-INJ2 | PT | Injured PT 2 | Beta-catenin, Megalin, Aquaporin 1, VCAM-1 |
| PT-Prolif | PT | Proliferating PT | Beta-catenin, Megalin, Aquaporin 1, Ki67 |
| tDL | tDL | Thin descending limb of the Loop of Henle | Beta-catenin, Aquaporin 1 |
| TAL-H | TAL | Thick ascending limb of the Loop of Henle with high uromodulin expression | Beta-catenin, Uromodulin |
| TAL-dd | TAL | Thick ascending limb of the Loop of Henle with low uromodulin expression | Beta-catenin, Uromodulin (low) |
| TAL-FACL4+ | TAL | Thick ascending limb of the Loop of Henle with high FACL4 expression | Beta-catenin, Uromodulin, FACL4 |
| TAL-Prolif | TAL | Proliferating thick ascending limb of the Loop of Henle | Beta-catenin, Uromodulin, Ki67 |
| DCT-Calb-hi | DCT | Distal convoluted tubule with high calbindin expression | Beta-catenin, Calbindin |
| DCT-Calb-low | DCT | Distal convoluted tubule with low calbindin expression | Beta-catenin, Calbindin (low) |
| DCT-FACL4+ | DCT | Distal convoluted tubule with high FACL4 expression | Beta-catenin, Calbindin, FACL4 |
| DCT-Prolif | DCT | Proliferating distal convoluted tubule | Beta-catenin, Calbindin, Ki67 |
| CNT | CNT | Connecting tubule | Beta-catenin, Calbindin, Cytokeratin 7 |
| CD-CK7-hi | CD | Collecting duct with high cytokeratin 7 expression | Beta-catenin, Cytokeratin 7 |
| CD-CK7-low | CD | Collecting duct with low cytokeratin 7 expression | Beta-catenin, Cytokeratin 7 (low) |
| CD-FACL4+ | CD | Collecting duct with high FACL4 expression | Beta-catenin, Cytokeratin 7, FACL4 |
| CD-Prolif | CD | Proliferating collecting duct | Beta-catenin, Cytokeratin 7, Ki67 |
| ddTE | ddTE | De-differentiated tubular epithelial cells | Beta-catenin |
| ddTE-FACL4+ | ddTE | De-differentiated tubular epithelial cells with high FACL4 | Beta-catenin, FACL4 |
| Podo | Podo | Podocytes | Nestin |
| Podo & Mesangial cells | (Mixed) Glomerular Stromal | Mixed podocytes and mesangial cells | Nestin, a-SMA |
| Podo & Vascular | (Mixed) Glomerular Stromal | Mixed podocytes and vascular cells | Nestin, [location inside glomerulus and outside glomerulus around endothelial cells] |
| Parietal | Parietal | Parietal cells | Vimentin, Beta-catenin [location lining Bowman's capsule] |
| ECs | ECs | Endothelial cells | ERG, CD31 |
| LECs | LECs | Lymphatic endothelial cells | M2a antigen, ERG, CD31 |
| Vascular | Vascular | Vascular smooth muscle cells | Nestin, [location outside glomerulus around endothelial cells] |
| Stromal | Stromal | Stromal cells including fibroblasts | a-SMA |
| Stromal & Vascular | (Mixed) Stromal | Mixed stromal cells and vascular cells | a-SMA, Nestin, [location outside glomerulus around endothelial cells] |
| CD4/8-low/- T cells | T cells | T cells with very low CD4 and CD8 expression | CD3 |
| CD4/8-low/- T-Prolif | T cells | Proliferating T cells with very low CD4 and CD8 expression | CD3, Ki67 |
| CD4+ T cells | T cells | CD4+ helper T cells | CD3, CD4 |
| CD4+ T-IL9+ | T cells | CD4+ helper T cells with high IL9 expression | CD3, CD4, IL9 |
| CD4+ T-Prolif | T cells | Proliferating CD4+ helper T cells | CD3, CD4, Ki67 |
| CD8+ T cells | T cells | CD8+ cytotoxic T cells | CD3, CD8 |
| CD8+ T-IL9+ | T cells | CD8+ cytotoxic T cells with high IL9 expression | CD3, CD8, IL9 |
| CD8+ T-Prolif | T cells | Proliferating CD8+ cytotoxic T cells | CD3, CD8, Prolif |
| Mixed B & T cells | B+T cells | Mixed B cells and T cells | CD3, CD20 |
| Mixed B & T-Prolif cells | B+T cells | Proliferating mixed B cells and T cells | CD3, CD20, Ki67 |
| MP-CD206+ | MP | Mononuclear phagocytes with high CD206 expression | CD68, CD206 |
| MP-CD206+-Prolif | MP | Proliferating mononuclear phagocytes with high CD206 expression | CD68, CD206, Ki67 |
| MP-CD206- | MP | Mononuclear phagocytes | CD68 |
| MP & T cells | Mixed Immune | Mixed mononuclear phagocytes and T cells | CD68, CD163, CD206, CD3, CD4, CD8 |
| MP & T cells-IL9+ | Mixed Immune | Mixed mononuclear phagocytes and T cells with high IL9 expression | CD68, CD163, CD206, CD3, CD4, CD8, IL9 |
| MP & T cells-Prolif | Mixed Immune | Proliferating mixed mononuclear phagocytes and T cells | CD68, CD163, CD206, CD3, CD4, CD8, Ki67 |
| MP & CD8+ T cells | Mixed Immune | Mixed mononuclear phagocytes and CD8+ T cells | CD68, CD163, CD206, CD3, CD8 |
| Mast | Mast | Mast cells | Chymase, CD68 |
| Eos | Eosinophils | Eosinophils | Major Basic Protein |
| UI | Undefined (excluded) | Unidentified cells | [Extreme values, artifactual appearance] |
| uIS | Undefined (excluded) | Undefined interstitial cells | [No interstitial cell markers, distinct cells, interstitial location] |
| uTE | Undefined (excluded) | Undefined tubular epithelial cells | [No tubular cell markers, distinct cells, tubular location] |
| uIS-Prolif | Undefined (excluded) | Proliferating undefined interstitial cells | Ki67, [No interstitial cell markers, distinct cells, interstitial location] |
| uTE-Prolif | Undefined (excluded) | Proliferating undefined tubular epithelial cells | Ki67, [No tubular cell markers, distinct cells, tubular location] |

**Supplementary Table 5**: Spatial transcriptomics slide layout and tissue types included in analysis

| **Slide (right):** | **1** | | | **2** | | | **3** | | | **4** | | |
|---|---|---|---|---|---|---|---|---|---|---|---|---|
| **Tissue type (below):** | **Discovery batch # of corresponding IMC tissues** | **Cortex** | **Medulla** | **Discovery batch # of corresponding IMC tissues** | **Cortex** | **Medulla** | **Discovery batch # of corresponding IMC tissues** | **Cortex** | **Medulla** | **Discovery batch # of corresponding IMC tissues** | **Cortex** | **Medulla** |
| **AIN** | 3 | 1 | 0 | 6 | 1 | 0 | 9 | 1 | 0 | 11 | 1 | 0 |
| | 2 | 1 | 0 | 5 | 0 | 1 | 8 | 1 | 0 | 4 | 1 | 0 |
| **ATI** | 23 | 1 | 0 | 18 | 1 | 0 | 2 | 1 | 0 | 3 | 1 | 1 |
| | 4 | 1 | 0 | 20 | 1 | 0 | | *X* | *X* | 19 | 1 | 0 |
| **Reference** | 21 | 1 | 0 | | *X* | *X* | 6 | 1 | 1 | 22 | 1 | 0 |
| | 17 | 0 | 1 | | *X* | *X* | | *X* | *X* | 18 | 1 | 0 |
| **Totals** | | **5** | **1** | | **3** | **1** | | **4** | **1** | | **6** | **1** |

**Supplementary Table 6**: Major cell clusters identified in ST and characteristic transcripts

| Category | Core Cell Type (UMAP | Cell Type | Abbreviation | N Cells | Defining Markers (% positive) | Notes |
|---|---|---|---|---|---|---|
| Glomerular | Podocytes | Podocyte | Podo | 2660 | NPHS2 (87.4%); PODXL (76.8%); NPHS1 (55.8%); WT1 (12.3%) | Glomerular visceral epithelial cells |
| Specialized | Glomerular stromal | Glomerular stromal cell | Glomerular_Stromal | 2543 | POSTN (32.5%); EPAS1 (28.5%) | Mesangial and glomerular stromal cells |
| Specialized | JGA | Juxtaglomerular apparatus | JGA | 673 | REN (99.4%); EPAS1 (33.9%) | Renin-producing cells |
| Glomerular | Parietal | Parietal epithelial cell | Parietal | 883 | PAX8 (21.5%); CLDN1 (28.7%); CFH (34.3%) | Bowman's capsule lining cells |
| Proximal tubule | PT | Proximal tubule - injured | PT-INJ | 20849 | HAVCR1 (19.7%); VCAM1 (25.2%); SPP1 (81.1%); LCN2 (3.1%); C3 (37.2%) | Encompasses both KIM-1+ and VCAM1+ injury phenotypes |
| Proximal tubule | PT | Proximal tubule - healthy | PT-H | 13024 | LRP2 (76.9%); CUBN (68.4%); SLC5A2 (13.2%); SLC34A1 (53.4%); SLC5A12 (69.4%) | Fully differentiated, functional transporters preserved |
| Proximal tubule | PT | Proximal tubule - dedifferentiated | PT-dd | 5850 | VIM (22.0%); APOE (45.4%); GPX3 (89.9%); GATM (61.2%) | Loss of differentiation markers, stress response activated |
| Loop of Henle | tDL | Thin descending limb | tDL | 9126 | PKHD1 (66.1%); CRYAB (66.9%) | Water-permeable descending limb |
| Loop of Henle | TAL | Thick ascending limb | TAL | 56887 | UMOD (63.1%); SLC12A1 (65.5%); CASR (32.4%); EGF (39.1%) | Salt reabsorption, Tamm-Horsfall protein production |
| Distal nephron | DCT | Distal convoluted tubule | DCT | 2929 | SLC12A3 (98.4%); CALB1 (17.1%); CA12 (71.0%) | NCC-expressing, thiazide-sensitive segment |
| Distal nephron | CNT | Connecting tubule | CNT | 3216 | SLC8A1 (98.8%); HSD11B2 (41.2%); CALB1 (33.9%) | Transitional segment, aldosterone-sensitive |
| Distal nephron | CD | Collecting duct | CD | 20639 | AQP2 (77.5%); FXYD4 (72.7%); HSD11B2 (63.9%); SCNN1G (45.9%) | Principal and intercalated cells |
| Tubular | ddTE | Dedifferentiated tubular epithelium | ddTE | 2917 | PAX8 (34.0%); CRYAB (27.5%) | Non-specific tubular dedifferentiation |
| Vascular | ECs | Endothelial cell | EC | 28765 | PECAM1 (41.2%); CDH5 (20.1%); FLT1 (35.3%); KDR (23.6%) | Capillary and arteriolar endothelium |
| Vascular | LECs | Lymphatic endothelial cell | LEC | 2142 | LYVE1 (31.5%); PROX1 (4.5%); CCL21 (74.6%) | Lymphatic vessels |
| Stromal | Fibroblasts | Fibroblast | Fibroblasts | 40964 | COL1A1 (64.6%); COL1A2 (60.8%); PDGFRA (26.3%); LUM (51.7%) | Interstitial fibroblasts |
| Stromal | Stromal | Stromal cell | Stromal | 22313 | COL1A2 (37.5%); LUM (41.5%); VIM (66.6%) | General stromal population |
| Stromal | vSMC | Vascular smooth muscle cell | vSMC | 7457 | TAGLN (58.2%); CALD1 (33.6%) | Arteriolar smooth muscle |
| Stromal | Pericyte | Pericyte | Pericyte | 3386 | PDGFRB (56.3%); RGS5 (43.1%); NOTCH3 (58.6%) | Capillary-associated contractile cells |
| Stromal | Myofibroblasts | Myofibroblast | Myofibroblast | 1518 | COL1A1 (20.6%); TAGLN (82.1%) | Activated fibroblasts, pro-fibrotic |
| Lymphoid | T cells | CD8+ T cell | T_cell_CD8 | 10849 | CD3E (35.4%); CD8A (26.2%); GZMB (0.7%) | Cytotoxic T cells |
| Lymphoid | Plasma cells | Plasma cell | Plasma_cell | 8801 | MZB1 (23.7%); IGKC (76.3%); XBP1 (50.4%) | Antibody-producing B cell lineage |
| Lymphoid | B cells | B cell | B_cell | 5165 | MS4A1 (32.9%); CD79A (19.0%); CD19 (17.0%) | Mature B lymphocytes |
| Lymphoid | T cells | Double negative T cell | T_cell_DN | 3636 | CD3E (10.9%); PTPRC (58.9%) | CD4-CD8- T cells |
| Lymphoid | T cells | CD4+ T cell | T_cell_CD4 | 3291 | CD3E (23.1%); CD4 (15.8%); IL7R (21.1%) | Helper T cells |
| Lymphoid | T cells | Low quality T cell | T_cell_LQ | 1906 | CD3E (5.7%); PTPRC (44.8%) | T cells with low transcript detection |
| Lymphoid | T cells | Gamma-delta T cell | T_cell_gd | 302 | CD3E (23.8%); TRDC (18.2%) | γδ T cells, innate-like lymphocytes |
| Myeloid | Macrophages | Macrophage | Macrophage | 9584 | CD68 (27.6%); CD14 (48.4%); MRC1 (43.1%) | Tissue-resident and infiltrating macrophages |
| Myeloid | Dendritic cells | Dendritic cell | Dendritic_cell | 2194 | ITGAX (21.1%); FCER1A (26.6%); CLEC10A (31.1%) | Antigen-presenting cells |
| Myeloid | APCs | Antigen presenting cell | APC | 1227 | CD74 (84.8%); CD86 (0.9%) | Professional antigen presenting cells |
| Myeloid | Mast cells | Mast cell | Mast_cell | 469 | KIT (26.9%); MS4A2 (20.9%) | Tissue-resident mast cells |
| Myeloid | Eosinophils | Eosinophil | Eosinophils | 271 | (None) | Not detected in spatial transcriptomics platform; filtered out as low quality cells with low to no transcript expression. Identified manually through identification on H&E performed on tissue sample after ST, and image alignment with ST segmentation mask. |
| Mixed | Mixed immune cells | Mixed immune population | Immune_mixed | 21586 | PTPRC (29.3%); CD74 (76.9%) | Heterogeneous immune cells |
| Mixed | Stromal | Mixed epithelial - VIM high | Mixed_VIM_high | 3311 | VIM (59.0%); PAX8 (10.5%) | Stromal cells with high vimentin with rare dedifferentiated epithelial cells |

**Supplementary Table 7**: Molecular characterization of proximal tubule cell states through unsupervised subclustering in spatial transcriptomics

| Unsupervised PT subcluster number | Final annotation type | n_cells | HAVCR1_mean | HAVCR1_pct_pos | VCAM1_mean | VCAM1_pct_pos | LCN2_mean | LCN2_pct_pos | VIM_mean | VIM_pct_pos | LRP2_mean | LRP2_pct_pos | CUBN_mean | CUBN_pct_pos | SPP1_mean | SPP1_pct_pos | CD74_mean | CD74_pct_pos | C3_mean | C3_pct_pos | MIF_mean | MIF_pct_pos | ICAM1_mean | ICAM1_pct_pos |
|---|---|---|---|---|---|---|---|---|---|---|---|---|---|---|---|---|---|---|---|---|---|---|---|---|
| 0 | PT-INJ | 14030 | 0.131 | 17.8 | 0.202 | 26.1 | 0.016 | 2.2 | 0.811 | 70.4 | 0.292 | 35.9 | 0.103 | 13.8 | 0.597 | 53.8 | 0.26 | 29.3 | 0.2329 | 25.8 | 0.1745 | 23.3 | 0.013 | 1.8 |
| 1 | PT-H | 12884 | 0.029 | 4.1 | 0.05 | 6.8 | 0.001 | 0.2 | 0.107 | 13 | 0.538 | 58.7 | 0.408 | 48.9 | 0.644 | 59.7 | 0.194 | 23.4 | 0.0222 | 2.9 | 0.1394 | 18.3 | 0.0028 | 0.4 |
| 2 | PT-INJ | 10068 | 0.137 | 18.3 | 0.193 | 24 | 0.011 | 1.4 | 0.89 | 78 | 0.29 | 35.4 | 0.064 | 8.5 | 0.69 | 61.5 | 0.187 | 22.4 | 0.357 | 34.2 | 0.1688 | 23.1 | 0.0113 | 1.5 |
| 3 | PT-H | 8148 | 0.007 | 1 | 0.024 | 3.4 | 0 | 0 | 0.028 | 3.8 | 0.567 | 64.9 | 0.425 | 53.4 | 0.65 | 62.9 | 0.1 | 13.4 | 0.0222 | 2.9 | 0.0947 | 13.4 | 2.00E-04 | 0 |
| 4 | PT-H | 6210 | 0.014 | 2 | 0.089 | 11.8 | 0.001 | 0.1 | 0.103 | 11.9 | 0.825 | 77.4 | 0.718 | 73.8 | 0.755 | 70.9 | 0.287 | 33.5 | 0.026 | 3 | 0.2938 | 36.5 | 0.0048 | 0.7 |
| 5 | PT-dd | 5868 | 0.021 | 2.7 | 0.035 | 4.6 | 0.002 | 0.2 | 0.137 | 16.4 | 0.341 | 38.6 | 0.202 | 25.2 | 0.591 | 52.5 | 0.184 | 21.7 | 0.0195 | 2.6 | 0.0915 | 12.2 | 0.0019 | 0.3 |
| 6 | PT-INJ | 4659 | 0.109 | 14.9 | 0.21 | 26.8 | 0.018 | 2.2 | 0.413 | 42 | 0.722 | 74 | 0.177 | 23.6 | 1.789 | 97 | 0.708 | 74.7 | 0.6352 | 55 | 0.2172 | 29.8 | 0.0111 | 1.5 |
| 7 | PT-INJ | 3792 | 0.151 | 20.2 | 0.203 | 26.2 | 0.032 | 4.1 | 0.516 | 47.6 | 0.596 | 64.5 | 0.205 | 27.2 | 1.799 | 97.2 | 0.725 | 71.4 | 0.4524 | 45.1 | 0.236 | 30.7 | 0.0299 | 4.2 |
| 8 | PT-H | 3202 | 0.002 | 0.3 | 0.04 | 5.4 | 0 | 0 | 0.033 | 4.3 | 0.772 | 79.6 | 0.769 | 81.1 | 0.676 | 71.5 | 0.119 | 15.9 | 0.0064 | 0.9 | 0.2453 | 33.5 | 2.00E-04 | 0 |
| 9 | PT-dd | 2948 | 0.006 | 0.8 | 0.017 | 2.3 | 0 | 0 | 0.077 | 9.1 | 0.243 | 29.2 | 0.113 | 14.4 | 0.465 | 43.7 | 0.097 | 11 | 0.0119 | 1.5 | 0.0624 | 8.2 | 0 | 0 |
| 10 | PT-INJ | 2922 | 0.203 | 26.2 | 0.363 | 42 | 0.015 | 2.1 | 1.497 | 92.3 | 0.582 | 60.6 | 0.235 | 28.5 | 0.859 | 69.1 | 0.511 | 52.7 | 0.2908 | 28.4 | 0.3665 | 44.9 | 0.0248 | 3.5 |
| 11 | PT-dd | 2620 | 0.018 | 2.4 | 0.053 | 7.1 | 0.005 | 0.6 | 0.478 | 45 | 0.287 | 30.8 | 0.236 | 25.2 | 0.699 | 58.5 | 0.625 | 55.6 | 0.0753 | 7.8 | 0.209 | 24.9 | 0.0083 | 1.1 |
| 12 | PT-H | 2438 | 0.036 | 4.8 | 0.08 | 10.2 | 0 | 0 | 0.116 | 12.6 | 1.224 | 90.8 | 1.249 | 91.5 | 0.985 | 84 | 0.394 | 45.4 | 0.015 | 1.9 | 0.4606 | 51.9 | 0.0027 | 0.4 |
| 13 | PT-INJ | 2301 | 0.149 | 20.3 | 0.133 | 16.2 | 0.016 | 2.1 | 0.547 | 49.2 | 0.432 | 47.9 | 0.249 | 26.7 | 0.896 | 73.1 | 0.326 | 39 | 0.1984 | 18.3 | 0.1698 | 22.5 | 0.019 | 2.7 |
| 14 | PT-INJ | 2176 | 0.296 | 36.4 | 0.315 | 32.7 | 0.034 | 4.1 | 0.907 | 66.4 | 0.943 | 75.8 | 0.568 | 52 | 1.548 | 85.3 | 0.394 | 40.2 | 0.085 | 9.1 | 0.5974 | 62.4 | 0.0294 | 4 |
| 15 | PT-INJ | 1649 | 0.305 | 37.8 | 0.155 | 19.3 | 0.033 | 3.9 | 0.784 | 64.3 | 1.088 | 87.4 | 0.643 | 64.3 | 1.921 | 97.2 | 0.273 | 31 | 0.0914 | 9.7 | 0.5176 | 60.8 | 0.009 | 1.2 |
| 16 | PT-H | 1132 | 0.002 | 0.4 | 0.041 | 5.5 | 0 | 0 | 0.036 | 4.6 | 1.202 | 95.6 | 1.409 | 98.1 | 0.891 | 88 | 0.196 | 26 | 0.0018 | 0.3 | 0.4912 | 61.7 | 0 | 0 |
| 17 | PT-H | 1012 | 0.079 | 10.7 | 0.13 | 15.8 | 0 | 0 | 0.29 | 31.8 | 1.894 | 98.7 | 2.199 | 99.8 | 1.633 | 97 | 0.601 | 64.3 | 0.0435 | 5.8 | 1.0471 | 88.8 | 0.0062 | 0.9 |
| 18 | PT-INJ | 991 | 0.151 | 20.2 | 0.274 | 33.1 | 0.005 | 0.7 | 1.449 | 93.5 | 0.669 | 69.5 | 0.292 | 35.6 | 0.96 | 78.5 | 0.419 | 47.6 | 0.2394 | 26.2 | 0.3829 | 49.6 | 0 | 0 |
| 19 | PT-dd | 868 | 0.04 | 5.2 | 0.052 | 6.7 | 0.01 | 1.4 | 0.193 | 23.6 | 0.241 | 27.8 | 0.077 | 9.6 | 0.305 | 29.3 | 0.155 | 17.7 | 0.0892 | 11.1 | 0.0421 | 5.9 | 0.0056 | 0.8 |
| 20 | PT-INJ | 418 | 0.083 | 11.5 | 0.002 | 0.2 | 0.152 | 17.5 | 0.081 | 9.6 | 0.015 | 2.2 | 0.007 | 0.7 | 2.725 | 98.8 | 0.441 | 43.1 | 0.1541 | 14.6 | 0.2861 | 37.6 | 0.0142 | 1.9 |

**Supplementary Table 8**: Validation of IMC cell segmentation accuracy using napari The Segmentation Game

| | **F1-score** |
|---|---|
| CKD | 0.81 |
| | 0.77 |
| | 0.85 |
| Reference tissue: Tumor remote nephrectomy | 0.84 |
| | 0.83 |
| | 0.83 |
| AKI | 0.88 |
| | 0.83 |
| | 0.86 |
| Reference tissue: Living donor biopsy | 0.81 |
| | 0.88 |
| | 0.85 |
| DKD | 0.84 |
| | 0.86 |
| | 0.83 |
| **Mean ± SD:** | **0.838 ± 0.028** |

**Supplementary Table 9**: Xenium Custom add-on panel

| Gene | Ensembl ID | Probesets | Codewords |
|---|---|---|---|
| EMCN | ENSG00000164035 | 4 | 4 |
| GPX3 | ENSG00000211445 | 3 | 3 |
| IGFBP7 | ENSG00000163453 | 3 | 3 |
| IGKC | ENSG00000211592 | 4 | 4 |
| SPP1 | ENSG00000118785 | 4 | 4 |
| ACKR2 | ENSG00000144648 | 5 | 5 |
| ACSM2B | ENSG00000066813 | 5 | 5 |
| APOD | ENSG00000189058 | 5 | 5 |
| APOE | ENSG00000130203 | 3 | 3 |
| AQP2 | ENSG00000167580 | 5 | 5 |
| ARNT2 | ENSG00000172379 | 5 | 5 |
| ATP6V0D2 | ENSG00000147614 | 5 | 5 |
| ATP6V1G3 | ENSG00000151418 | 5 | 5 |
| BMX | ENSG00000102010 | 5 | 5 |
| BST2 | ENSG00000130303 | 5 | 5 |
| C3 | ENSG00000125730 | 5 | 5 |
| CALD1 | ENSG00000122786 | 5 | 5 |
| CALM1 | ENSG00000198668 | 5 | 5 |
| CCL2 | ENSG00000108691 | 5 | 5 |
| CCL21 | ENSG00000137077 | 5 | 5 |
| CCL4 | ENSG00000275302 | 5 | 5 |
| CD300LG | ENSG00000161649 | 5 | 5 |
| CD69 | ENSG00000110848 | 5 | 5 |
| CD74 | ENSG00000019582 | 5 | 5 |
| CD9 | ENSG00000010278 | 5 | 5 |
| CHMP1A | ENSG00000131165 | 5 | 5 |
| CLDN11 | ENSG00000013297 | 5 | 5 |
| COL1A1 | ENSG00000108821 | 5 | 5 |
| COL1A2 | ENSG00000164692 | 5 | 5 |
| CXCL1 | ENSG00000163739 | 5 | 5 |
| CYP7B1 | ENSG00000172817 | 5 | 5 |
| DPEP1 | ENSG00000015413 | 4 | 4 |
| COL12A1 | ENSG00000111799 | 5 | 5 |
| FABP4 | ENSG00000170323 | 5 | 5 |
| FLT1 | ENSG00000102755 | 5 | 5 |
| NPHS1 | ENSG00000161270 | 5 | 5 |
| GADD45B | ENSG00000099860 | 5 | 5 |
| GATM | ENSG00000171766 | 5 | 5 |
| GLIS3 | ENSG00000107249 | 5 | 5 |
| GNG11 | ENSG00000127920 | 5 | 5 |
| FXYD4 | ENSG00000150201 | 5 | 5 |
| ID1 | ENSG00000125968 | 5 | 5 |
| IER3 | ENSG00000137331 | 5 | 5 |
| IGHG4 | ENSG00000211892 | 5 | 5 |
| IGLC2 | ENSG00000211677 | 4 | 4 |
| IL7R | ENSG00000168685 | 5 | 5 |
| ITGA8 | ENSG00000077943 | 5 | 5 |
| KCNIP4 | ENSG00000185774 | 5 | 5 |
| KLF6 | ENSG00000067082 | 5 | 5 |

| | | | |
|---|---|---|---|
| LCN2 | ENSG00000148346 | 5 | 5 |
| LTBP4 | ENSG00000090006 | 5 | 5 |
| LUM | ENSG00000139329 | 5 | 5 |
| LYVE1 | ENSG00000133800 | 5 | 5 |
| MDK | ENSG00000110492 | 5 | 5 |
| MIF | ENSG00000240972 | 3 | 3 |
| MIOX | ENSG00000100253 | 5 | 5 |
| MMP7 | ENSG00000137673 | 5 | 5 |
| NAMPT | ENSG00000105835 | 5 | 5 |
| NEAT1 | ENSG00000245532 | 5 | 5 |
| NFKBIA | ENSG00000100906 | 5 | 5 |
| NPHS2 | ENSG00000116218 | 5 | 5 |
| NR4A2 | ENSG00000153234 | 5 | 5 |
| ORAI1 | ENSG00000276045 | 4 | 4 |
| PLPP3 | ENSG00000162407 | 5 | 5 |
| PLSCR2 | ENSG00000163746 | 5 | 5 |
| PTPRB | ENSG00000127329 | 5 | 5 |
| PTPRC | ENSG00000081237 | 5 | 5 |
| RGS1 | ENSG00000090104 | 5 | 5 |
| RPL23 | ENSG00000125691 | 5 | 5 |
| PCLAF | ENSG00000166803 | 5 | 5 |
| SLC12A1 | ENSG00000074803 | 5 | 5 |
| SLC12A3 | ENSG00000070915 | 4 | 4 |
| SLC13A3 | ENSG00000158296 | 5 | 5 |
| SLC14A1 | ENSG00000141469 | 5 | 5 |
| SLC14A2 | ENSG00000132874 | 5 | 5 |
| SLC15A2 | ENSG00000163406 | 5 | 5 |
| SLC22A6 | ENSG00000197901 | 5 | 5 |
| SLC22A7 | ENSG00000137204 | 5 | 5 |
| SLC26A4 | ENSG00000091137 | 5 | 5 |
| SLC26A7 | ENSG00000147606 | 5 | 5 |
| SLC34A1 | ENSG00000131183 | 5 | 5 |
| ESCO2 | ENSG00000171320 | 5 | 5 |
| SLC4A4 | ENSG00000080493 | 5 | 5 |
| SLC4A9 | ENSG00000113073 | 5 | 5 |
| SLC5A12 | ENSG00000148942 | 5 | 5 |
| PRC1 | ENSG00000198901 | 5 | 5 |
| SLC6A19 | ENSG00000174358 | 5 | 5 |
| SLC7A13 | ENSG00000164893 | 5 | 5 |
| SLC8A1 | ENSG00000183023 | 5 | 5 |
| SOD2 | ENSG00000112096 | 5 | 5 |
| CRYAB | ENSG00000109846 | 5 | 5 |
| TAGLN | ENSG00000149591 | 5 | 5 |
| TFCP2L1 | ENSG00000115112 | 5 | 5 |
| VIM | ENSG00000026025 | 5 | 5 |
| XIST | ENSG00000229807 | 5 | 5 |
| YPEL2 | ENSG00000175155 | 5 | 5 |
| ZFP36 | ENSG00000128016 | 5 | 5 |
| SPP2 | ENSG00000072080 | 5 | 5 |
| SLC7A10 | ENSG00000130876 | 5 | 5 |
| SLC6A18 | ENSG00000164363 | 5 | 5 |
